# Learning induces persistent chromatin loops underlying robust gene expression during memory recall

**DOI:** 10.1101/2025.09.30.678146

**Authors:** Peibo Xu, Keerthivasan Raanin Chandradoss, Bradley Lukasak, Alekh Paranjapye, Abraham J. Waldman, Kenneth Pham, Han-Seul Ryu, Constin Liu, Katelyn R. Titus, Rahul Sureka, Jason Shepherd, Erica Korb, Jennifer E. Phillips-Cremins

## Abstract

Long-term memories are stored in neuronal ensembles called engrams, but the existence of persistent molecular traces in nuclei of engram neurons remain unknown. Using activity-dependent nuclear tagging *in vivo*, we profiled higher-order chromatin folding and DNA methylation in thousands of single hippocampal neurons up to a month after contextual fear conditioning (CFC). We find CFC-induced chromatin loop plasticity genome-wide, including persistently gained and lost loops with enduring traces *in vivo*. DNA methylation showed minimal CFC-induced persistence at promoters and enhancers. Persistently gained and lost loops connect distinct enhancers and promoters in excitatory and inhibitory subtypes and correlate with robust CFC-upregulated and -downregulated gene expression, respectively, upon recall. Synaptic genes associated with post-traumatic stress disorder and autism anchor neuronal subtype-specific persistent loops, suggesting relevance to neuropsychiatric dysfunction. We harness the power and sensitivity of multi-modal single-cell measurements to find enduring chromatin traces linked to robust gene expression during fear memory recall.

**Structured Abstract:** *Introduction:* Long-term memory is thought to be stored in specific neural circuits called engrams (*1*). Mechanisms centered at the synapse have been proposed, but what is altered within each engram cell that persists for the duration of the memory is unknown. Neural circuits fire action potentials in response to experiences. Such electrochemical signals are converted to molecular signaling pathways which travel from the synapse to the nucleus to activate new gene expression programs. However, experience-dependent RNA and protein molecules as well as synaptic plasticity phenomena are short-lived, lasting for only a few hours to several days (*2*). Thus, the extent to which chromatin and gene expression changes induced by behavior persist and functionally contribute to memory encoding, consolidation, and retrieval remains an important answered question. DNA is folded in the mammalian nucleus into higher-order long-range chromatin looping interactions (*3*). Loops form mechanistically through extrusion in which cohesin subunits form a ring that shuttles along chromatin and extrudes out the intervening DNA until it stalls at boundaries occupied by architectural proteins such as CTCF (*4, 5*). CTCF-independent looping mechanisms have also been uncovered (*6–10*). A subset of loops regulates gene expression by bringing distant enhancers into spatial proximity with their target promoters. Loops connect activity-dependent enhancers to their distal target genes to govern gene expression during *in vitro* neural stimulation and *in vivo* behavior paradigms (*11–14*). However, the extent to which chromatin changes persist on time scales to support long-term memory storage *in vivo* remains unclear.

*Rationale:* Multiple studies support the idea that higher-order chromatin architecture is plastic and can undergo activity- and experience-dependent remodeling linked to gene expression. Electron microscopy measurements provide direct evidence of chromatin architecture plasticity within minutes, with some changes persisting for up to an hour after potassium chloride stimulation of primary hippocampal neurons (*15*). Perturbing loops by selectively eliminating either *CTCF* or cohesin subunits impairs memory encoding in multiple behavior models (*11, 14, 16–18*), disrupts long-term potentiation (*17, 18*), and alters dendritic morphology (*13, 19, 20*). Cohesin-mediated loops are necessary for the establishment of new gene expression programs in post-mitotic neurons, including the upregulation of genes encoding axon guidance, dendritic spine morphology, and synaptic plasticity during neuron maturation *in vivo* as well as activity-dependent regulation of secondary response genes during neural stimulation *in vitro* (*12, 13*). The state of histone modifications before learning influences which neurons are recruited into memory traces, suggesting chromatin carries long-lasting yet adaptable information during memory encoding and long-term storage (*21*). Moreover, a subset of enhancers retains chromatin accessibility after learning *in vivo* (*11*). Together, these observations suggest that chromatin might provide a durable regulatory scaffold that persists long after the initial experience to govern gene expression programs required for the aspects of long-term memory storage.

*Results:* Here, we employ activity-dependent nuclear labeling in TRAP2 (targeted recombination in active populations) mice to isolate nuclei from neurons stimulated during contextual fear conditioning (CFC) (*22, 23*). Using single-nucleus methyl 3C-sequencing (snm3C-seq3), we simultaneously profiled DNA methylation and 3D genome folding in the same single tagged neurons from the hippocampus in a time course up to 28 days after CFC and during fear memory recall. Our multi-modal inquiry offered ability to group neurons by their neuronal subtype-specific DNA methylation profile, and thereby computationally generate pseudobulk Chromosome Conformation Capture heatmaps per each sorted hippocampal neuron subtype. We find CFC-induced chromatin loop plasticity genome-wide, including persistently gained and persistently lost loops with enduring structural traces up to 28 days after training. Persistent loops connect distinct distal enhancers to target genes encoding synaptic plasticity and neurotransmitter signaling pathways unique to each excitatory and inhibitory neuron cell type. By contrast to loops, we observe negligible persistence of CFC-induced changes in DNA methylation at enhancers and promoters. Persistently gained and lost loops show upregulation and downregulation of gene expression upon recall, respectively, with the functional effect focused on genes with mRNA levels influenced during training. We uncovered a strong enrichment for genes associated with post-traumatic stress disorder and autism spectrum disorder anchoring the base of persistent loops, suggesting the relevance of enduring chromatin architecture traces to dysregulation of fear, anxiety, and stress in neuropsychiatric disorders.

*Conclusion:* Our findings reveal that learning induces neuronal subtype-specific changes in chromatin loops, a subset of which remain durable for at least a month *in vivo*. Persistent chromatin loops are linked to expression of disease-associated synaptic genes during fear memory recall, thus highlighting relevance to neuropsychiatric and neurodevelopmental disorders. Our work opens up new avenues for investigating enduring chromatin traces for their role in governing cell-wide mRNA levels of genes that impact synaptic plasticity during fear learning and long-term memory storage.

## Main Text

Encoding and storage of memory involves plasticity in the brain at tissue, circuit, synapse, and nuclear scales. A leading theory is that long-term memory is stored in distributed neuronal circuits termed engrams (*1*), but what is altered within each individual neuron that persists for the duration of the memory is unknown. Structural and functional synaptic plasticity occurs on rapid time scales of seconds to minutes and individual RNA and protein molecules exhibit short half-lives of hours to days. Therefore, the identification of durable features in the brain that can persist after the original stimulus remain an important unanswered question (*2*).

The extent to which experience-induced chromatin and gene expression changes persist over long time scales and functionally contribute to memory storage and retrieval in engram cells is unknown. Loops connecting activity-dependent enhancers to distal genes are induced during neural stimulation *in vitro* and behavior paradigms *in vivo* (*11–14*). Genetic elimination of the architectural proteins CTCF or cohesin prior to learning substantially impairs memory encoding in multiple behavior tasks *in vivo* (*11, 14, 16–18*). Cohesin-mediated loops are necessary for the establishment of new gene expression programs in post-mitotic neurons, including the upregulation of synaptic genes during neuron maturation *in vivo* and activity-dependent secondary response genes during neural stimulation *in vitro* (*12, 13*). Fear conditioning induces genome-wide chromatin accessibility changes at enhancers, and such changes persist at least five days after fear learning (*11*). Moreover, a recent work shows that a neuron’s pre-learning chromatin state biases its recruitment into memory ensembles, and experimental manipulation of chromatin plasticity increases a cell’s allocation to the learning ensemble (*21*). Together, these data suggest that understanding activity-dependent loops *in vivo*, the extent of their persistence, and the functional effect of loop persistence would advance our understanding of an important scale of memory encoding and storage.

Testing for persistent chromatin loops has been technically challenging because memories are encoded by sparse populations of neurons (*24–28*). As a result, bulk genomic approaches in whole brain tissue cannot resolve the subtle, cell type-specific epigenetic variations that may define memory ensembles (*11, 24, 26–30*). We employed contextual fear conditioning (CFC) by exposing mice to a novel context followed by an aversive foot shock (*31*). CFC is a rigorous behavioral model suited for finding persistent chromatin traces because one training event elicits a strong freeze response and context re-exposure evokes a fear memory (*32*). We use multi-modal single-nucleus methyl 3C-sequencing (snm3C-seq3) (*33, 34*) to profile DNA methylation and chromatin architectural changes in the same single-neurons sorted from the hippocampus and search for chromatin patterns active during fear memory encoding, persistent for up to 28 days, and linked to gene expression during memory recall. To overcome the sparsity of single cell Chromosome Conformation Capture data, we analyze DNA methylation genome-wide, cluster neurons by similar signatures, create pseudobulk genome folding maps in neurons matched by DNA methylation profile, and identify loops with computational tools developed by our group. We identified specific promoter-enhancer chromatin loops in distinct neuron subtypes that arise shortly after memory encoding and persist for at least four weeks *in vivo* on time scales relevant for memory storage.

## Results

### Longitudinal multi-modal profiling of higher-order genome folding and DNA methylation in single-neurons after contextual fear conditioning in vivo

CFC engages 5-20% of excitatory neurons in the hippocampus during fear memory encoding (*31, 35–37*). To genetically access neurons activated during CFC, we bred the homozygous Targeted Recombination of Active Populations (TRAP2) mouse with a homozygous Sun1-GFP nuclear reporter mouse (*22, 23, 38*). TRAP2 is engineered with a transgene encoding a tamoxifen-inducible CRE recombinase (iCreERT2) knocked in-frame to *Fos*. Upon circuit activation during CFC and tamoxifen injection, CreERT2 binds to externally administered tamoxifen, translocates to the nucleus, and removes a stop codon within the pCAG-Sun1sfGFP transgene for permanent fluorophore expression within the stimulated neuron nucleus **(Fig. 1A, fig. S1A)**.

**Fig. 1.**
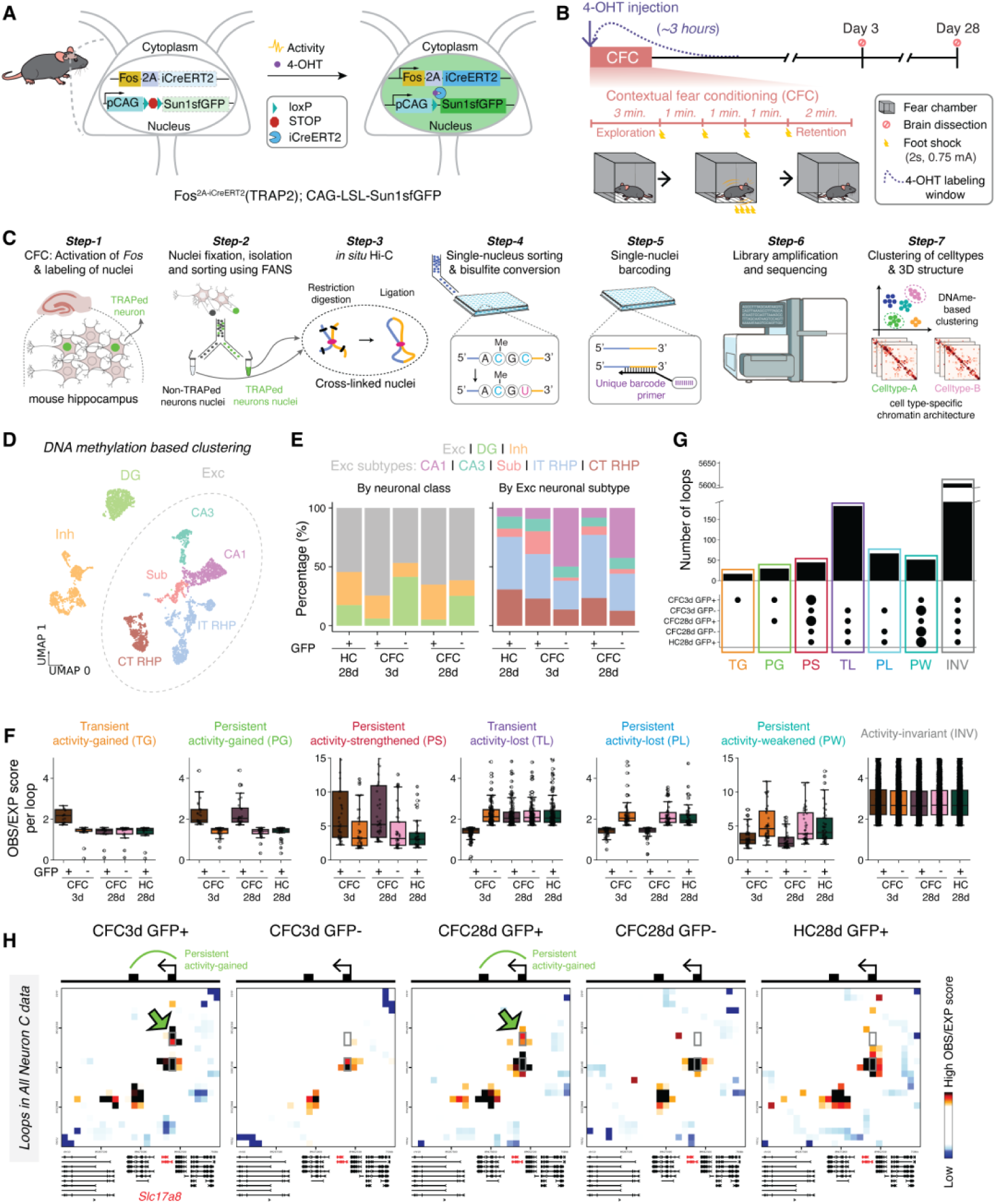
Persistent activity-dependent chromatin loop plasticity for up to a month after neuronal activation in contextual fear conditioning. (A-B) Schematic of TRAP2; Sun1sfGFP mice. Mice are injected with 4-OHT and subjected to contextual fear conditioning leading to a rapid upregulation of Fos and iCreERT2. Injection of 4-OHT during the training window enables iCreERT2 to translocate to the nucleus where it removes the stop cassette, resulting in the permanent expression of Sun1-sfGFP and nuclear labeling of neurons active during the approximately 3-hour 4-OHT administration window. During CFC, mice explored the fear chamber for 3 minutes followed by four 2-second foot shocks (0.75 mA) delivered at 1-minute intervals, with a 2-minute retention period. Hippocampal brain tissue was collected at two timepoints: day 3 and day 28 post-CFC. (C) Schematic of snm3C-seq3 multi-modal measurements of DNA methylation and higher-order chromatin folding in TRAPed+ and TRAPed-neurons. (D) Uniform manifold approximation and projection (UMAP) visualization of hippocampal NeuN+ cells (n = 4,667 cells from 2 male mice/condition) showing distinct clusters including excitatory (Exc), inhibitory (Inh), and DG neurons. Exc further includes CA1, CA3, subiculum (Sub), corticothalamic (CT), and intratelencephalic (IT) retrohippocampal region (RHP) neuronal subtypes. (E) Quantification of major neuronal class composition (left) and excitatory neuronal subtypes composition (right) across behavioral conditions. (F) Boxplots show the Observed/Expected (OBS/EXP) interaction frequency for chromatin loop classes across CFC3d GFP+/-, CFC28d GFP+/-, and HC28d GFP+ conditions. (G) Upset plot shows the number of loops in 7 loop classes. (H) Interaction frequency heatmaps (OBS/EXP) around the *Slc17a8* gene across 5 conditions with schematics representing the rewiring of loops from promoter to distal non-coding enhancers (black squares) on the top of the heatmaps. Grey box, location of loop called. Green arrow, persistent activity-gained loop.

To label neurons activated by CFC, we administered 4-hydroxytamoxifen (4-OHT) to TRAP2/Sun1sfGFP mice prior to training. Both male and female mice underwent classic fear conditioning and exhibited the expected freeze response (**Materials and Methods, Fig. 1B, fig. S1B-D, Table S2**). Control mice received 4-OHT alone and remained in their home cages (HC) for 28 days. We micro-dissected hippocampal and retrohippocampal regions from CFC and HC mice (n = 2 mice per group, **fig. S2**) and used fluorescence-activated nuclei sorting (FANS) to isolate TRAPed+ (GFP+) and TRAPed- (GFP-) neuronal nuclei at recent (3-day) and remote (28-day) memory timepoints (*39*). Immunohistochemistry and FANS results confirm that TRAPed+ and TRAPed-populations are present in tissue and can be cleanly sorted from HC 28 days, CFC 3 days, and CFC 28 days conditions (**fig. S1E-F, fig. S2**).

We used snm3C-seq3 to jointly profile DNA methylation and genome folding at loop resolution in 4,667 hippocampal single neurons across all experimental conditions (**Fig. 1C**). We sequenced to a depth of on average of ∼2 million paired end reads per cell and achieved an average of ∼1 million usable reads per cell after systematic filtering per methods previously reported (**fig. S3A-M, Materials and Methods, Tables S1, S3-S4**). Using established published methods (*34, 40*), we normalized DNA methylation data at 100 kilobase bin resolution for equal mean sequencing depth across neurons (**figs. S4-S5**) and completed dimensionality reduction via principal component analysis (**figs. S6-S8)**. We identified distinct neuronal subtypes by clustering on genome-wide DNA methylation and integration with subtype specific markers established by the Allen Brain Atlas (*41*) (**figs. S9-S11**, **Materials and Methods**).

Our analysis of HC TRAPed+, CFC TRAPed+, and CFC TRAPed- at 3 and 28 days post-CFC revealed seven neuronal subtypes across the hippocampal formation: five excitatory populations (*cornu ammonis* 1 (CA1), CA3, subiculum, intratelencephalic (IT RHP), and corticothalamic (CT RHP)), one inhibitory population, and dentate gyrus (DG) neurons, consistent with the anatomical sections with the hippocampal formation (**Fig. 1D, figs. S11-S12**). TRAPed+ neurons were detected in both hippocampal and retrohippocampal regions **(fig. S12)**, and in excitatory, inhibitory, and DG neurons (**Fig. 1E, fig. S12**), consistent with the literature and validated with immunohistochemistry analysis of GFP+ neurons (*42–44*) **(fig. S1E-F, Tables S5-S7)**. Altogether, we produce high sequencing depth, high resolution snm3C-seq3 data for multi-modal DNA methylation and higher-order genome folding analysis in more than 4,700 hippocampal neurons in a time course of CFC, both those trapped as active during learning and those that were not trapped as internal controls.

### CFC-induced chromatin loop plasticity persists for up to a month in TRAPed+ fear memory neuronal ensembles

Single-cell 3C-based methods produce sparse matrices that lack the complexity for sensitive and specific detection of chromatin loops. To overcome this substantial technical barrier, we developed a computational pipeline to call loops genome-wide after pooling neurons with matched DNA methylation profiles into pseudobulk Chromosome Conformation Capture interaction frequency matrices (**fig. S13, Materials and Methods**).

We adapted our previously published methods to scale single-neuron counts matrices by distance to correct for cell-to-cell variation in sequencing depth, generated pseudobulk matrices matched by subtype-specific DNA methylation signatures, and Knight-Ruiz matrix balanced the pseudobulk data to correct for bin-to-bin read mapping efficiency differences (**fig. S13, Materials and Methods**) (*6, 9, 12, 13, 45–52*). We applied imputation with a Gaussian smoothing window to improve signal-to-noise and adapted our published methodologies to use the donut geometrical window to compute a local expected interaction frequency (*6, 9, 12, 13, 45–52*) (**Materials and Methods**). We called loops as those pixels exhibiting a significantly higher Observed compared to Expected interaction frequency **(figs. S13-S14, Table S5**). Heatmaps representing Observed/Expected interaction frequency were of high signal to noise ratio, dot-like structures indicative of loops being visually apparent, and maps were highly similar between biological replicates of neurons derived from different animals (**fig. S15**). Our data confirm the generation of loop-resolution heatmaps with snm3C-seq3 from small numbers of neuronal nuclei derived from tissue *in vivo*.

We next set out to classify loops by the extent to which they were induced or lost by CFC and exhibited persistent signal between 3- and 28-days *in vivo*, using HC control (HC28d TRAPed+ neurons) as the activity-dependence threshold **(Fig. 1F-G**, **Materials and Methods)**. We employed a thresholding strategy to identify seven classes of loops in pseudobulk heatmaps with all cell types pooled together: (1) transiently activity-gained (TG) loops present in TRAPed+ versus TRAPed-neurons at 3 days but absent in both TRAPed+ and TRAPed-neurons at 28 days post-CFC, as well as in HC control, (2) persistently activity-gained (PG) loops present in TRAPed+ versus TRAPed-neurons at 3 days and still present in TRAPed+ neurons at 28 days post-CFC but absent in HC control, (3) persistently activity-strengthened (PS) loops present in both TRAPed+ and TRAPed-neurons at 3-and 28-days post-CFC, but further strengthened in TRAPed+ neurons at 3- and 28-days (with more than 1.25-fold higher interaction frequency compared to TRAPed-neurons at 3 and 28-days and compared to HC control), (4) transiently activity-lost (TL) loops lost in TRAPed+ versus TRAPed-neurons at 3 days but gained loop in both cell types at 28 days post-CFC and HC control, (5) persistently activity-lost (PL) loops lost in TRAPed+ versus TRAPed-neurons at 3 days and still lost in TRAPed+ versus TRAPed-neurons at 28 days post-CFC, but present in HC control, (6) persistently activity-weakened (PW) loops present in both TRAPed+ and TRAPed-neurons at 3- and 28-days post-CFC and further weakened in interaction frequency in TRAPed+ versus TRAPed-neurons (with more than 1.25-fold lower interaction frequency compared to TRAPed-neurons at 3- and 28-days post-CFC and compared to HC control), and (7) invariant loops exhibiting strong interaction frequency across all conditions (**Fig. 1F-G, Table S9**). When using pseudobulk heatmaps relying on the pooling of all neuronal subtypes together, most loops remained invariant (INV) across conditions (n = 5604) (**Table S9**). Nevertheless, our data demonstrate proof of concept for stratifying loops by their structural plasticity across a longitudinal time course of CFC in TRAPed neurons.

We note an example of a persistent activity-gained loop at the *Slc17a8* locus, which encodes the vesicular glutamate transporter protein vGlut3 and is enriched in a subset of hippocampal neurons (*53*). In TRAPed+ neurons (CFC3d GFP+ and CFC28d GFP+), we observed long-range (∼300 kb) interactions connecting the *Slc17a8* promoter to an upstream putative enhancer, which were absent in TRAPed-neurons (CFC3d GFP- and CFC28d GFP-) and the HC control (HC28d GFP+) **(Fig. 1H)**. Together, these findings demonstrate that CFC-activated neurons harbor persistent, CFC-induced chromatin loops, in addition to transient loop plasticity that resumes the original structural state once the stimulus is removed.

### CFC-induced persistent loops are distinct between excitatory and inhibitory neurons

Given DNA methylation profiles genome-wide are neuronal subtype-specific, we sought to understand the extent to which CFC-induced or -lost persistent chromatin loops exhibit cell type-specific features. We generated pseudobulk heatmaps from pooled excitatory neurons only and matched equal numbers from each subtype across all conditions (43 per subtype: CA1, CA3, CT RHP and IT RHP, 172 total per pseudobulk, **Table S6, Table S9**). We stratified loop classes as in Figure 1 and identified N=100 persistently gained loops and N=123 persistently lost loops in TRAPed+ excitatory neurons across both 3- and 28-days post-CFC **(Fig. 2A, fig. S16, Materials and Methods).** We provide an example of an excitatory neuron-specific, persistently gained loop between a putative activity-dependent enhancer and the *Serpinh1* promoter **(Fig. 2B)** which encodes a molecular chaperone for gamma-aminobutyric acid type A (GABA-A) receptors (*54*). Thus, persistently gained loops can be detected in TRAPed+ excitatory neurons at 3- and 28-days post-CFC (Exc-CFC3d GFP+ and Exc-CFC28d GFP+) but absent in TRAPed-counterparts.

**Fig. 2.**
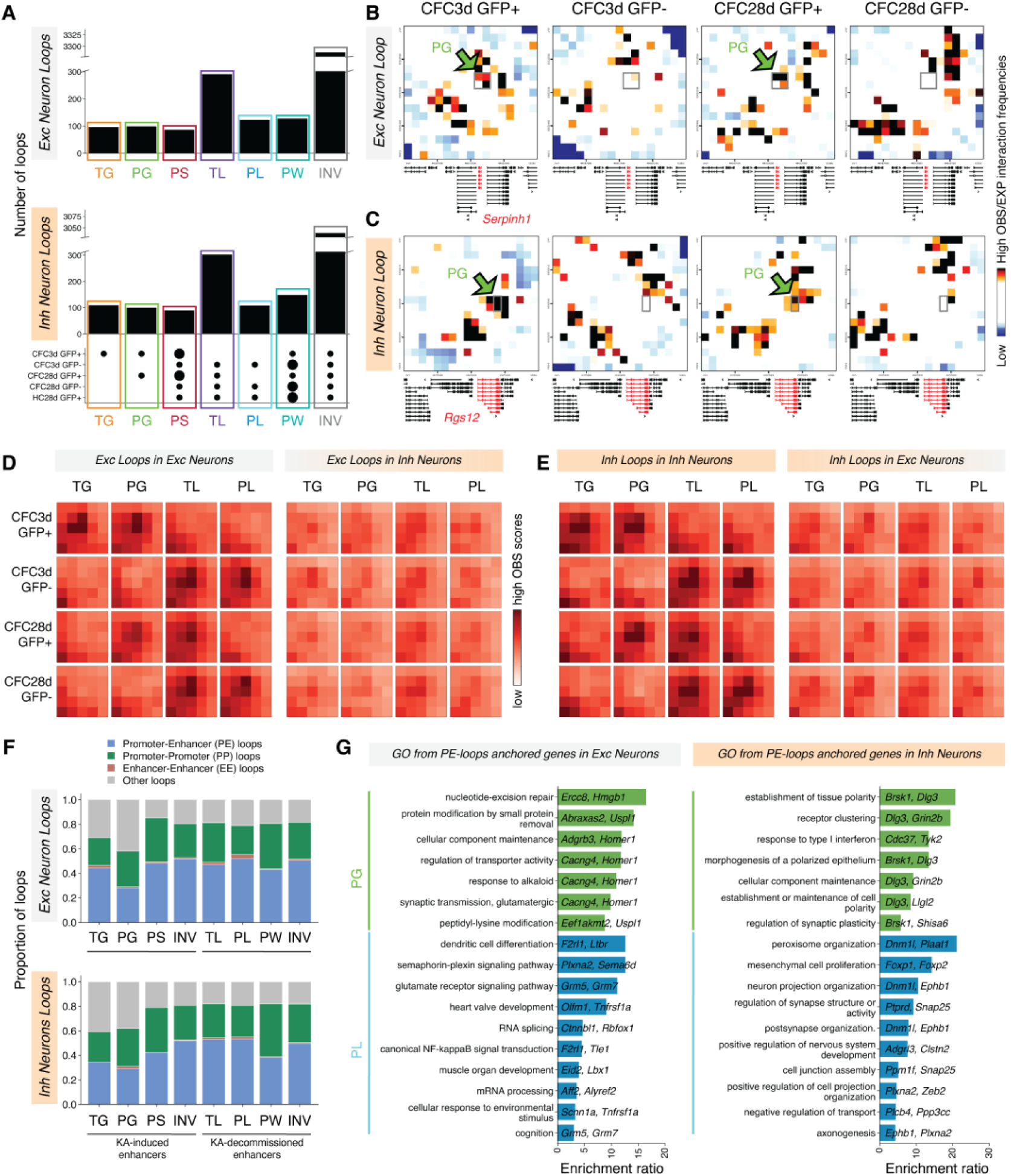
Persistent chromatin loops are unique to excitatory and inhibitory neurons and anchor distinct synaptic plasticity genes. **(A)** Upset plot showing excitatory (top) and inhibitory (bottom) loop numbers across 5 conditions (CFC3d GFP+/-, CFC28d GFP+/-, HC28d GFP+) for persistently activity-gained, PG; transiently activity-gained, TG; persistently activity-strengthened, PS; persistently activity-lost, PL; persistently activity-weakened, PW; transiently activity-lost, TL; and invariant loops. (**B, C**) Heatmaps showing interaction frequency (OBS/EXP) at **(B)** the *Serpinh1* gene in excitatory neurons or **(C)** at the *Rgs12* gene in inhibitory neurons across CFC3d GFP+/- and CFC28d GFP+/- conditions. **(D-E)** APA plots of interaction frequency for (**D)** loops called in excitatory neurons using snm3C-seq data from excitatory and inhibitory neurons and **(E)** loops called in inhibitory neurons using snm3C-seq data from excitatory and inhibitory neurons. **(F)** Stacked bar plot showing the distribution of P-P, P-E, and E-E loop categories for excitatory (top) and inhibitory (bottom) neurons across loop classes listed in (A). **(G)** Ontology of the highest enriched genes anchoring persistent activity-gained (PG) and persistent activity-lost (PL) loops in excitatory (left) and inhibitory neurons (right).

Inhibitory neuron electrophysiology is altered in fear memory (*55, 56*). We identified N=100 persistently gained loops and N=110 persistently lost loops in TRAPed+ inhibitory neurons across both 3- and 28-days post-CFC (**Fig. 2A-C, fig. S16, Tables S9-S10**). To compare chromatin reorganization between inhibitory and excitatory neurons after CFC, we performed aggregate peak analysis (APA) to quantify the strength of chromatin interactions at loop anchors. We found that persistently and transiently gained and lost CFC-induced loops identified in inhibitory neurons were absent in excitatory neurons and vice versa (**Fig. 2D-E**). Together, these data demonstrate unique patterns of inhibitory and excitatory neuron-specific chromatin architecture during memory storage.

### Persistent promoter-enhancer chromatin loops anchor genes linked to unique synaptic signaling pathways in excitatory and inhibitory neurons

Chromatin loops can regulate gene expression by physically co-localizing distal enhancers with their target genes (*57*). To approximate the genomic location of putative activity-gained and activity-lost enhancers, we re-analyzed ChIP-seq data for the histone modification H3K27ac generated in samples derived from a paradigm of pharmacologically induced stimulation of hippocampal CA1 neurons (*58*). We identified non-coding putative cis-regulatory elements genome-wide as intergenic or intronic activity-gained or activity-lost H3K27ac peaks and stratified loops into Promoter-Promoter (P-P), Promoter-Enhancer (P-E), Enhancer-Enhancer (E-E), and other loops encompassing all remaining genomic interactions (**Materials and Methods**) (**Fig. 2F**). To determine the biological relevance of the genes regulated by P-E loops, we next assessed the ontology of loop-anchored genes. We found that persistently gained promoter-enhancer loops anchored genes involved in synaptic transmission (*e.g.*, *Homer1*, *Cacng4*) and DNA repair (*e.g.*, *Ercc8*, *Hmgb1*) in excitatory neurons and genes associated with synaptic receptors (*e.g.*, *Dlg3*, *Grin2b*) and regulation of synaptic plasticity (*e.g.*, *Brsk1, Shisa6*) in inhibitory neurons (**Fig. 2G, Table S11-S12**). Moreover, another form of durable promoter-enhancer loop, those that are persistently strengthened at 3 days post-CFC and persist in TRAPed+ neurons at 28 days post-CFC, anchored genes involved in neurotrophic signaling pathways, transmembrane transport, and synaptic transmission (*e.g.*, *Bdnf, Magi2, Slc38a1, Trpm1, Cacnb3, Kcne2*) in excitatory neurons and genes associated with mitochondria depolarization, GABA signaling pathways, and circadian rhythms (*e.g.*, *Huwe1, Vdac1, Gabra4, Gabrb1, Ptger4*) in inhibitory neurons (**fig. S17, Tables S11-12**). Our data demonstrate genes of significant relevance of genes anchored by CFC-induced persistently gained and persistently strengthened loops to synaptic signaling pathways and general synaptic molecular processes unique to excitatory and inhibitory neurons.

Another significant phenomenon in our data is activity-lost enhancers anchoring persistently lost loops that remain lost at 28 days post-CFC. We examined the genes anchoring persistently lost promoter-enhancer loops (**Fig. 2G, Tables S11-12**). In excitatory neurons, the unique persistently lost promoter-enhancer loops anchored genes associated with metabotropic glutamate receptors, semaphoring-plexin signaling pathways, RNA splicing, and cellular response to environmental stimuli (*e.g.*, *Grm5, Grm7*, *Plxna2*, *Sema6d, Rbfox1, Ctnnbl1, Scnn1a, Tnfrsf1a*). Moreover, in inhibitory neurons, the anchored genes were linked to neural projections, synaptic structure, ion transport, synaptic signaling, and axon outgrown (*e.g.*, *Dnm1l, Ephb1, Ptprd*, *Snap25, Ppm1f, Plxna2, Zeb2, Plcb4*) (**Fig. 2G**). Collectively, our findings establish that CFC-induced persistently gained and persistently lost loops in either TRAPed+ excitatory or inhibitory neurons are not merely structural but also couple activity-dependent enhancers to distal target genes central to synaptic plasticity.

### Negligible persistence of CFC-induced DNA methylation changes at enhancers and promoters

Locus-specific changes in DNA methylation at regulatory elements such as promoters and enhancers has been reported in a range of model systems (*5, 59, 60*) and activity-dependent changes in DNA methylation have also been reported in mammalian neurons (*24, 26*). How DNA methylation relates to looped enhancers and promoters generally, and specifically in the context of fear conditioning *in vivo* is unknown.

Given the subtype specific patterns of DNA methylation genome-wide, we projected that loops called in individual excitatory subtypes would be essential for the integration with DNA methylation phenomenon. We identified loops according to the established classes in four major excitatory subtypes: CA1, CA3, CT RHP, and IT RHP, as well as DG and inhibitory neurons (**Materials and Methods, fig. S18, Table S7, Tables S9-S10**). To confirm the specificity of subtype-specific loops, we performed APA analyses across excitatory subtypes and neuronal classes. Subtype-specific loops showed robust APA signals within their originating subtype but weak or absent signals in DG or inhibitory neurons (**Fig. 3A-B, figs. S18-S21**). Our results confirm that persistently gained and lost loops in CFC are unique not only in comparing excitatory to inhibitory neurons, but also among excitatory neurons derived from different regions of the hippocampus. The results highlight the importance of single-cell multi-modal methods to evaluate chromatin in the brain and suggest that bulk analyses may miss important observations conflated in an amalgam of subtype-specific events.

**Fig. 3.**
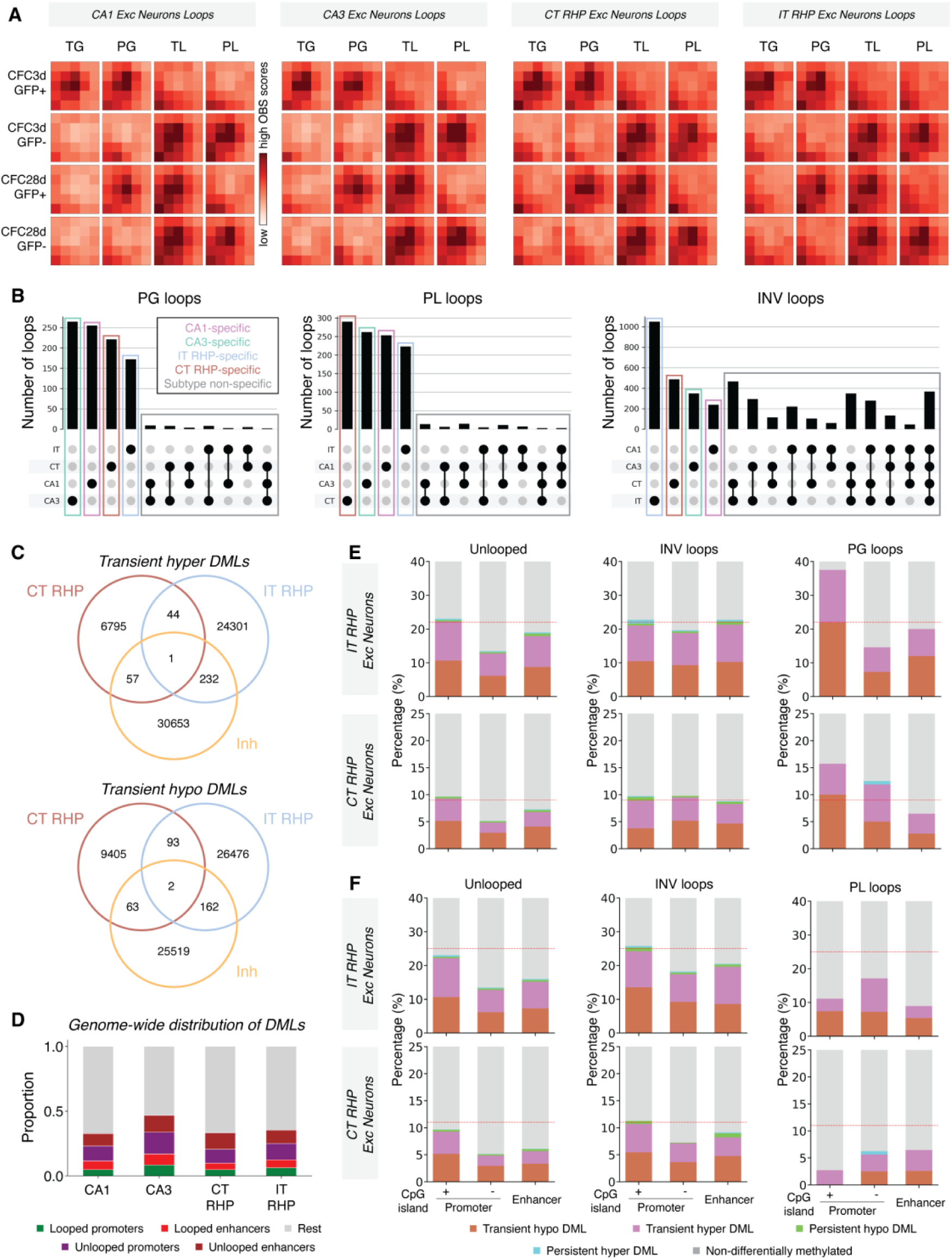
Persistent chromatin loops are excitatory subtype-specific and minimal persistence of DNA methylation at promoters and enhancers genome-wide is observed. (**A**) APA plots of subtype-specific loops in excitatory subtypes. (**B**) Upset plots showing the overlap of persistently activity-gained (PG), persistently activity-lost (PL), and invariant loops identified in each excitatory subtype. (**C**) Venn diagram showing the overlap of transient hyper and hypo DMLs identified in IT RHP, CT RHP, and Inh neuron type. (**D**) Stacked bar plots show the relative distribution of persistent DMLs overlapping looped promoters, and looped enhancers, as well as unlooped promoters, unlooped enhancers, and remaining genomic regions (Rest) in CA1, CA3, CT RHP, and IT RHP neurons. (**E, F**) Distribution of DMLs across CpG island promoters, non-CpG island promoters, and enhancers. Stacked bar plots show the relative proportions of transient hyper, transient hypo, persistent hyper, persistent hypo DMLs, and non-differentially methylated sites within each loop class. Y-axes are set to a maximum of 40 (IT) and 25 (CT) to highlight detail. For enhancer analysis, KA-induced enhancers were used in (E), and KA-decommissioned enhancers were used in (F).

We conducted a rigorous assessment of genome-wide locations of differentially methylated loci (DMLs) at CpG resolution between TRAPed+ and TRAPed-conditions at 3- and 28-days post-CFC in the individual neuronal subtypes (**Materials and Methods, Fig. 3C-F, figs. S22-26, Table S13**). Although sensitivity is limited by cell numbers, we could still use rigorous thresholds and published gold-standard software to identify hyper- and hypomethylated DMLs (Materials and Methods). DMLs changing in TRAPed+ versus TRAPed-conditions were largely neuron subtype-specific when detectable (**Fig. 3C, Table S13, fig. S26**). An examination of persistent DMLs genome-wide revealed that 30-50% were co-localized with enhancers and promoters with no bias toward whether or not they were looped (**Fig. 3D**). The majority of DMLs were not localized with enhancers or promoters.

It is particularly noteworthy that across all neuronal subtypes, we observed negligible overlap of CFC-induced DMLs that remained persistent through 28 days at promoters and enhancers (**Fig. 3E**). We observed a baseline level of either transient hyper- or hypo-DMLs unique to only the 3 days post-CFC timepoint or the 28 days post-CFC timepoint at promoters or enhancers independent of their loop status. Moreover, we observed minimal to no persistent DMLs at any promoter or enhancer element, whether in transient loops, persistent loops, or unlooped. Our data suggest that DMLs induced by CFC are predominantly confined to the time point of 3 days or 28 days post-CFC when localized to promoters and enhancers, suggesting that DML persistence is not a predominant feature of regulatory elements. Our data are consistent with a model in which there is negligible DML persistence at persistently gained or persistently lost enhancer-promoter loops.

### Persistently lost promoter-enhancer loops anchor downregulated genes during memory recall

To determine the link between loop persistence and gene expression, we designed a memory recall experiment (**Fig. 4A**). We re-exposed mice to the same context at three days post-CFC, but without foot shock delivery (**Fig. 4A)**. Exposure to the context alone without foot shock elicited robust memory recall, as evidenced by significantly increased freezing time compared to pre-stimulus (**figs. S27**). We sorted TRAPed+ and TRAPed-neuronal nuclei three hours post-recall (CFC3d RC) and compared to neurons from the no-recall CFC3d group for single-nucleus profiling of gene expression using Smart-seq3 (61) (**fig. S28**).

**Fig. 4.**
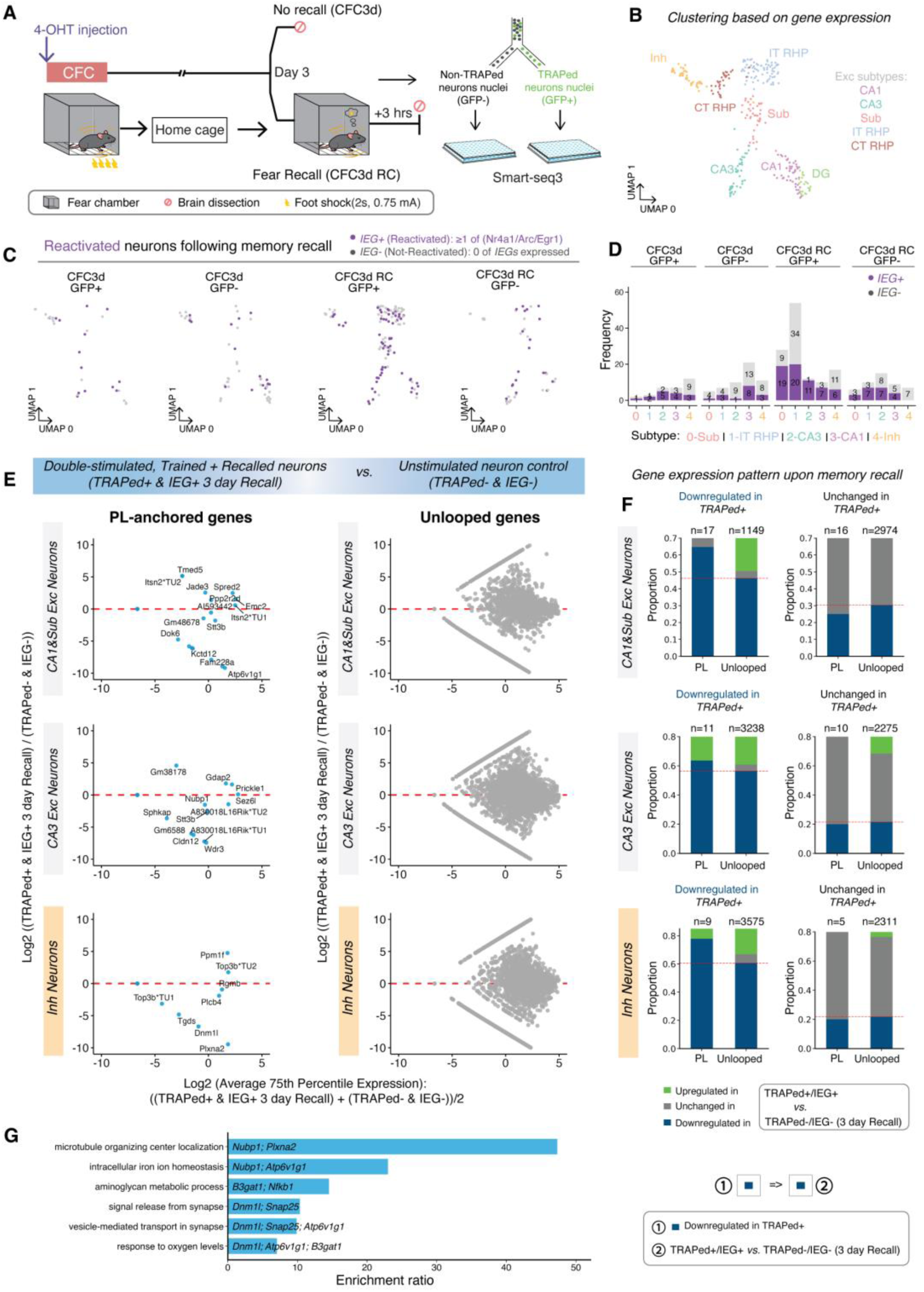
Persistently lost promoter-enhancer loops are associated with robust and selective downregulation of CFC-repressed genes during memory recall. (**A**) Experimental design for contextual fear memory recall. CFC-trained mice were re-exposed to context alone after 3 days to trigger memory recall. Brains were harvested 3 hours after recall for single-cell profiling of gene expression using Smart-seq3. (**B**) UMAP visualization of hippocampal neuron populations based on gene expression profiles from Smart-seq3 across CFC3d and CFC3d Recall groups. (**C-D**) UMAP visualization and quantification of neurons classified as upregulated during recall by a panel of IEG expression across neuronal subtypes and experimental conditions. (**E**) M-A plots comparing double-stimulated neurons (TRAPed+IEG+ at 3 day recall) to unstimulated neuron control (TRAPed-IEG- at 3 day recall). Left, persistently lost promoter-enhancer loop-anchored genes; right, unlooped genes. X-axis: log2 average 75th-percentile across the two conditions; Y-axis: log2 fold change between conditions. (**F**) Proportion of genes upregulated (green), downregulated (blue), or unchanged (grey) upon recall in TRAPed+IEG+ versus TRAPed-IEG-neurons. PL, genes anchoring persistently lost promoter-enhancer loops; Unlooped, genes not engaged in looping interactions. Column 1 represents genes downregulated at 3 days post-CFC in TRAPed+ vs. TRAPed-neurons. Column 2 represents genes unchanged at 3 days post-CFC in TRAPed+ vs. TRAPed-neurons. (**G**) Gene ontology for persistently lost loop-anchored genes that were downregulated in TRAPed+ vs. TRAPed-neurons at 3 days post-CFC and further downregulated in TRAPed+IEG+ vs. TRAPed-IEG-neurons upon memory recall.

After rigorous quality control steps to ensure high-quality and rigorous analyses, we performed clustering based on mRNA levels using established methods (**Materials and methods, fig. S29-30**) (*61*). Consistent with our snm3C-seq3, we identified three major neuronal cell types from our tissue, including excitatory (*Slc17a7*+), inhibitory (*Gad2*+), and DG (*Prox1*+), as well as further subdivided excitatory subtypes **(Fig. 4B; figs. S29-S31)**. We note a strong independent validation of our neuronal subtype proportions given clustering based on snm3C-seq3 and on Smart-seq3 data show similar results within each biological condition (**figs. S31**). Moreover, we see strong concordance of neuronal subtype proportions within each biological condition across mouse replicates (**figs. S31**), thus further highlighting the reproducibility of our data and analyses between independent animal replicates.

To distinguish reactivated from non-reactivated neurons during memory recall, we classified cells as immediate early gene-positive (IEG+) if they expressed at least one of the three established IEGs *Nr4a1*, *Arc*, or *Egr1* (**Fig. 4C-D**). The three IEGs were selected due to their essential role in hippocampus-dependent contextual memory formation (*62–64*). We did not use *Fos* as a reactivation marker because the sequencing depth and gene coverage limits of Smart-seq was insufficient for reliable detection of *Fos* mRNA. As expected, the recall TRAPed+ group showed higher enrichment of reactivated (IEG+) cells compared to TRAPed-recall, TRAPed+ no-recall, and TRAPed-no-recall groups (**Fig. 4C-D**). In our data, neurons expressing IEGs in recall were predominantly excitatory, with fewer cells from inhibitory populations. Thus, we can identify TRAPed+IEG+ neurons active during both learning and in recall and TRAPed-IEG-neurons inactive in both learning and recall in multiple excitatory subtypes for downstream integration with loops.

Since promoter-enhancer loops facilitate gene expression by bringing distal regulatory elements into spatial proximity with target promoters (*57*), we reasoned that the dissolution of contacts in persistently lost loops would result in altered expression of anchored genes. We first compared mRNA levels in TRAPed+ to TRAPed-neurons at 3 days post-CFC in the absence of recall stimulation (**fig. 32A**) or those TRAPed+IEG- to TRAPed-IEG-neurons not stimulated during recall (**fig. 32B**). We identified genes from those two baseline comparisons that were downregulated or unchanged in expression by the initial CFC training at 3 days (**fig. 32C**). We next carried forward the CFC-downregulated and CFC-unchanged gene sets from 3 days post-training and compared gene expression in these two sets for changes between TRAPed+IEG+ and TRAPed-IEG-neurons upon recall (**Fig. 4E-F).** We observed an enrichment for downregulation of gene expression upon recall for genes anchoring persistently lost promoter-enhancer loops compared to unlooped controls (**Fig. 4E-F, Table S14**). It is particularly noteworthy that the downregulation during recall was enriched at persistently lost loops only at the subset of genes that were also downregulated during training due to CFC. We also note that we used activity-lost enhancers identified in CA1 tissue, which can explain the strongest enrichment for downregulated genes at persistently lost promoter-enhancer loops in CA1/Sub. Ontology analysis revealed that genes downregulated upon recall at persistently lost loops encode microtubule organizing center localization and vesicle-mediated synaptic transport (*e.g.*, *Dnm1l*, *Snap25, Atp6v1g1, Nubp1, Plxna2*; **Fig. 4G, Table S14**). Our data reveal a link between downregulation of synaptic genes during memory recall and persistently lost promoter-enhancer loops and highlight the potential relevance of persistent loop loss in the repression of synaptic signaling pathways that are meant to be permanently repressed during memory.

### Persistent activity-gained promoter-enhancer loops enable neuronal subtype-specific upregulation of synaptic genes during memory recall

We also sought to understand the relationship between persistently gained promoter-enhancer loops and gene expression. Using the same analyses paradigm as described (**Materials and Methods**), we compared mRNA levels in TRAPed+ to TRAPed-neurons at 3 days post-CFC in the absence of recall stimulation to identify genes that were upregulated or unchanged in expression by the initial CFC training at 3 days (**fig. S33**). We carried forward the CFC-upregulated and CFC-unchanged gene sets and compared gene expression in these two sets for changes between TRAPed+IEG+ and TRAPed-IEG-neurons upon recall (**Fig. 5).** We found a robust enrichment for upregulated gene expression upon recall for genes anchoring persistently gained promoter-enhancer loops compared to transiently gained loops and unlooped controls (**Fig. 5A-B, Table S15**). By contrast, genes unchanged during CFC at 3 days post-training did not respond to recall when anchored at persistent loops or unlooped. Ontology analysis showed that genes robustly upregulated at persistently gained loops encode DNA repair and chromatin remodeling pathways, synaptic receptors, and mitochondria functions (e.g., *Usp20*, *Brcc3*, *Fam168a, Shisa6, Actr10*) (**Fig. 5C)**.

**Fig. 5.**
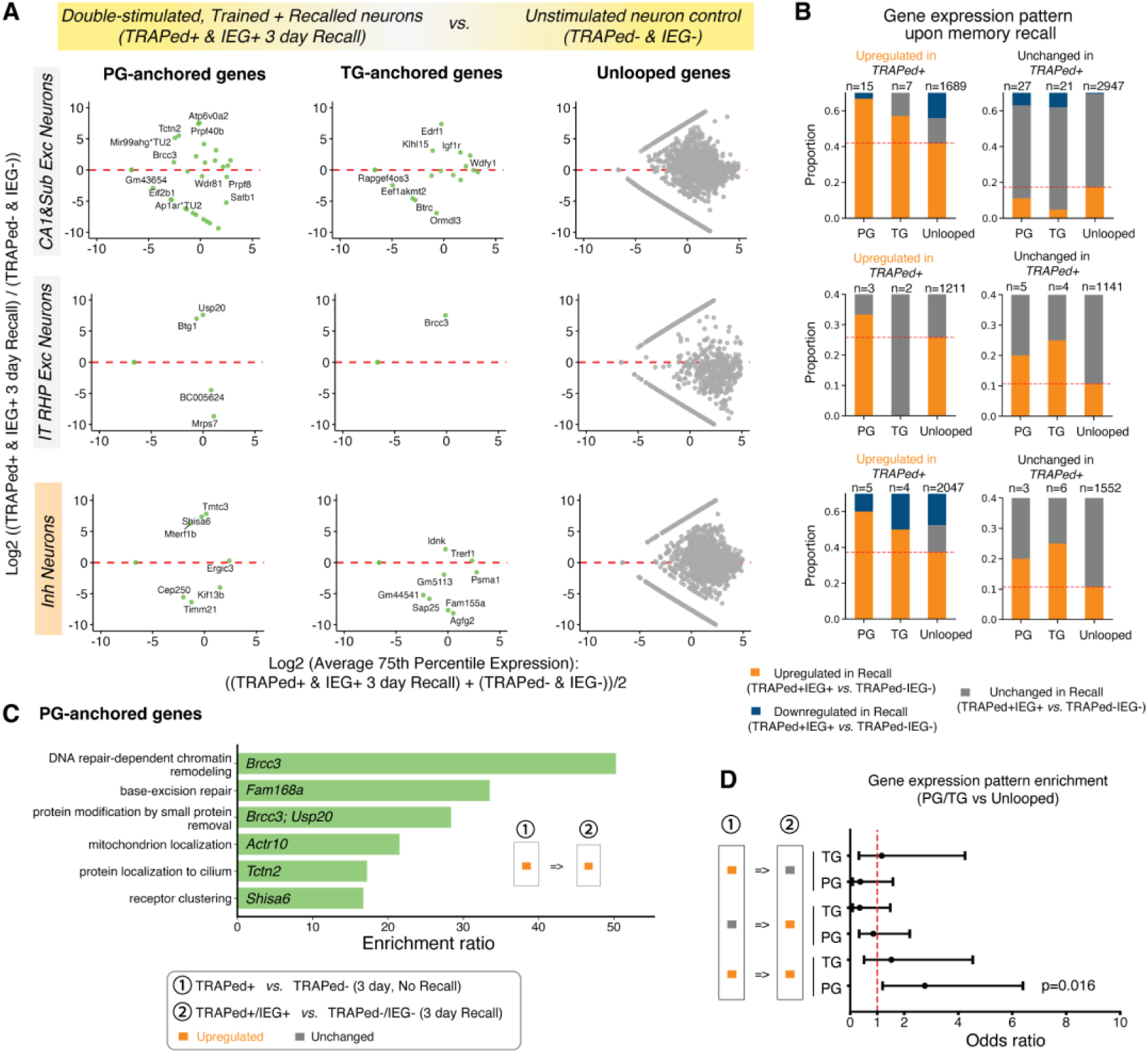
Persistently gained promoter-enhancer loops are associated with robust and selective upregulation of CFC-induced genes during memory recall. **(A)** M-A plots comparing double-stimulated (TRAPed+IEG+ at 3 days recall) to unstimulated (TRAPed-IEG-) neurons after CFC and recall. Left, persistently gained promoter-enhancer loop-anchored genes; middle, transiently gained promoter-enhancer loop-anchored genes; right, unlooped genes. X-axis: log2 average 75th-percentile across the two conditions; Y-axis: log2 fold change between conditions. (**B**) Proportion of genes upregulated (orange), downregulated (blue), or unchanged (grey) upon recall in TRAPed+IEG+ versus TRAPed-IEG-neurons. PG, genes anchoring persistently gained promoter-enhancer loops; TG, genes anchoring transiently gained promoter-enhancer loops; Unlooped, genes not engaged in looping interactions. Column 1 represents genes upregulated at 3 days post-CFC in TRAPed+ vs. TRAPed-neurons. Column 2 represents genes unchanged at 3 days post-CFC in TRAPed+ vs. TRAPed-neurons. (**C**) Gene ontology for persistently gained loop-anchored genes that were upregulated in TRAPed+ vs. TRAPed-neurons at 3 days post-CFC and further upregulated in TRAPed+IEG+ vs. TRAPed-IEG-neurons upon memory recall. (**D**) Odds ratio representing association of genes exhibiting specific expression profiles at 3 days post-CFC and 3 hours post-recall when anchoring persistently gained and transiently gained loops vs. unlooped.

We computed odds ratios and conducted a Fisher’s Exact test to compare the proportion of genes with expression changes during recall. Persistently gained promoter-enhancer loops were significantly enriched for genes showing persistent upregulation (orange-orange, i.e., up in TRAPed+ vs. TRAPed- and further up in TRAPed+IEG+ vs. TRAPed-IEG-during recall) compared to transiently gained loop and unlooped controls (**Fig. 5D**, **fig. S32**; OR = 2.76, P = 0.016). Genes anchoring transiently gained loops showed no significant enrichment across any loop class. Together, our data reveal a strong correlation between the upregulation of gene expression, in particular genes encoding DNA repair, during memory recall and persistently gained promoter-enhancer loops.

### Persistent loops anchor genes associated with post-traumatic stress disorder and autism

Finally, given the reported dysregulation of fear, anxiety, and memories in both post-traumatic stress disorder (PTSD) (*65, 66*) and autism spectrum disorder (ASD) (*67–69*), we examined genes with published associations with PTSD or ASD. We found that genes associated with PTSD (*70*) were significantly enriched at persistently lost promoter-enhancer loops compared to unlooped controls in excitatory neurons (odds ratio = 5.21, Pvalue = 0.0034), with a weaker but consistent trend in inhibitory neurons (odds ratio = 3.29, Pvalue = 0.0673) (**Fig. 6A**). Notable genes anchoring persistently lost loops include *LPXN*, *OLFM1*, *RBFOX1*, *SLC4A1AP*, and *ZFP91* in excitatory neurons, and *FOXP1*, *FOXP2*, and *JAK1* in inhibitory neurons. Among these genes, *RBFOX1* is well known for its role in synaptic plasticity through BDNF-TrkB signaling, a pathway critical for fear memory consolidation (*71, 72*), while *FOXP* genes control cortical development and fear-related circuits (*73*). We also found that genes associated with ASD were significantly enriched at persistently lost and persistently weakened promoter-enhancer loops in inhibitory neurons (odds ratio = 1.85, Pvalue = 0.0245), with a weaker but consistent trend in excitatory neurons (odds ratio = 1.60, Pvalue = 0.0766) (**Fig. 6B**). Genes associated with ASD were also significantly enriched at persistently gained and persistently strengthened promoter-enhancer loops in excitatory neurons (**Fig. 6B**, odds ratio = 2.13, Pvalue = 0.0130). Notable genes at persistently lost/weakened-anchored loops in inhibitory neurons encode synaptic adhesion and neurotransmitter signaling (*CTNND2*, *NRXN1*, *GABRG3*, *GRM5*, *SNAP25*) and at persistently gained/strengthened-anchored loops in excitatory neurons include chromatin/protein homeostasis regulators (*ARID2*, *UBE2H*, *UBR3*), suggesting that persistent loops can anchor both synaptic and chromatin programs disrupted in ASD. Together, these data demonstrate that CFC-induced chromatin loop persistence can anchor crucial synaptic and chromatin regulatory genes associated with PTSD and ASD.

**Fig. 6.**
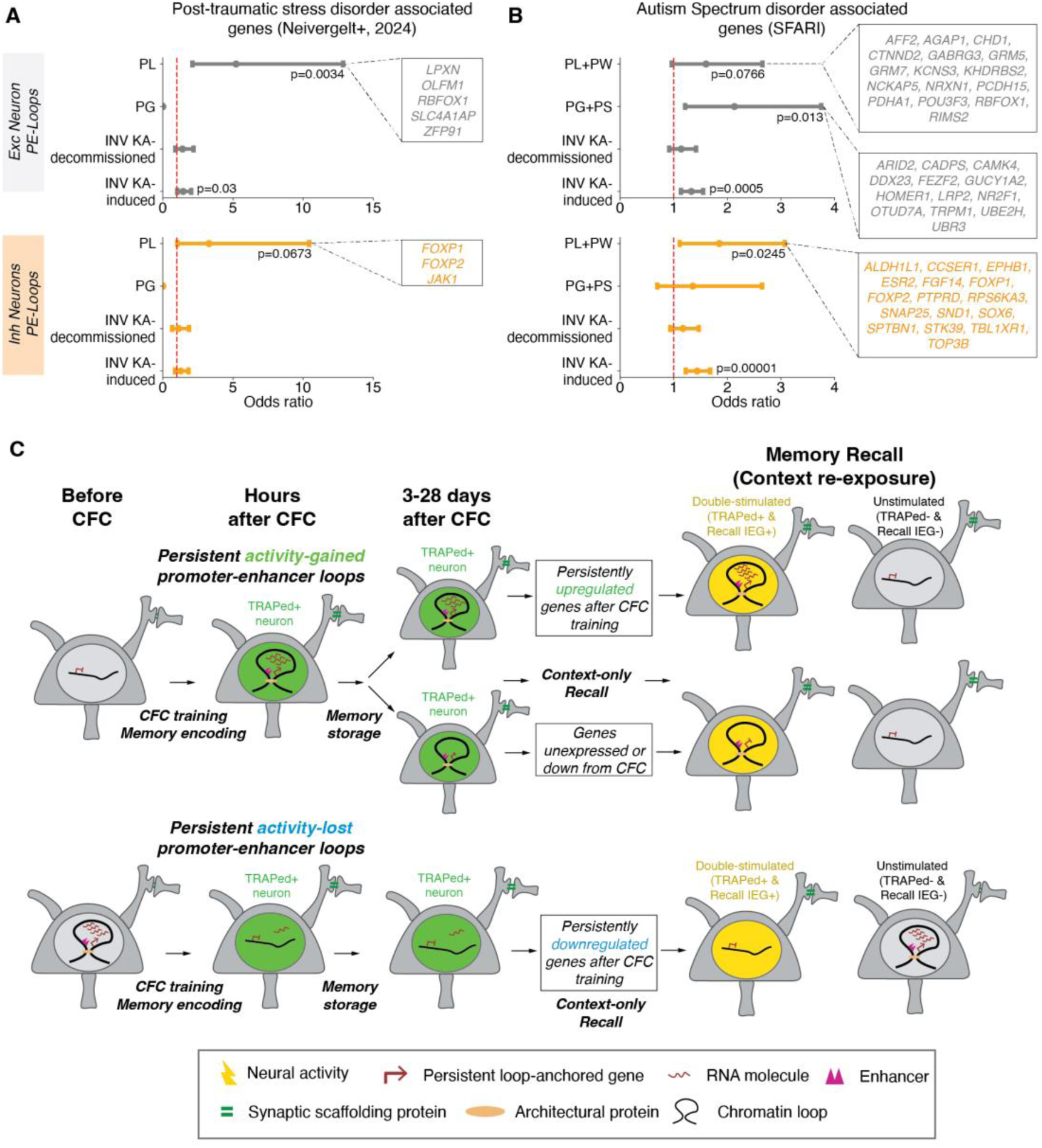
Persistent promoter-enhancer chromatin loops anchor genes associated with post-traumatic stress disorder and autism spectrum disorder. (**A-B**) Odds ratio enrichment of loop-anchored gene sets at published human risk loci for **(A)** post-traumatic stress disorder (PTSD) and **(B)** autism spectrum disorder. (**C**) Proposed working model for the link between persistent gained and persistent lost promoter-enhancer loops and gene expression during memory encoding, storage, and recall.

## Discussion

Long-term durability of activity-dependent chromatin changes may occur on time scales of long-term memory storage, but limited direct evidence exists. Using multi-modal single-nucleus experimental and computational technologies paired with activity-dependent nuclear tagging in double-transgenic mice *in vivo*, we profiled DNA methylation and higher-order chromatin folding across thousands of single hippocampal neurons in a time course after CFC. We discovered CFC-induced genome-wide reorganization of chromatin loops in both excitatory and inhibitory neurons marked during learning. A subset of CFC-induced and CFC-decommissioned loops persisted for at least four weeks after learning. Persistently gained and lost loops were unique to each neural subtype and anchored distinct synaptic signaling-related genes. Altogether, our work demonstrates that cell type-specific chromatin loops induced by learning can endure for up to a month *in vivo*. Previous work using electron microscopy demonstrated the genome-wide restructuring of chromatin upon neural stimulation *in vitro* (*11*). We build on previous work by finding the specific locations of loops genome-wide that change during training *in vivo* and retain durable structural traces for weeks after memory encoding. We provide lists of synaptic genes anchored by persistently gained and lost promoter-enhancer loops and rigorously examine the effect of loop persistence upon recall. Our work sheds new light on the underappreciated role of higher-order chromatin folding in learning and memory.

It is established that a distributed ensemble of neurons is stimulated during training in CFC and that the neurons can be genetically labeled to permanently express GFP (*23*). We combined bleeding edge technologies with the TRAP2 mouse and uncovered two types of stable chromatin changes and their correlation with gene expression recall *in vivo* (**Fig. 6C**). We used single-nucleus gene expression profiling and found that persistent activity-gained promoter-enhancer loops were significantly enriched for genes initially upregulated at 3 days post-CFC that exhibit further robust upregulation in recall-responsive neurons upon memory retrieval. Conversely, persistently activity-lost promoter enhancer loops anchored genes exhibiting robust downregulation upon memory recall. It is particularly noteworthy that the gene expression effect of loop persistence occurs during recall, in that only those genes upregulated or downregulated by CFC at 3 days post-training were specifically and robustly further up and downregulated at persistently gained and lost loops, respectively. Altogether, we find that loop persistence is directly linked to robust recall of mRNA levels for crucial neuronal subtype-specific synapse signaling genes during fear memory recall. A subset of such genes has been associated in independent studies to PTSD and autism, thus highlighting the relevance of our observations for neurodevelopmental and neuropsychiatric disorders.

Because our neuron-tagging strategy relies on the IEG *Fos* as a proxy for neuronal activation, it likely captures only a subset of the full activity-dependent ensemble, as recent studies show only partial overlap between *Fos* and other IEGs such as *Arc* (*74*). Non-TRAPed neurons in our experimental system likely include additional activity-dependent neuronal ensembles that evade *Fos*-based tagging. Future studies employing multiplexed IEGs tagging or calcium activity-based tagging systems could resolve the full spectrum of memory ensembles and their associated epigenetic modifications (*74, 75*). Beyond the hippocampus, regions such as the prefrontal cortex and amygdala (*76*) will be important to investigate for tissue-specific chromatin folding and gene expression phenomena during memory encoding, consolidation, recall, and extinction. Finally, dissecting the molecular machinery that maintains persistent loops and testing their causal role in memory will be essential, with implications for therapeutic targeting in memory disorders.

Our finding provides a more holistic picture of memory storage that complements and builds upon traditional structural and functional synaptic plasticity models to include chromatin in the nucleus and the regulatory principles of cell-wide availability of RNA species (*77–80*). While previous studies identified various epigenetic modifications associated with learning (*11, 24–29*), our results demonstrate that learning induces 3D genome reorganization that persists for at least a month *in vivo* during time frames of memory storage. The nuclear engram we propose (**Fig. 6C**) anchors synaptic genes associated with PTSD and autism that exhibit robust regulation of mRNA levels during recall. We hypothesize that memory of long-range gene expression regulatory mechanisms through loop persistence influences cell-wide levels of crucial synaptic RNA species that in turn could in principle transport from nucleus to synapse to govern synaptic transmission and neurophysiology (*12, 81, 82*).

Our work represents the first step toward understanding chromatin-synapse communication in long-term memory storage and recall. It is a thorough characterization of loop persistence as a nuclear engram and plausible effects on gene expression recall *in vivo*. Altogether, our work opens up new areas of inquiry into the communication and interplay between nuclear chromatin, long-range gene expression regulation, and synapse-specific RNA mechanisms in normal neurophysiology and in neurological disorders.

## Acknowledgments

We thank members of the Cremins lab for helpful discussions.

## Author contributions

*Conceptualization:* PX, KRC, JS, EK, JEPC

*Methodology/Visualization:* PX, KRC, BL, RS, AP, CL, KT, AJW, KP, HR, JS, EK, JEPC

*Investigation:* PX, KRC, RS, JEPC

*Funding:* JEPC

*Administration:* JEPC

*Writing & Editing:* PX, KRC, BL, RS, JEPC

*Reagents:* PX, BL, AP, EK, JEPC

## Data and materials availability

All data is provided at GEO under the accessions: GSEXXXX (snm3C-seq3) and GSEXXXX (Smart-seq3). All data and code will be freely available upon publication.

