## Supplementary Materials for "Learning induces persistent chromatin loops underlying robust gene expression during memory recall"

**This PDF file includes:**

Materials and Methods
Supplementary Text
Figs. S1 to S33
Tables S1 to S16
Supplementary References

### Supplementary Figures

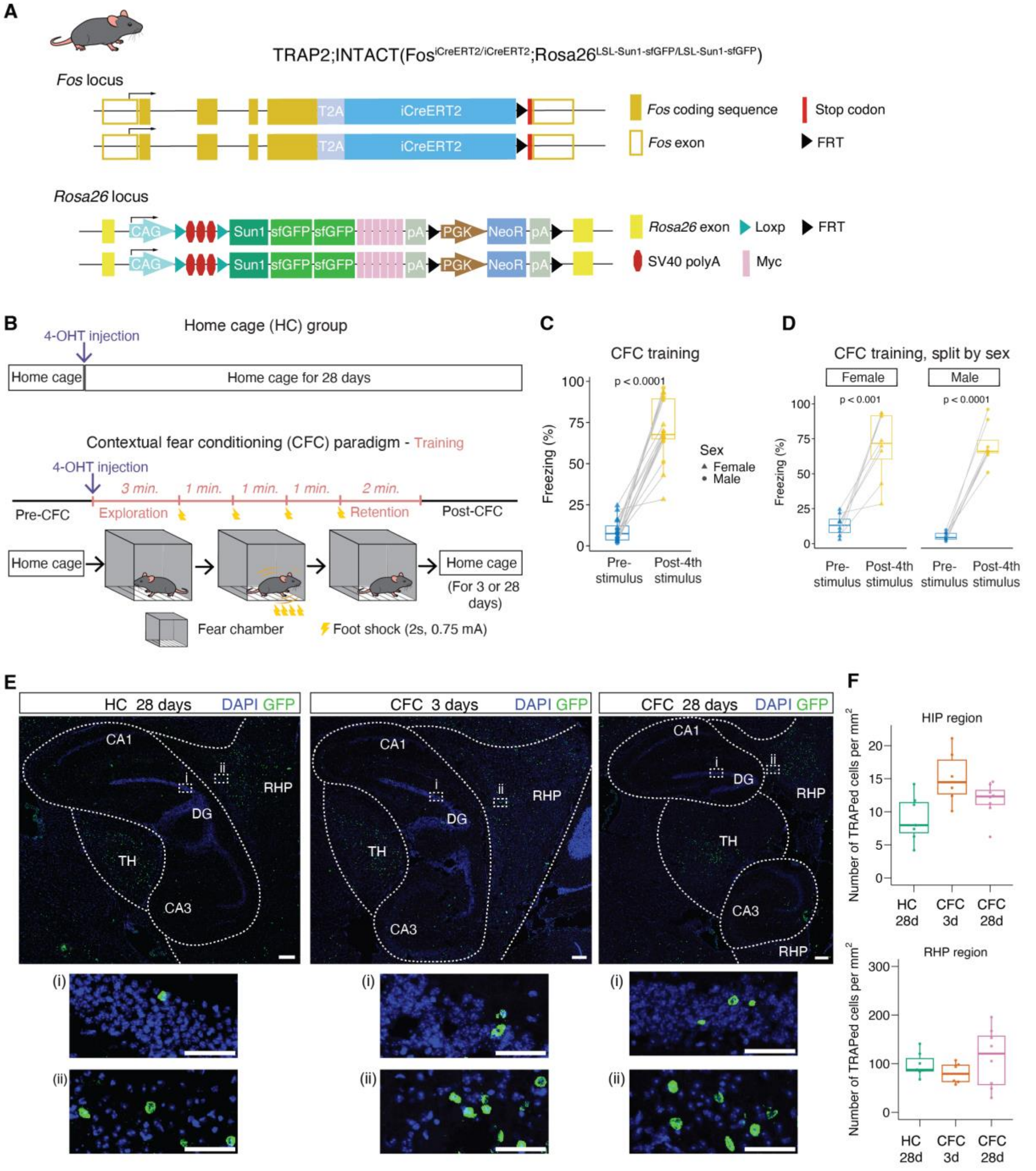


**Fig. S1. Behavioral analysis of TRAP2;INTACT transgenic mice in contextual fear conditioning.**

**(A)** Schematics of TRAP2; INTACT (Sun1sfGFP) transgenic mice. **(B)** Schematic of contextual fear conditioning protocol (see Materials and Methods for details). **(C-D)** Quantification of freezing behavior (corresponding to Fig. 1) with (C) and without (D) the separation of males and females. Box plots show the percentage of time mice spent freezing pre-stimulus and post fourth stimulus (n = 16; 8 males and 8 females). Circles represent males and triangles represent females. Pvalues were calculated using two-tailed paired t-tests. **(E)** Top panels: Example confocal images (E) with GFP and DAPI staining of in hippocampal region (HIP) and retrohippocampal region (RHP) in HC28d, CFC3d and CFC 28 days. Bottom panels: (i) and (ii) show higher magnification views within dashed boxes in the top panels of the HIP and RHP regions, respectively. Scale bars: 100 µm (E - top panels); 50 µm (E – bottom panels: magnified views, i and ii). **(F)** Quantifications of percentage of TRAPed (GFP+) cells in hippocampal region (HIP, top panel) and retrohippocampal region (RHP, bottom panel), respectively. Analysis included n=2 mice per group, with each dot representing one slice.

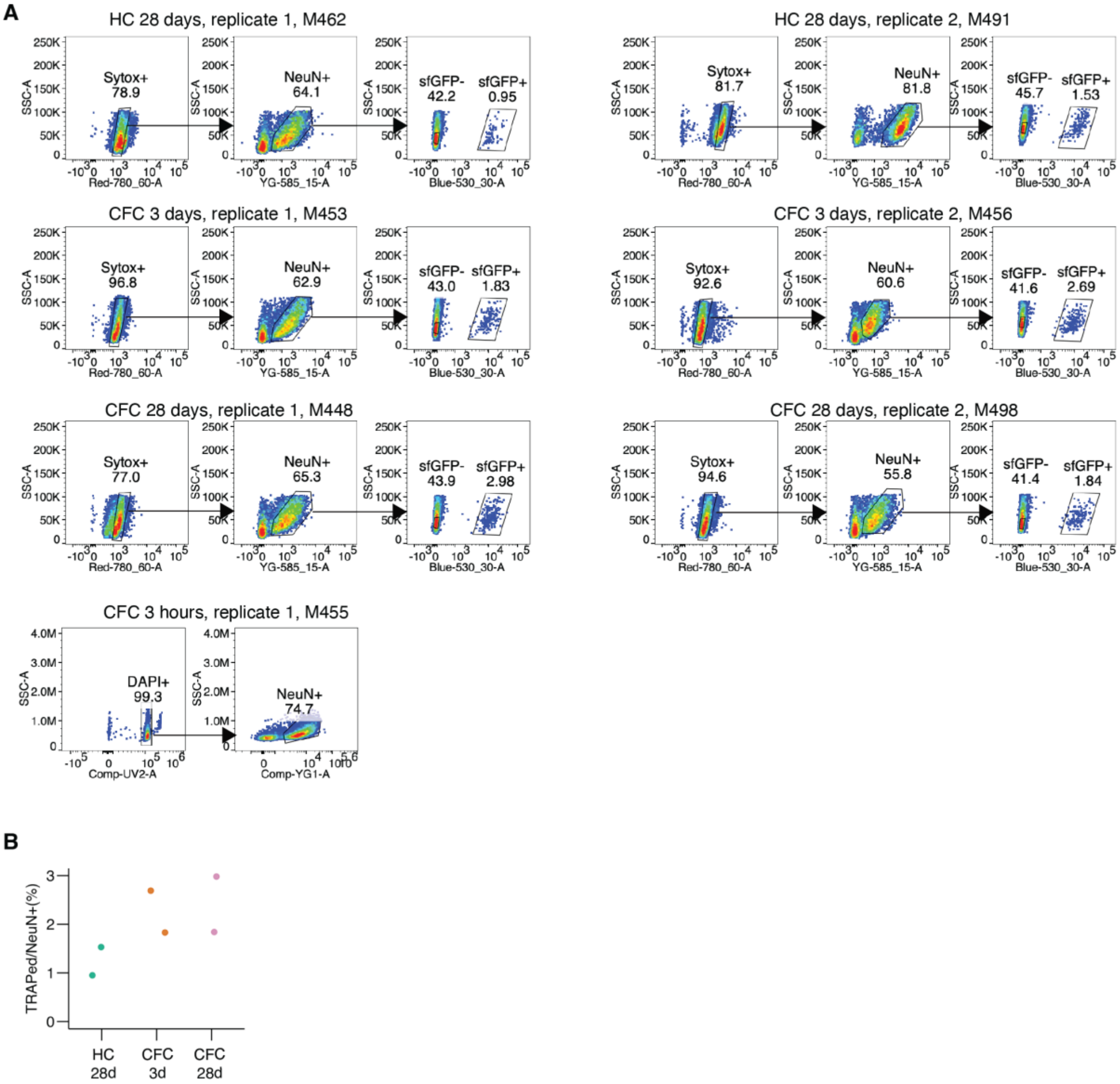


**Fig. S2. Representative flow cytometry plots and metrics of sorted neurons used for snm3C-seq experiments.**

**(A)** Flow cytometry analysis of hippocampal formation neurons under different behavioral conditions. Left panels: DAPI or Sytox staining for identification and gating of all nuclei. Middle panels: Neuronal nuclei selection using anti-NeuN antibody staining, gated on Sytox+ or DAPI+ population. Right panels: superfolder GFP (sfGFP) expression analysis, gated on the NeuN+ population. We collected NeuN+ nuclei for CFC3h group. **(B)** Strip plot showing the percentage of TRAPed cells among NeuN+ neurons for different experimental groups.

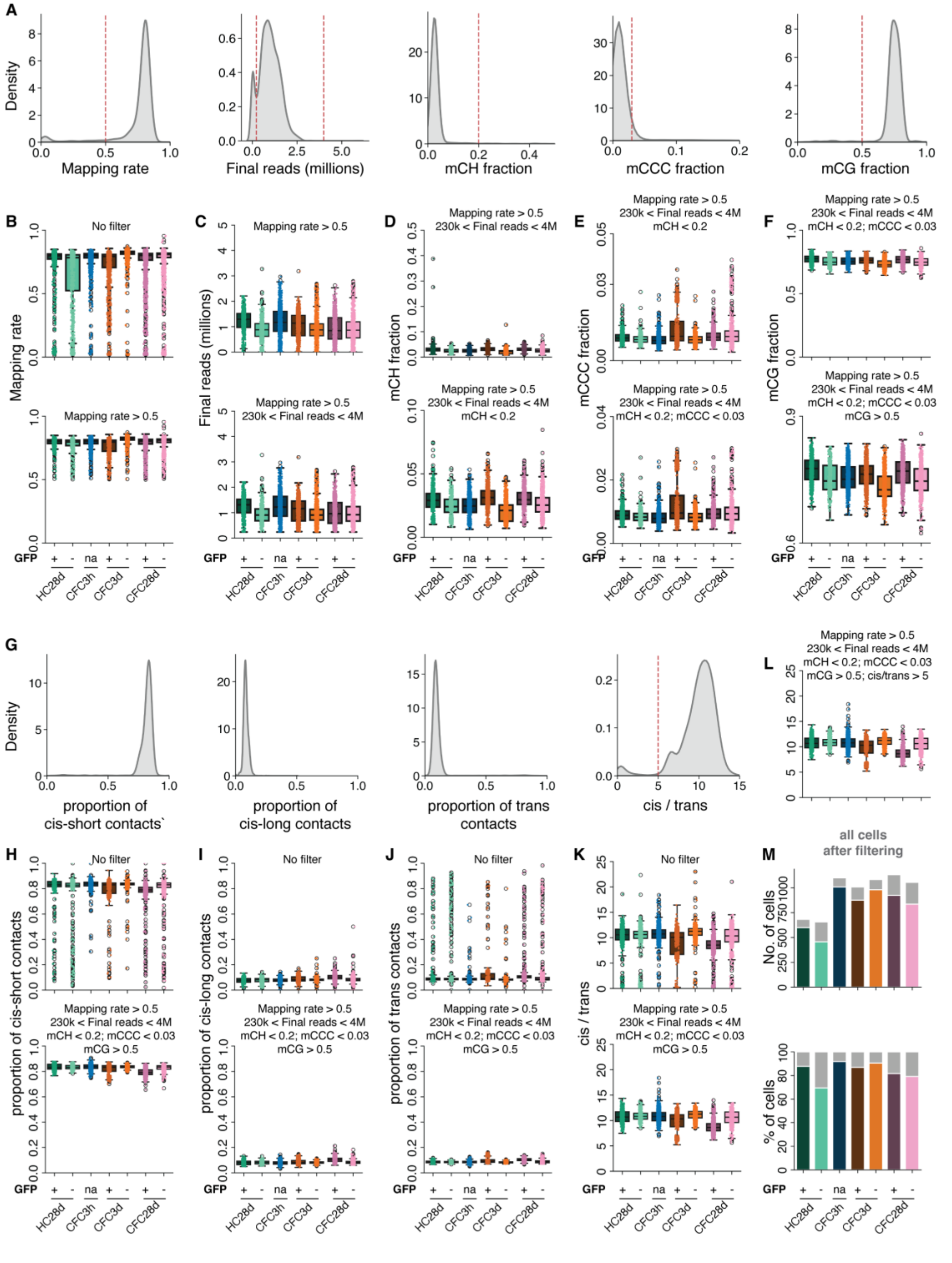


**Fig. S3. Quality control metrics to ensure rigorous snm3C-seq experimental data and analysis.**

**(A)** Quality control metrics of DNA methylation profiles. Gaussian kernel density estimates for mapping rate, final deduplicated reads, mCH fraction, mCCC fraction, and mCG fraction. Red dashed lines indicate filtering thresholds. **(B-F)** Boxplots with dots representing cells of quality control metrics across sorted cell populations before (top) and after (bottom) filtering in a cumulative manner. Mapping rate (B), Final deduplicated reads (C), mCH fraction (D), mCCC fraction (E), and mCG fraction (F). **(G)** Quality control metrics of chromatin contacts before applying filters on mapping rate, final deduplicated reads, mCH fraction, mCCC fraction, and mCG fraction. Gaussian kernel density estimates for proportions of cis-short range, cis-long range, trans contacts and cis/trans ratio. Red dashed lines indicate filtering thresholds. **(H-K)** Boxplots with dots representing cells of chromatin contact metrics across sorted cell populations before (top) and after (bottom) filtering for DNA methylation-based thresholds. Proportion of cis-short range contacts (H), Proportion of cis-long range contacts (I), Proportion of trans contacts (J), Cis/trans ratio (K). **(L)** Cis/trans ratio distribution after applying DNA methylation-based thresholds. Cells with a ratio below 5 were excluded. **(M)** Summary bar plots indicating the number of cells (top) and percentage of cells (bottom) at different stages of quality control. Gray bars represent cells before filtering, while colored bars represent cells after filtering based on various quality metrics: mapping rate, final deduplicated reads, mCH fraction, mCCC fraction, mCG fraction and cis/trans ratio.

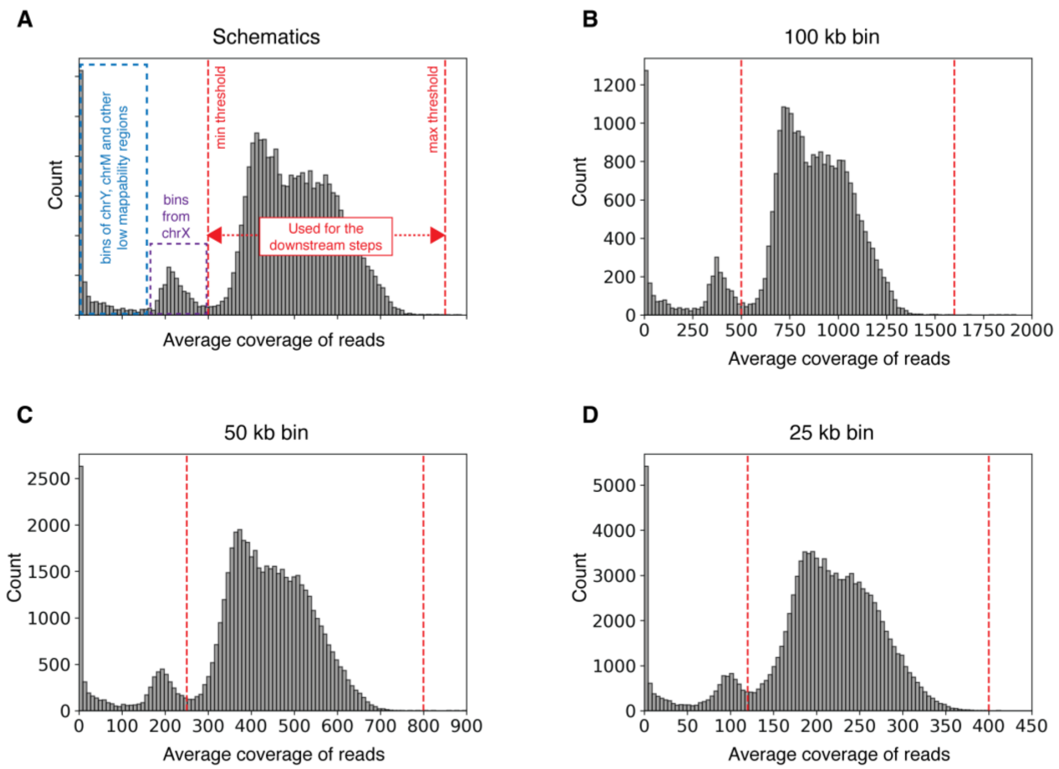


**Fig. S4. Average read coverage genome-wide across cells at different bin sizes.**
**(A)** Schematics of a histogram of average coverage of reads at a given resolution. Color-matched dashed boxes/lines and their corresponding labels represent different parts of range. **(B-D)** Histogram of average coverage of reads at 100 kb **(B)**, 50 kb **(C)**, and 25 kb **(D)** bin sizes.

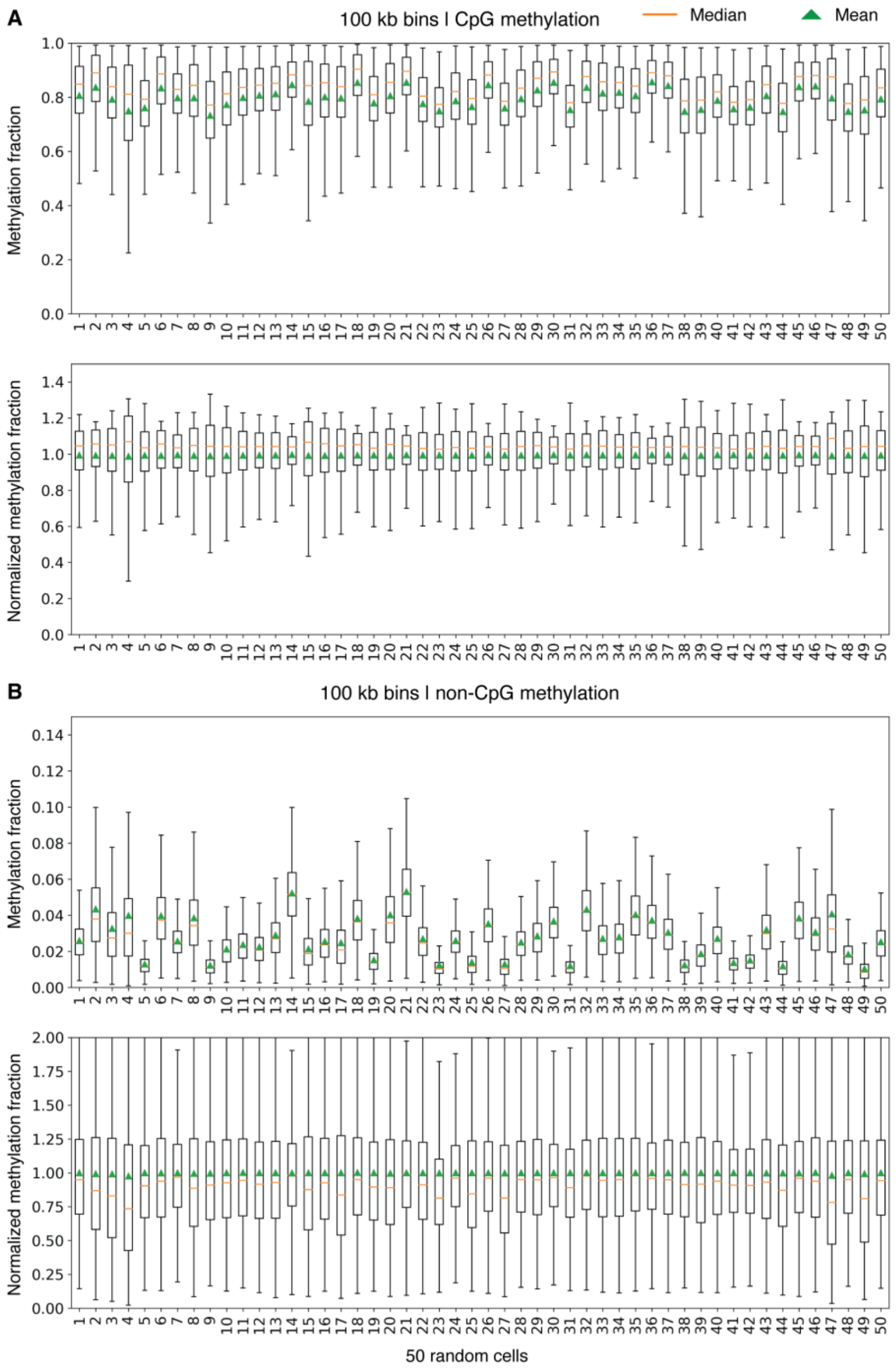


**Fig. S5. Single-neuron normalization of genome-wide DNA methylation at 100 kb bin resolution with the Beta distribution to correct for sequencing coverage differences.**
**(A)** Top: Raw CpG methylation fractions across 50 random cells. Bottom: Cell-to-cell normalization of the CpG methylation fractions across 50 random cells using beta normalization as described in (*83*). **(B)** Top: Raw non-CpG methylation fractions across 50 random cells. Bottom: Cell-to-cell normalization of the non-CpG methylation fractions across 50 random cells using beta normalization as described in (*83*).

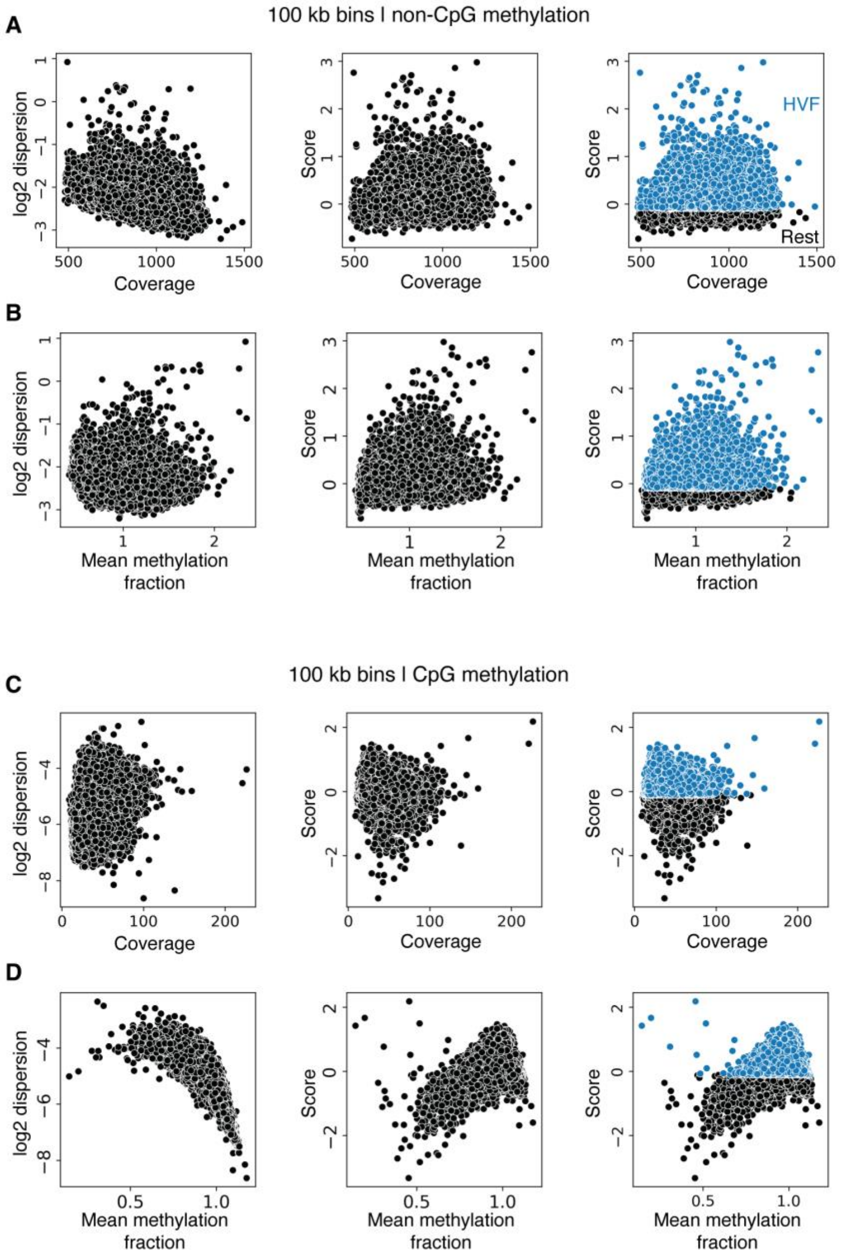


**Fig. S6. Selection of highly variable features (HVF) at 100 kb bin sizes.**
**(A, C)** Scatter plots showing the relationship between coverage and log2 dispersion (left), coverage and score (middle), and coverage over highly variable features highlighted in blue (right). **(B, D)** Scatter plots depicting the relationship between mean methylation fraction and log2 dispersion (left), mean methylation fraction and score (middle), and mean methylation fraction with HVFs highlighted in blue (right). Each point represents a 100 kb genomic bin. **(A, B)** non-CpG methylation and **(C, D)** CpG methylation.

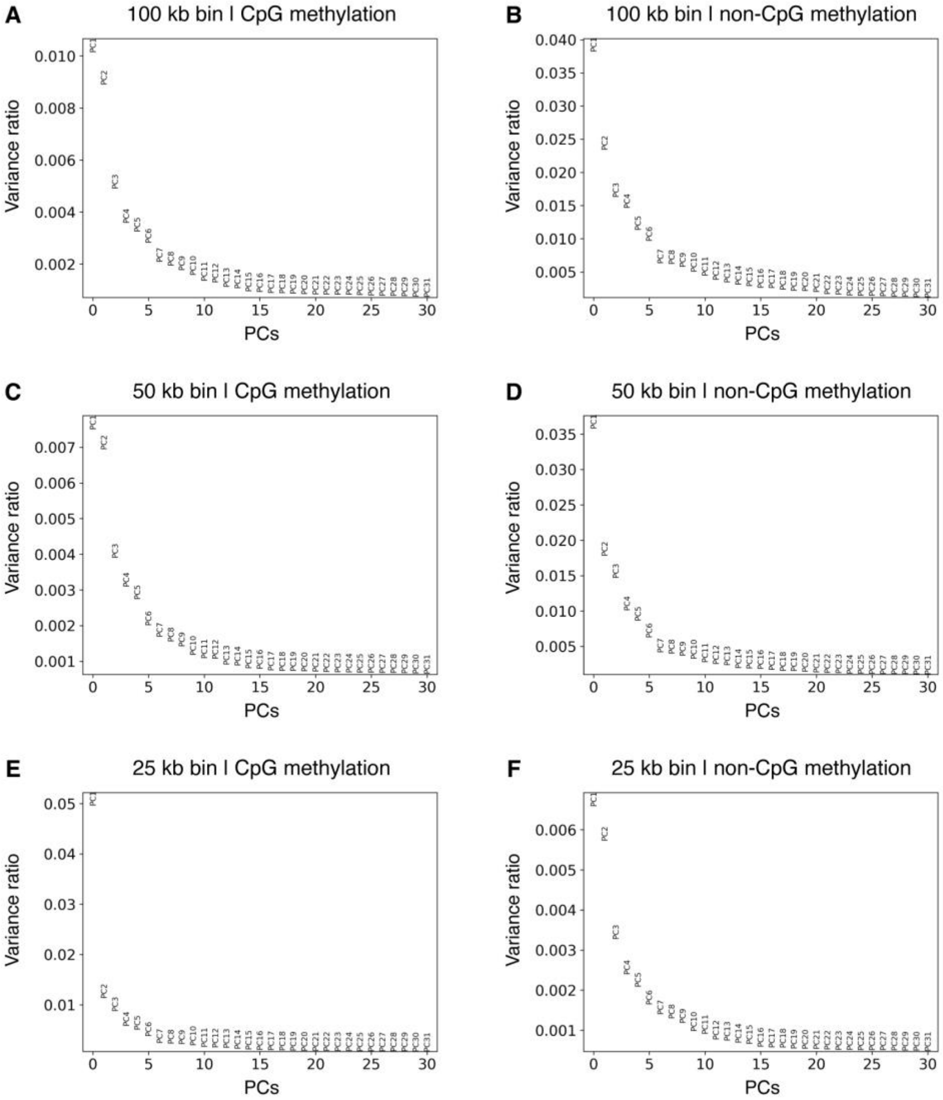


**Fig. S7. Variance ratio analysis of principal components (PCs) for DNA methylation patterns across different genomic bin sizes and methylation contexts.**
**(A, C, E)** Variance ratios of first 31 PCs CpG methylation. **(B, D, F)** Variance ratios of first 31 PCs non-CpG methylation. **(A, B)** 100 kb bins, **(C, D)** 50 kb bins, **(E, F)** 25 kb bins.

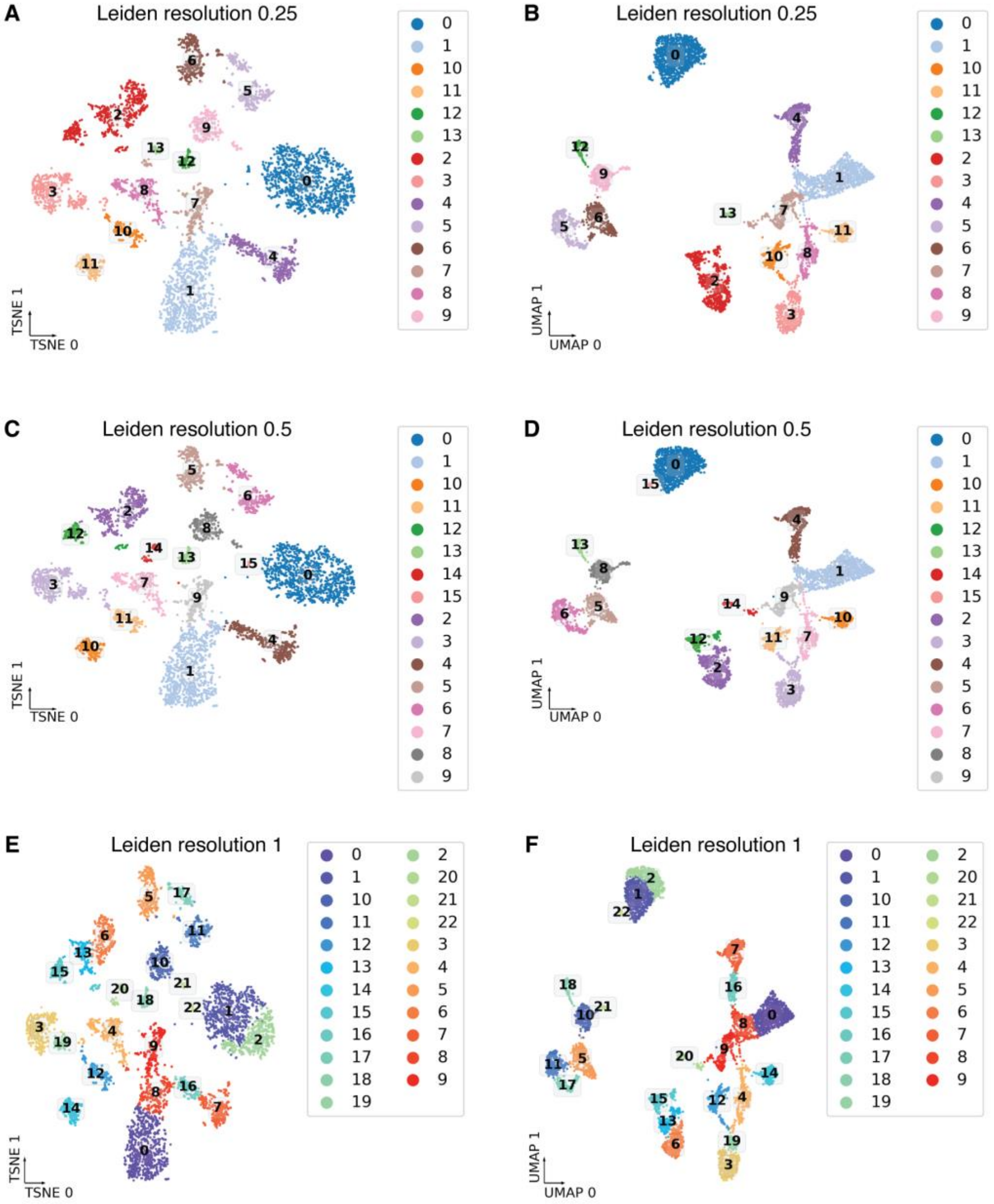


**Fig. S8. Dimensionality reduction and clustering analysis of snm3C-seq3 data using different Leiden resolution parameters at 100 kb resolution.**

**(A, C, E)** Clustering results of different Leiden resolutions on t-SNE embeddings. **(B, D, F)** Clustering results of different Leiden resolutions on UMAP embeddings. (A, B) Leiden resolution = 0.25; (C, D) Leiden resolution = 0.5; (E, F) Leiden resolution = 1. We selected the parameters of Leiden resolution 0.25 and 100 kb bins for the cell type clusters used for analyses in the manuscript.

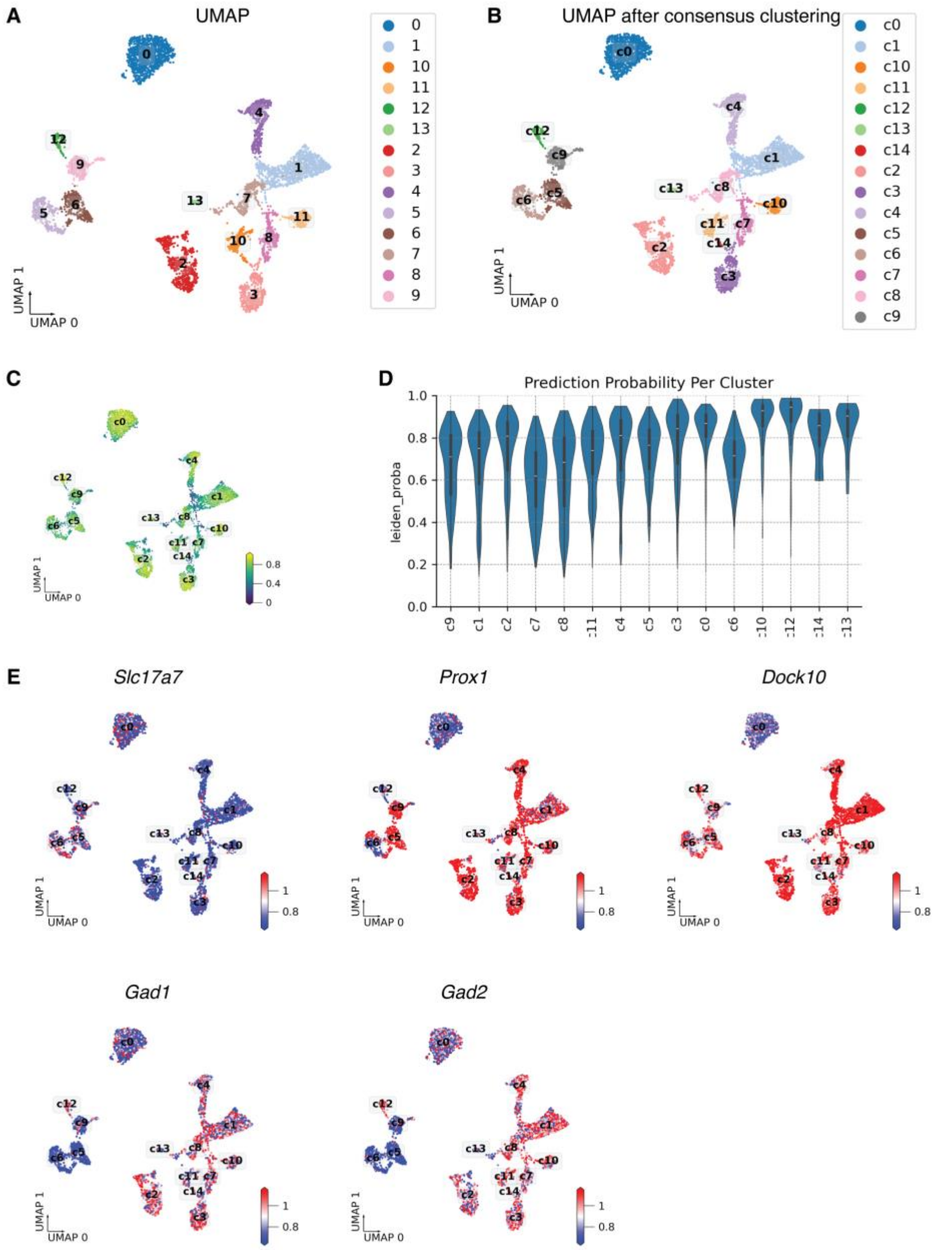


**Fig. S9. Consensus clustering, prediction probability, and non-CpG DNA methylation at 100 kb bin size overlayed with gene expression of neural subtype specific markers.**
**(A)** UMAP visualization of initial cell clusters at 100 kb bin size. **(B)** UMAP plot after consensus clustering with 100 rounds of Leiden clustering to counteract the randomness. **(C)** UMAP plot displaying cluster prediction probabilities for every cell. **(D)** Violin plot of prediction probabilities for each consensus cluster. **(E)** Normalized non-CpG methylation fraction within gene body +/- 2kb regions of marker genes of glutamatergic (*Slc17a7*), DG (*Prox1*, *Dock10*), INH (*Gad1*, *Gad2*) neurons overlaid on UMAP plots. Color intensity represents relative non-CpG methylation, with red indicating high levels (i.e., indicator of inactive gene) and blue indicating low levels (i.e., indicator of active gene).

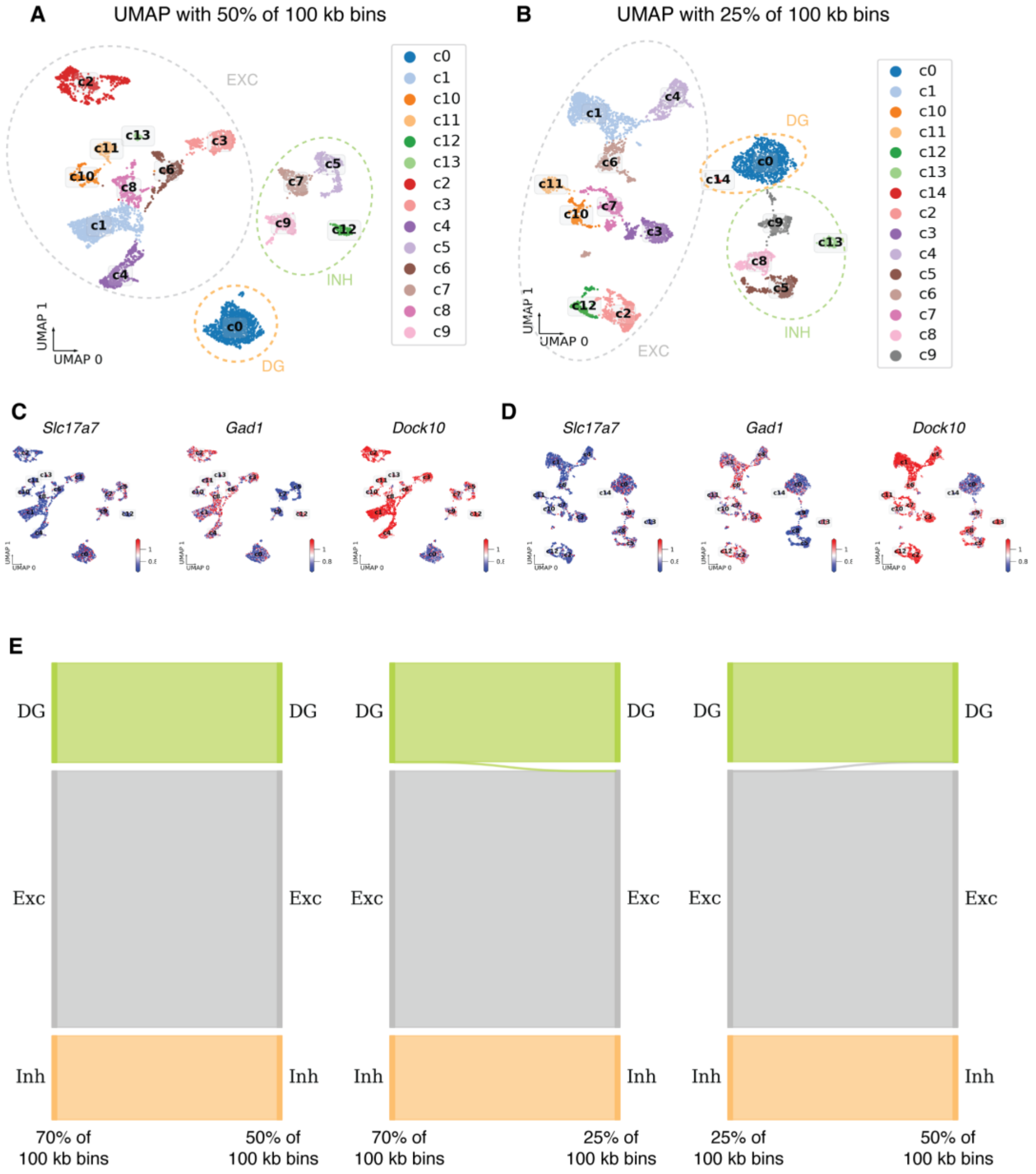


**Fig. S10. Comparison of major class annotation across three different highly variable feature (HVF) n_top_feature parameters.**

**(A-B)** UMAP visualization of initial cell clusters at 100 kb bin size with 50% **(A)** and 25% **(B)** of the bins during HVF. **(C-D)** Normalized non-CpG methylation fraction within gene body +/- 2kb regions of marker genes of excitatory (*Slc17a7*), DG (*Dock10*), inhibitory (*Gad1*) neurons overlaid on their UMAP plots in **(A)** and **(B)**. Color intensity represents relative non-CpG methylation, with red indicating high levels (i.e., indicator of inactive gene) and blue indicating low levels (i.e., indicator of active gene). **(E)** River plots showing the overlap among major neuron class annotation using 25%, 50% and 70% of the 100 kb bins during HVF. We selected the usage of 70% of the highly variable feature bins for the analyses in this paper.

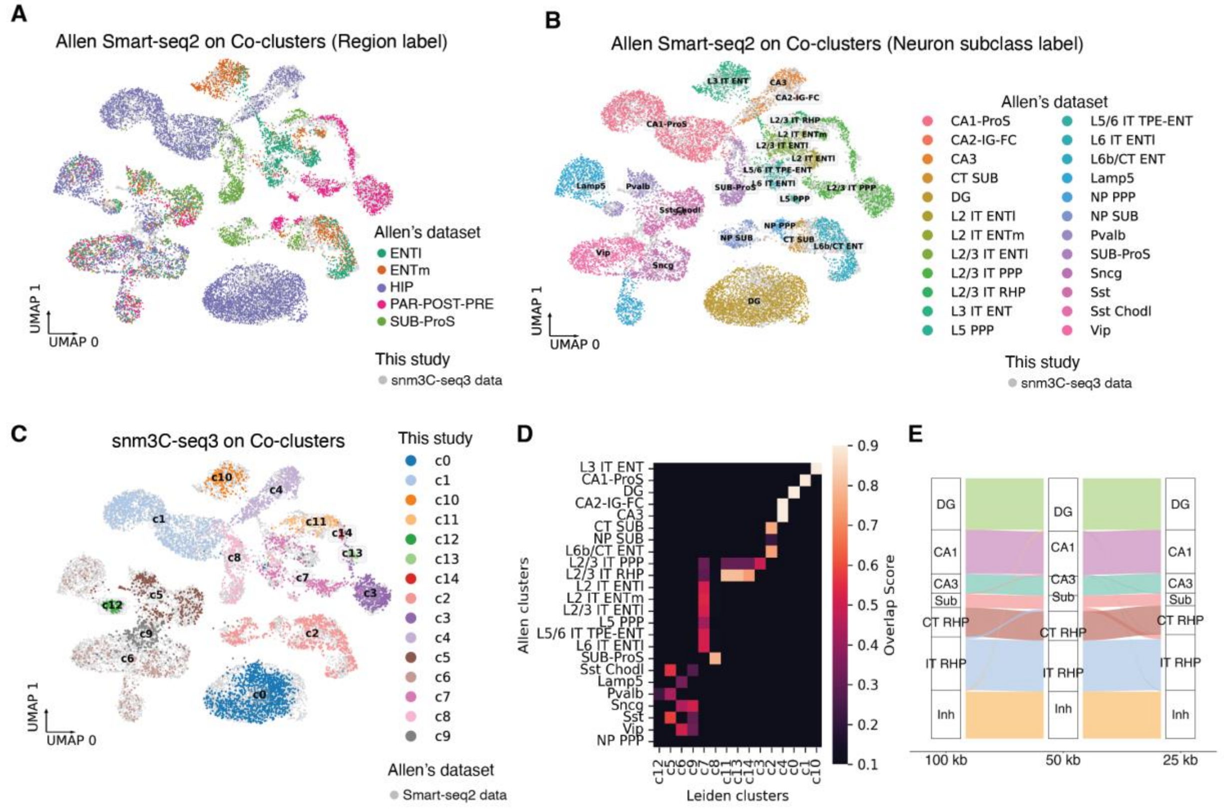

**Fig. S11. Annotation of neural subtypes in snm3C-seq3 data via integration with Allen Brain Atlas Smart-seq2 gene expression data.**

(**A-C**) UMAP plots displaying clustering results for snm3C-seq3 data and Allen Brain Atlas Smart-seq2 hippocampal formation neuronal data (*41*). We used the inverse relationship between gene body DNA methylation and gene expression to use DNA methylation profiles genome-wide and established atlases of single-neuron gene expression to define neural subtypes. **(A):** Allen Brain Atlas Smart-seq2 data with brain region labels projected onto integrated data, grey dots represent our snm3C-seq3 data; **(B):** Allen Brain Atlas Smart-seq2 neuronal subclass annotations projected onto integrated data, grey dots represent our snm3C-seq3 data; **(C):** snm3C-seq3 data projected onto integrated data, grey dots represent Allen Brain Atlas Smart-seq2 data. **(D)** Confusion heatmaps showing overlap scores between Allen Smart-seq2 clusters (y-axis) and snm3C-seq3 Leiden clusters (x-axis) Brain regions labels and neuronal subtype annotations are inherited from and Allen Brain Atlas Smart-seq2 hippocampal formation neuronal data (*41*). (**E**) Sankey diagram illustrating the relationships between neuronal subtypes clustering results across different bin sizes (100 kb, 50 kb, and 25 kb) for the snm3C-seq3 data. The width of the connecting bars indicates the proportion of cells shared between subtypes across different bin size clusters.

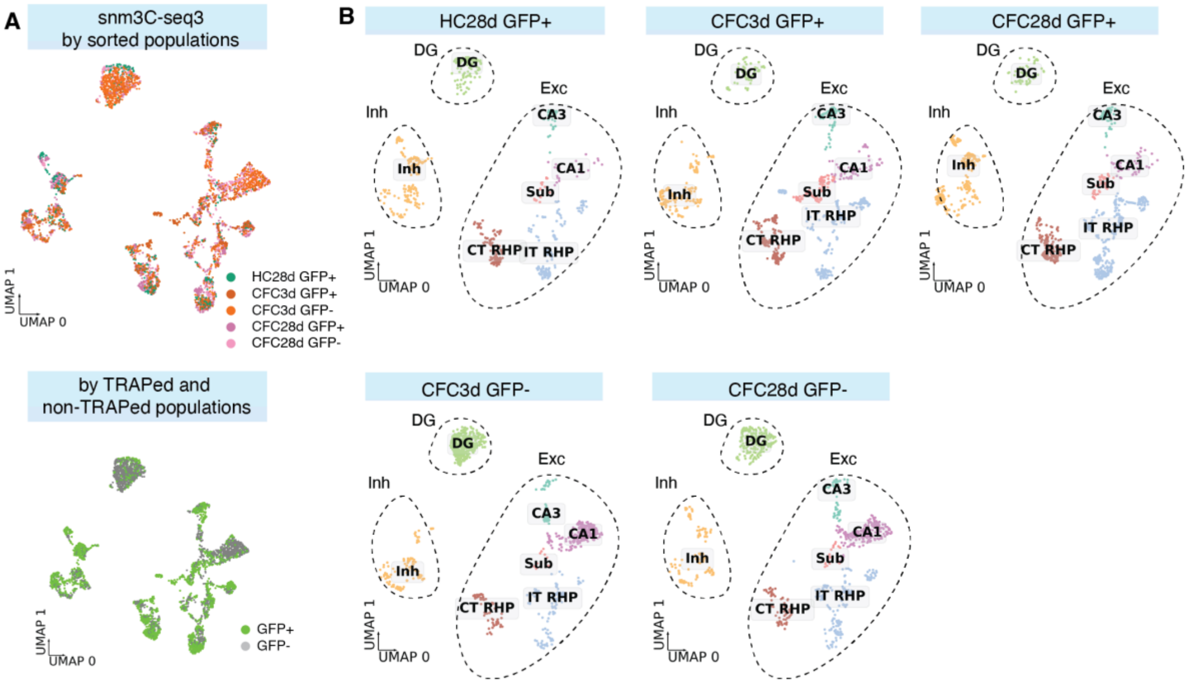


**Fig. S12. Neural subtype composition from hippocampal tissue across TRAPed GFP+ and unTRAPed GFP- cells at 3 days and 28 days post-training with contextual fear conditioning.**

**(A)** UMAP plots showing cell clusters for different behavioral conditions (top) and for TRAPed or non-TRAPed populations (bottom). **(B)** UMAP plots showing cell clusters split by different behavioral conditions of (A, top panel).

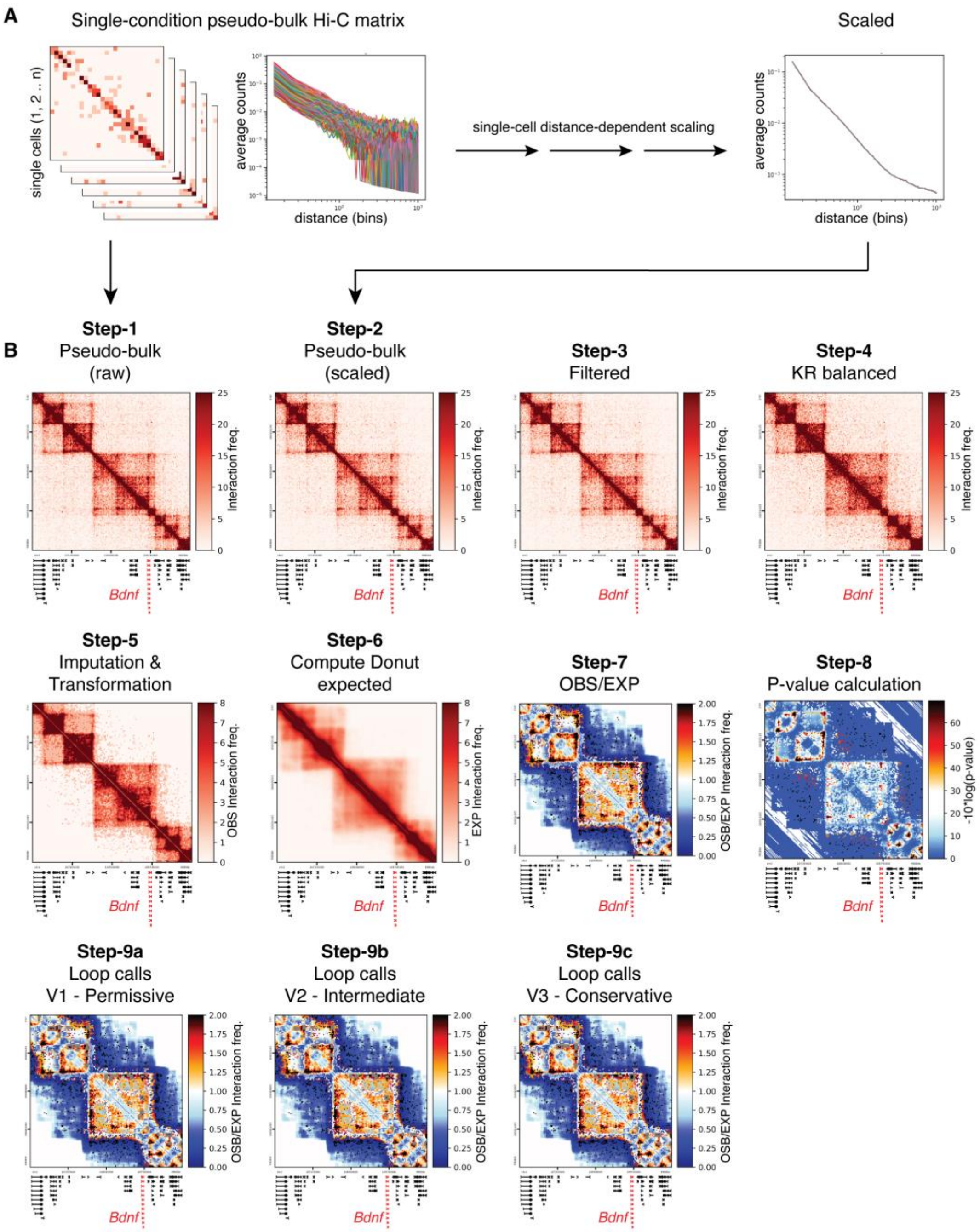


**Fig. S13. Analysis pipeline for rigorous detection and evaluation of long-range chromatin looping interactions in snm3C-seq data.**

**(A)** Schematics showing the scaling of contact maps across single cells using Hi-C contact distance-dependent scaling method. **(B)** Heatmaps show the contact maps around the gene *Bdnf* at every step of the loop calling pipeline: pseudobulking before scaling (step-1), pseudobulking after scaling (step-2), filtering (step-3), KR balancing (step-4), imputation (step-5), multiplication with a scalar (step-6), calculating donut expected values (step-7), calculating observed/expected (step-8), obtaining pvalues (step-9) and loop calling (step-10 v1-3). Resolution: 30 kb bins.

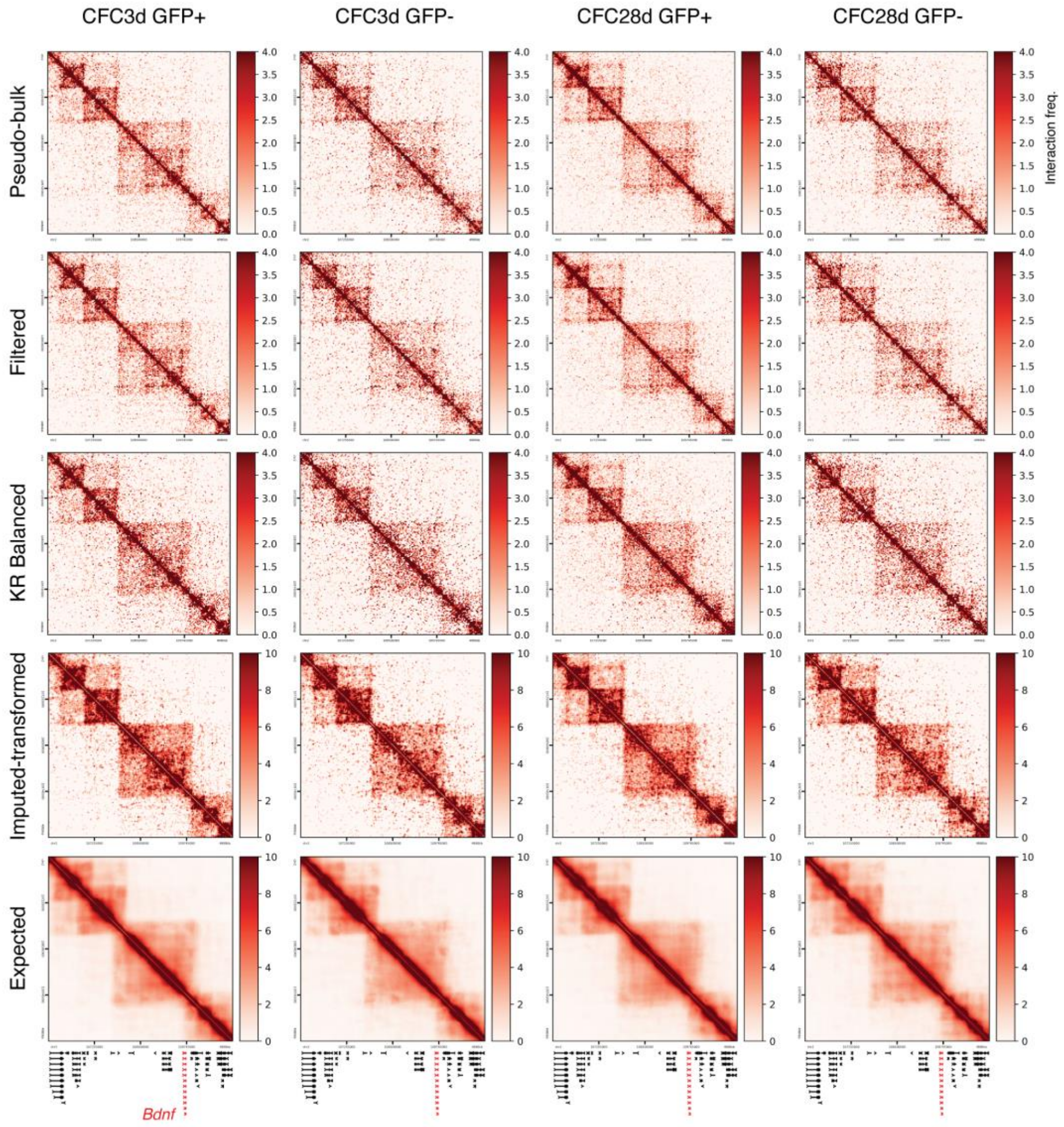


**Fig. S14. Observed and Expected interaction frequency heatmaps at the *Bdnf* locus in TRAPed+ and TRAPed- neurons at 3 days and 28 days after contextual fear conditioning.** Heatmaps show the contact maps around the gene *Bdnf* at every step of computing pseudobulked observed matrices. Phases from Fig. S13 pipeline shown: Scaling (step-2), filtering (step-3), KR balancing (step-4), imputation & transformation (step-5), calculating the Donut EXPECTED (step-6) for pseudobulk neuronal snm3C-seq heatmaps: CFC3d GFP+, CFC3d GFP-, CFC28d GFP+ and CFC28d GFP. Resolution: 30 kb.

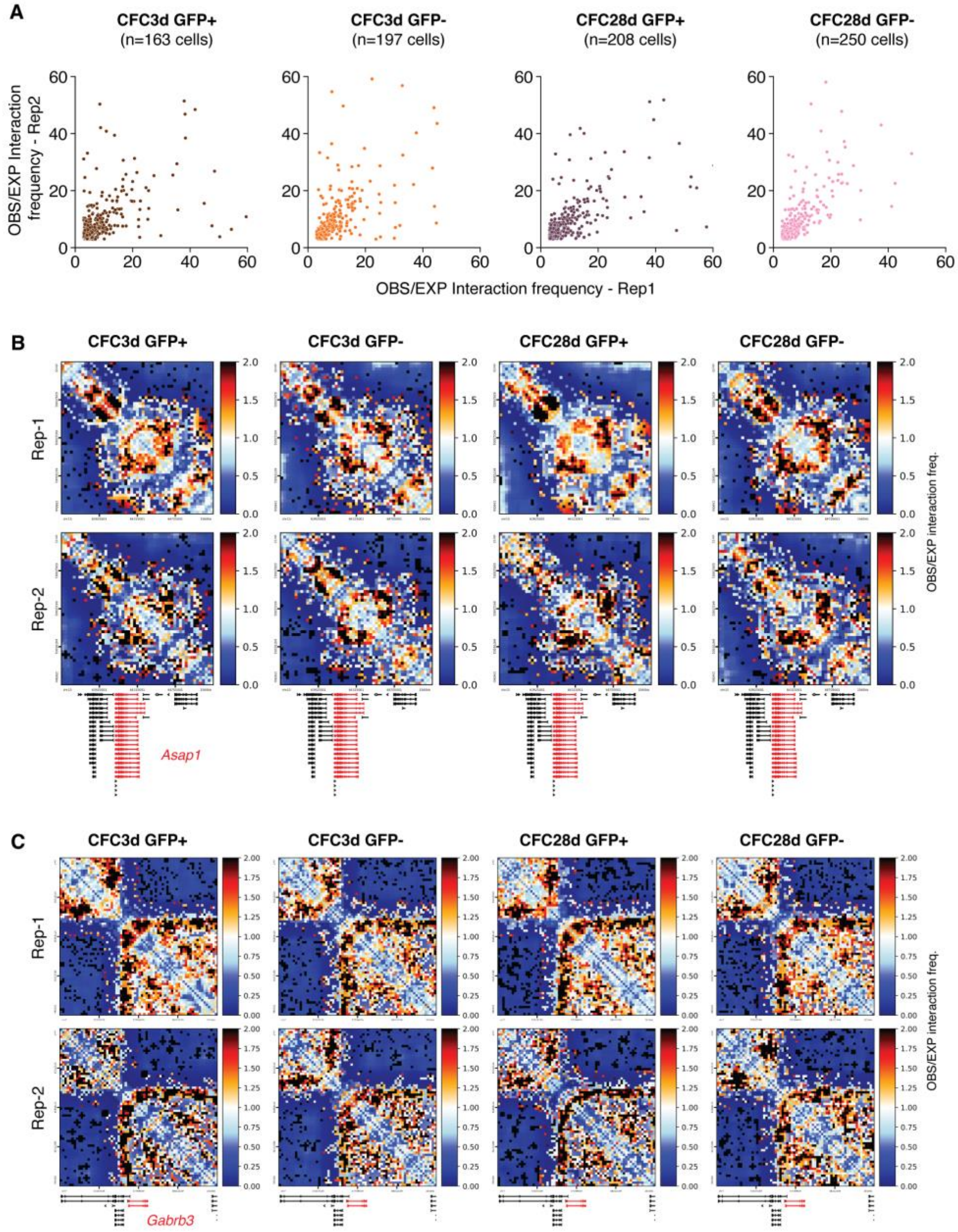


**Fig. S15. Chromatin loop analysis between two mouse replicates per condition showing strong replicate concordance at 3 days and 28 days post-training in TRAPed+ and TRAPed- neurons.**

**(A)** Scatter plots show OBS/EXP interaction frequencies between biological replicates across different experimental conditions: CFC3d GFP+ (n=163 cells), CFC3d GFP- (n=197 cells), CFC28d GFP+ (n=208 cells), and CFC28d GFP- (n=250 cells). Cell number (n) represent the number of cells after downsampling between the two replicates. Each dot represents the interaction frequency of an individual loop, with Spearman correlation coefficients (ρ) shown for each population. **(B-C)** Heatmaps show the OBSERVED/EXPECTED contact maps around **(B)** *Asap1* and **(C)** *Gabrb3* for replicate 1 (top panel) and 2 (bottom panel).

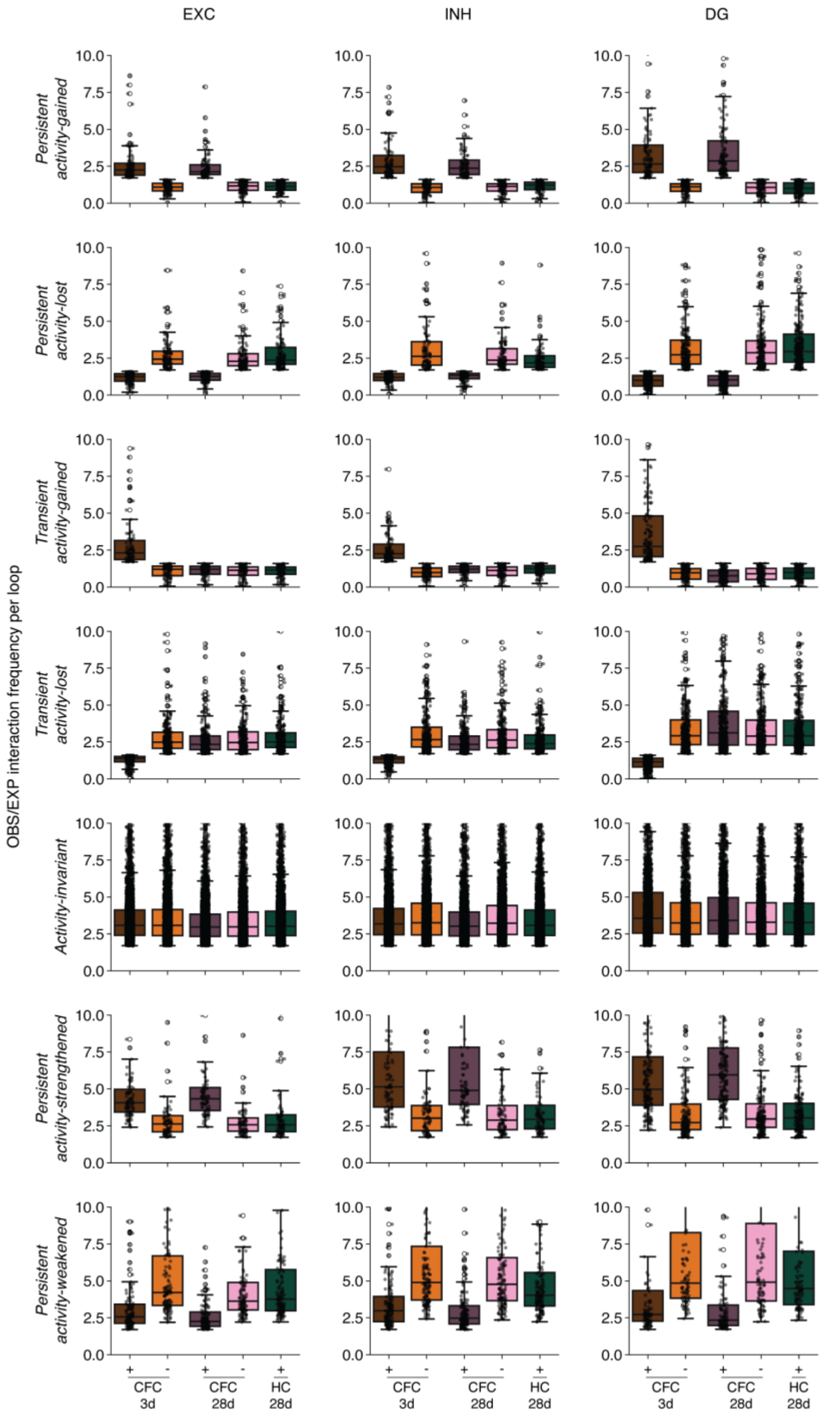


**Fig. S16. Chromatin loop classification across excitatory, inhibitory and DG subtypes.** Boxplots show the distribution of mean Observed/Expected interaction frequency per loop for persistent activity-gained (row 1), transient early activity-gained (row 2), persistent activity-lost (row 3), transient early activity-lost (row 4), invariant (row 4), persistent activity-strengthened (row 5), and persistent activity-weakened (row 6) loops across excitatory, inhibitory, and DG subtypes. Each dot in boxplots represents a loop.

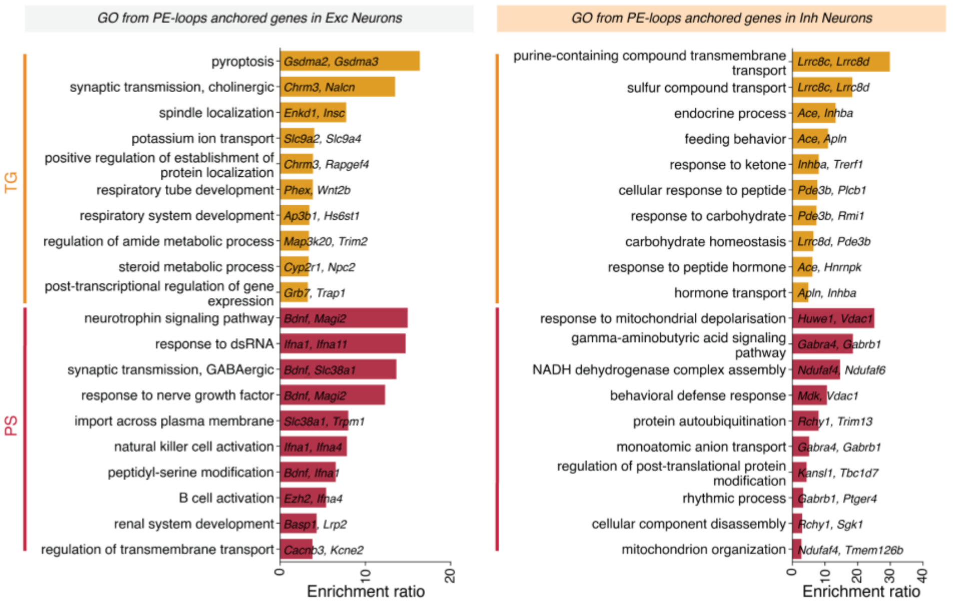


**Fig. S17. Ontology analysis of genes anchoring transient activity-gained and persistent activity-gained P-E loops called in excitatory and inhibitory neuron populations.**
Gene Ontology biological process enrichment for genes anchored by persistent strengthen (PS) and transient activity-gained (TG) P-E loops in excitatory and inhibitory neurons.

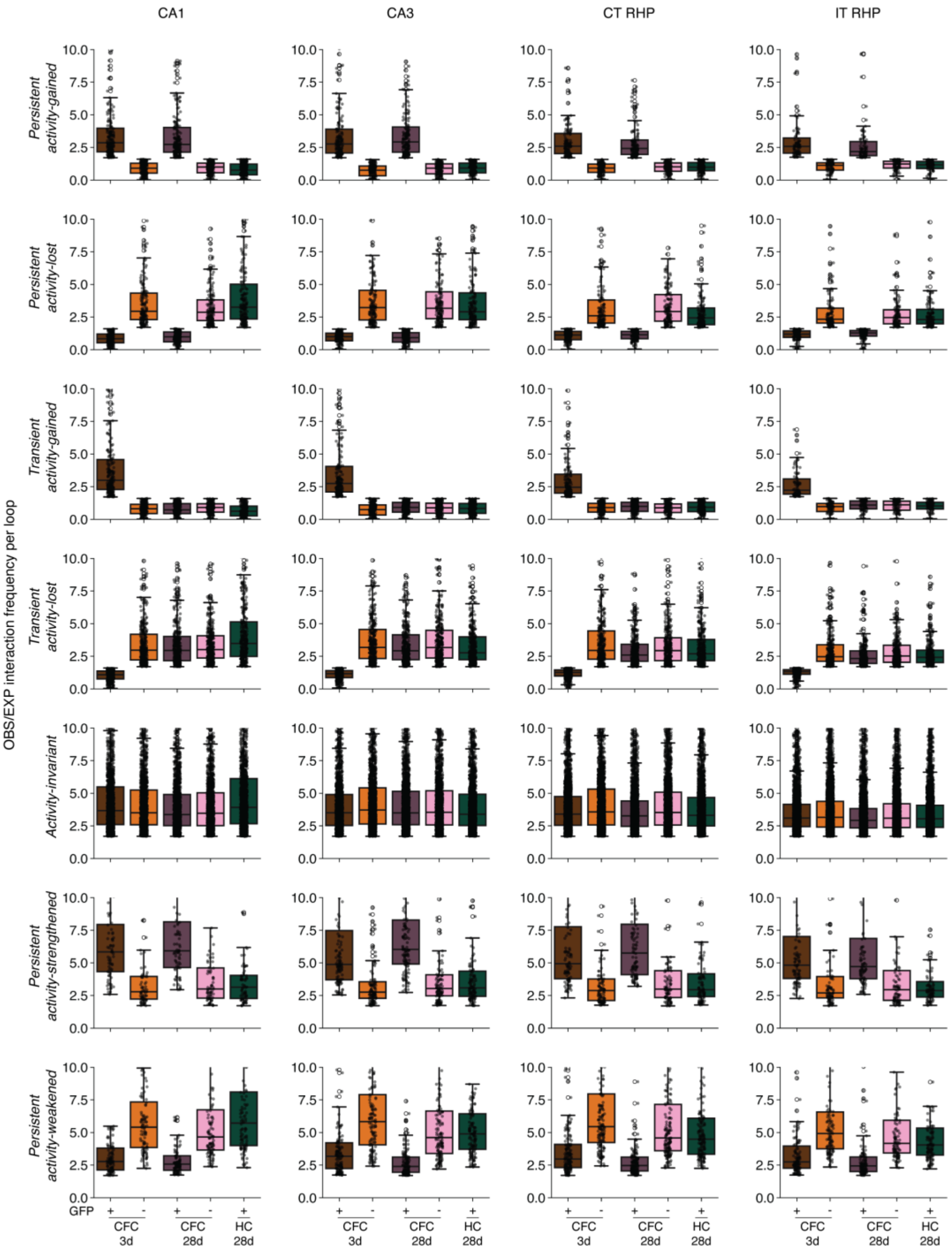


**Fig. S18. Chromatin loop classification across excitatory neuron subtypes.**
Boxplots show the distribution of mean Observed/Expected interaction frequency per loop for persistent activity-gained (row 1), transient early activity-gained (row 2), persistent activity-lost (row 3), transient early activity-lost (row 4), invariant (row 4), persistent activity-strengthened (row 5), and persistent activity-weakened (row 6) loops across excitatory, inhibitory, and DG subtypes. Each dot in boxplots represents a loop.

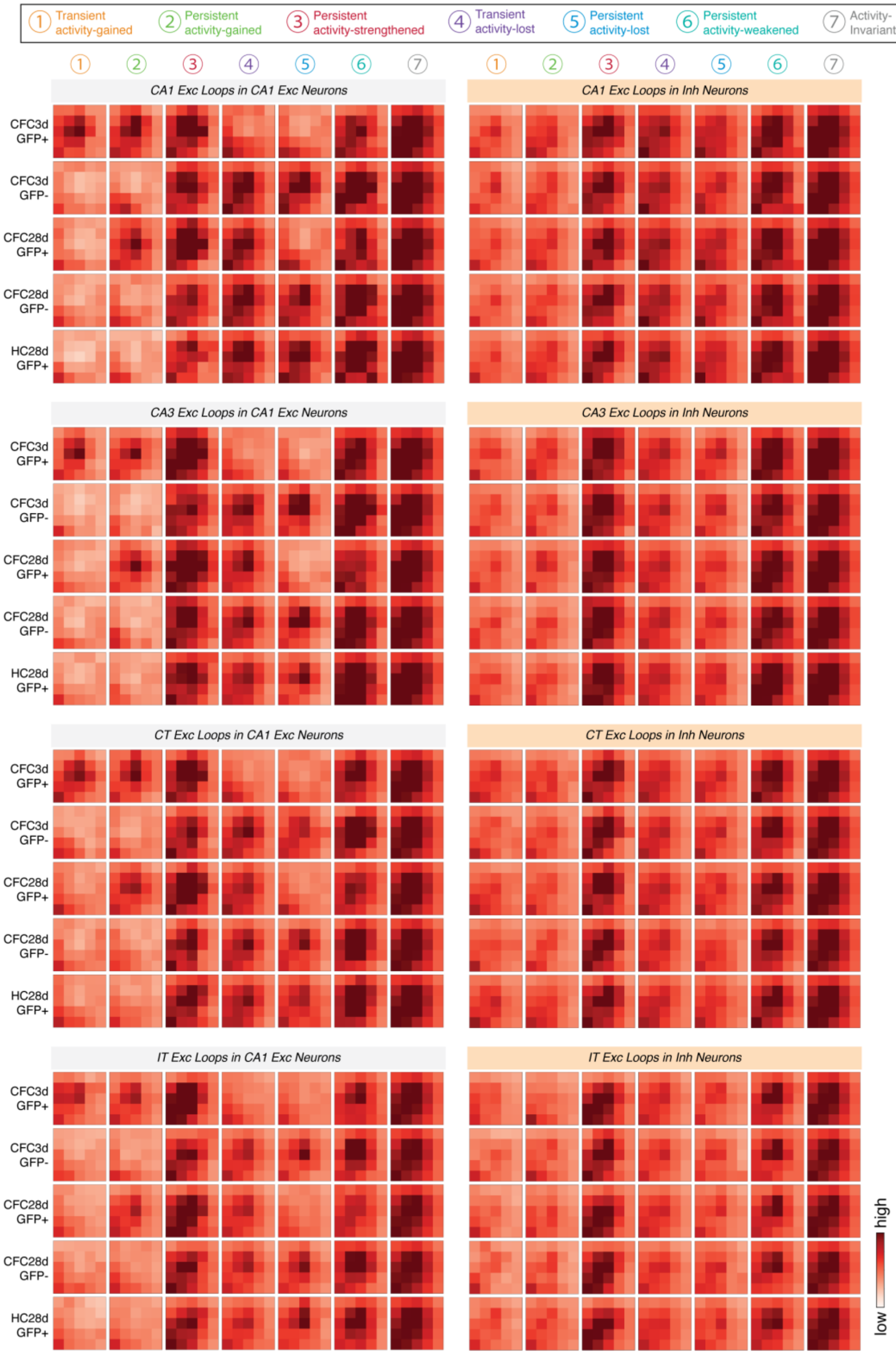


**Fig. S19. Aggregate Peak Analysis heatmaps for loop classes called independently in the excitatory neuron subtypes found in CA1, CA3, CT, and IT compared to inhibitory neuron snm3C-seq data.**

We plot the seven classes of chromatin loops identified at 3 days and 28 days post-training accounting for TRAPed+ and TRAPed- conditions. Loop classes were called independently on a merge of all excitatory subtypes (Fig. S16) and here on individual excitatory neural subtypes from CA1, CA3, CT, and IT. Examination of signal at the excitatory subtype loop classes in inhibitory neuron snm3C-seq data shows that loop classes are highly subtype specific.

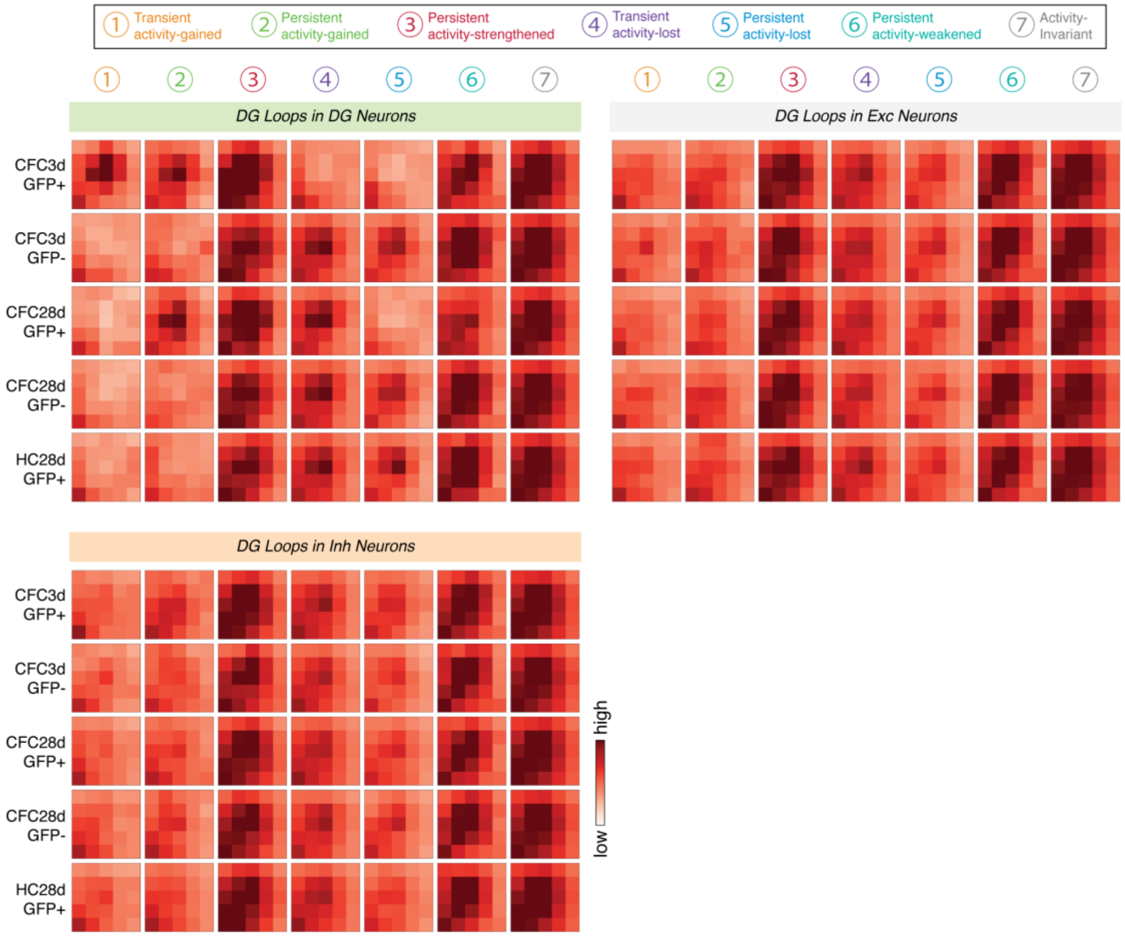


**Fig. S20. Aggregate Peak Analysis heatmaps for loop classes called independently in DG neurons.**

We plot the seven classes of chromatin loops identified at 3 days and 28 days post-training accounting for TRAPed+ and TRAPed- conditions. Loop classes were called independently on a merge of all DG neurons. Examination of signal at DG loop classes in excitatory and inhibitory neuron snm3C-seq data shows that DG loop classes are highly specific to the DG neurons.

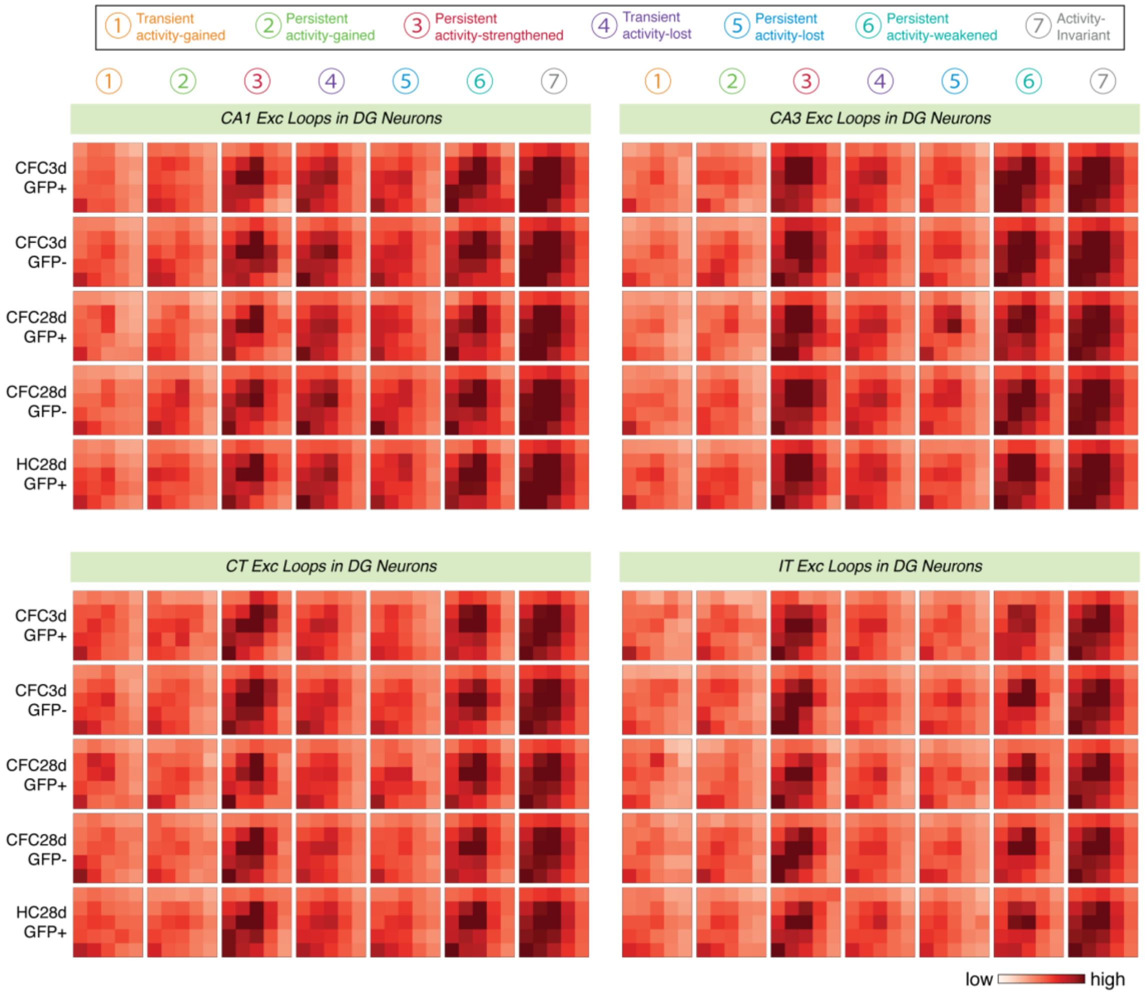


**Fig. S21. Aggregate Peak Analysis heatmaps for loop classes called independently in the excitatory neuron subtypes found in CA1, CA3, CT, and IT compared to DG neuron snm3C-seq data.**

We plot the seven classes of chromatin loops identified at 3 days and 28 days post-training accounting for TRAPed+ and TRAPed- conditions. Loop classes were called independently on a merge of all excitatory subtypes (Fig. S16) and also here on individual excitatory neural subtypes from CA1, CA3, CT, and IT. Examination of signal at the excitatory subtype loop classes in DG neuron snm3C-seq data shows that loop classes are highly subtype specific.

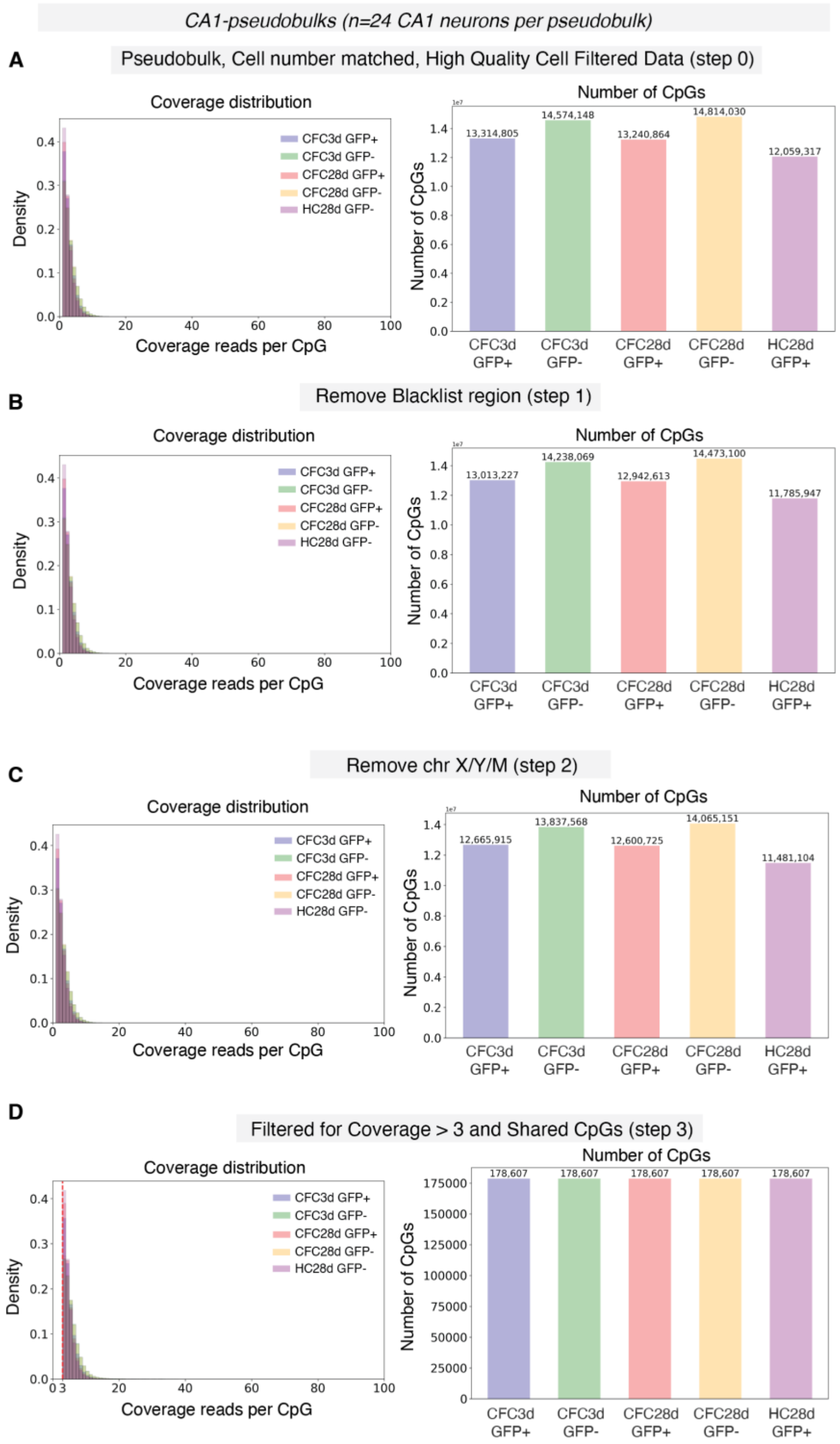


**Fig. S22. DNA methylation data processing at CpG resolution for CA1 neurons.**
(**A**) Pseudobulked raw data (step 0). (**B**) Filtered dataset after removal of CpGs overlapping with blacklist regions (step 1). (**C**) Filtered dataset after removal of CpGs belonging to chromosomes X, Y, and mitochondria (step 2). (**D**) Final filtered dataset containing only CpG sites shared across all four pseudobulked groups (step 3). Each stage displays three metrics: sequencing coverage distribution (left), and number of detected CpGs (right).

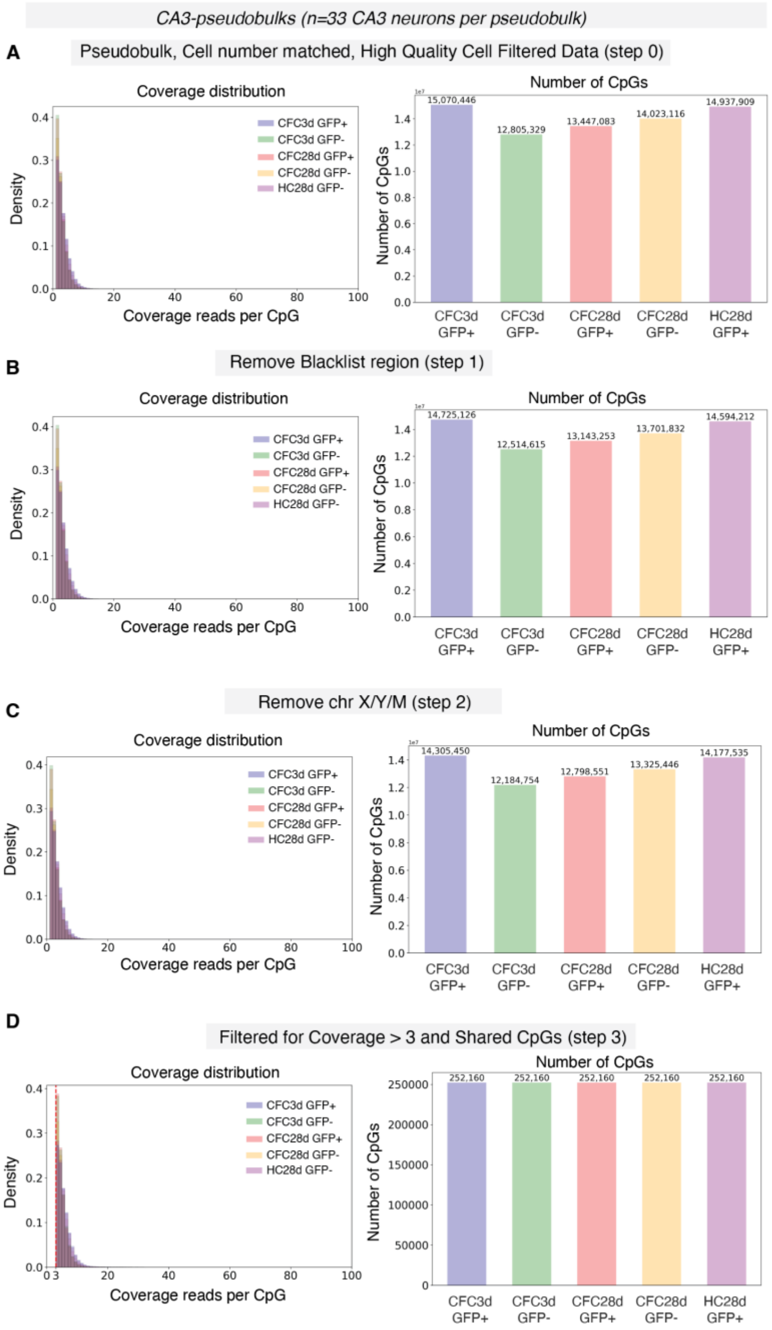


**Fig. S23. DNA methylation data processing at CpG resolution for CA3 neurons.**
(**A**) Pseudobulked raw data (step 0). (**B**) Filtered dataset after removal of CpGs overlapping with blacklist regions (step 1). (**C**) Filtered dataset after removal of CpGs belonging to chromosomes X, Y, and mitochondria (step 2). (**D**) Final filtered dataset containing only CpG sites shared across all four pseudobulked groups (step 3). Each stage displays three metrics: sequencing coverage distribution (left), and number of detected CpGs (right).

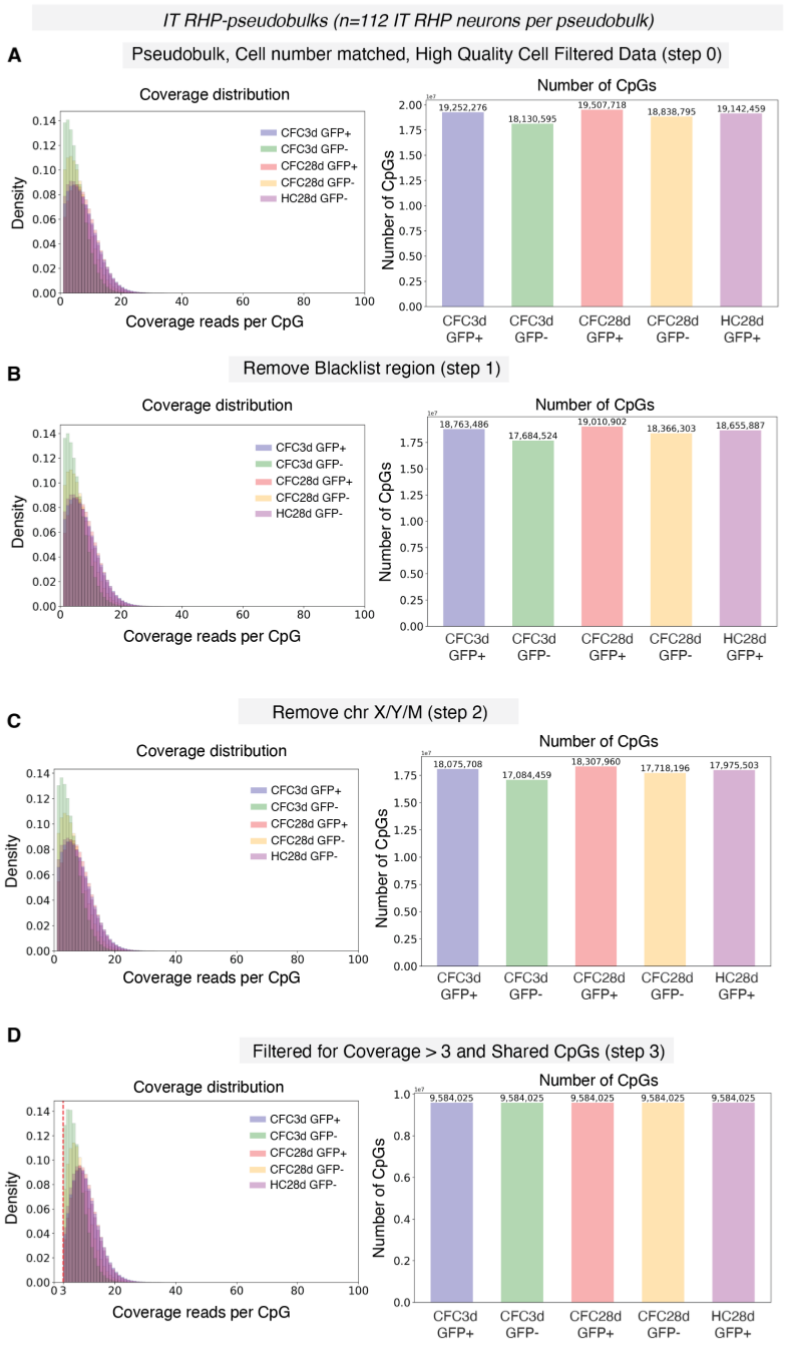


**Fig. S24. DNA methylation data processing at CpG resolution for IT neurons.**
(**A**) Pseudobulked raw data (step 0). (**B**) Filtered dataset after removal of CpGs overlapping with blacklist regions (step 1). (**C**) Filtered dataset after removal of CpGs belonging to chromosomes X, Y, and mitochondria (step 2). (**D**) Final filtered dataset containing only CpG sites shared across all four pseudobulked groups (step 3). Each stage displays three metrics: sequencing coverage distribution (left), and number of detected CpGs (right).

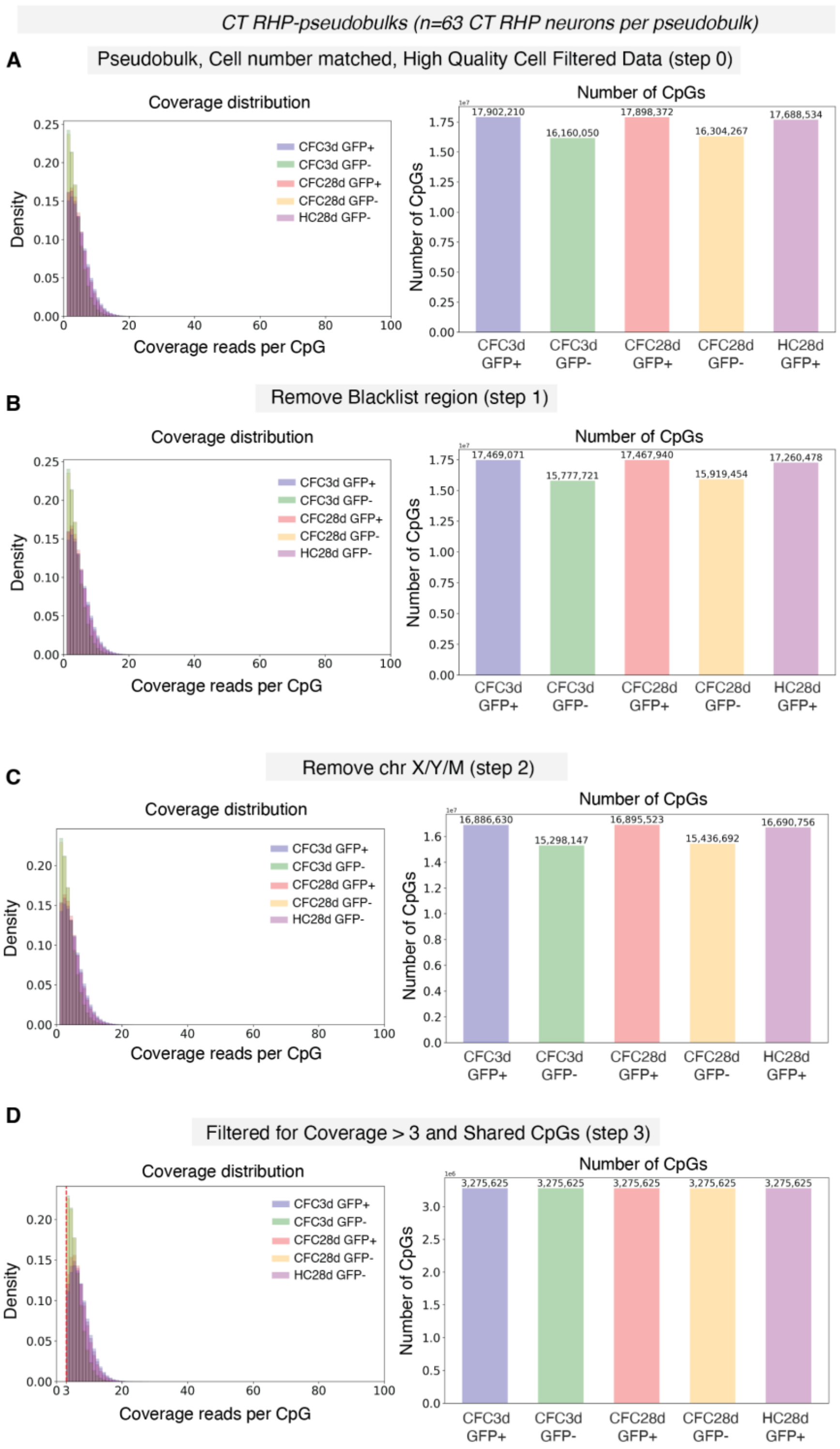


**Fig. S25. DNA methylation data processing at CpG resolution for CT neurons.**
(**A**) Pseudobulked raw data (step 0). (**B**) Filtered dataset after removal of CpGs overlapping with blacklist regions (step 1). (**C**) Filtered dataset after removal of CpGs belonging to chromosomes X, Y, and mitochondria (step 2). (**D**) Final filtered dataset containing only CpG sites shared across all four pseudobulked groups (step 3). Each stage displays three metrics: sequencing coverage distribution (left), and number of detected CpGs (right).

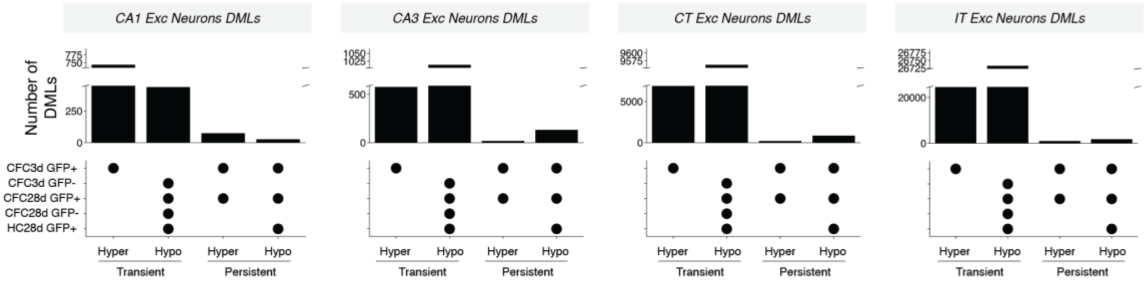


**Fig. S26. Excitatory subtype-specific differentially methylated CpG loci (DMLs) identified in CA1, CA3, CT, and IT accounting for 3 days and 28 days post-training in TRAPed and nonTRAPed conditions.**

Upset plots show the number of DMLs identified in each excitatory subtype for transient gained DMLs, persistently gained DMLs, transient lost DMLs, and persistent lost DMLs.

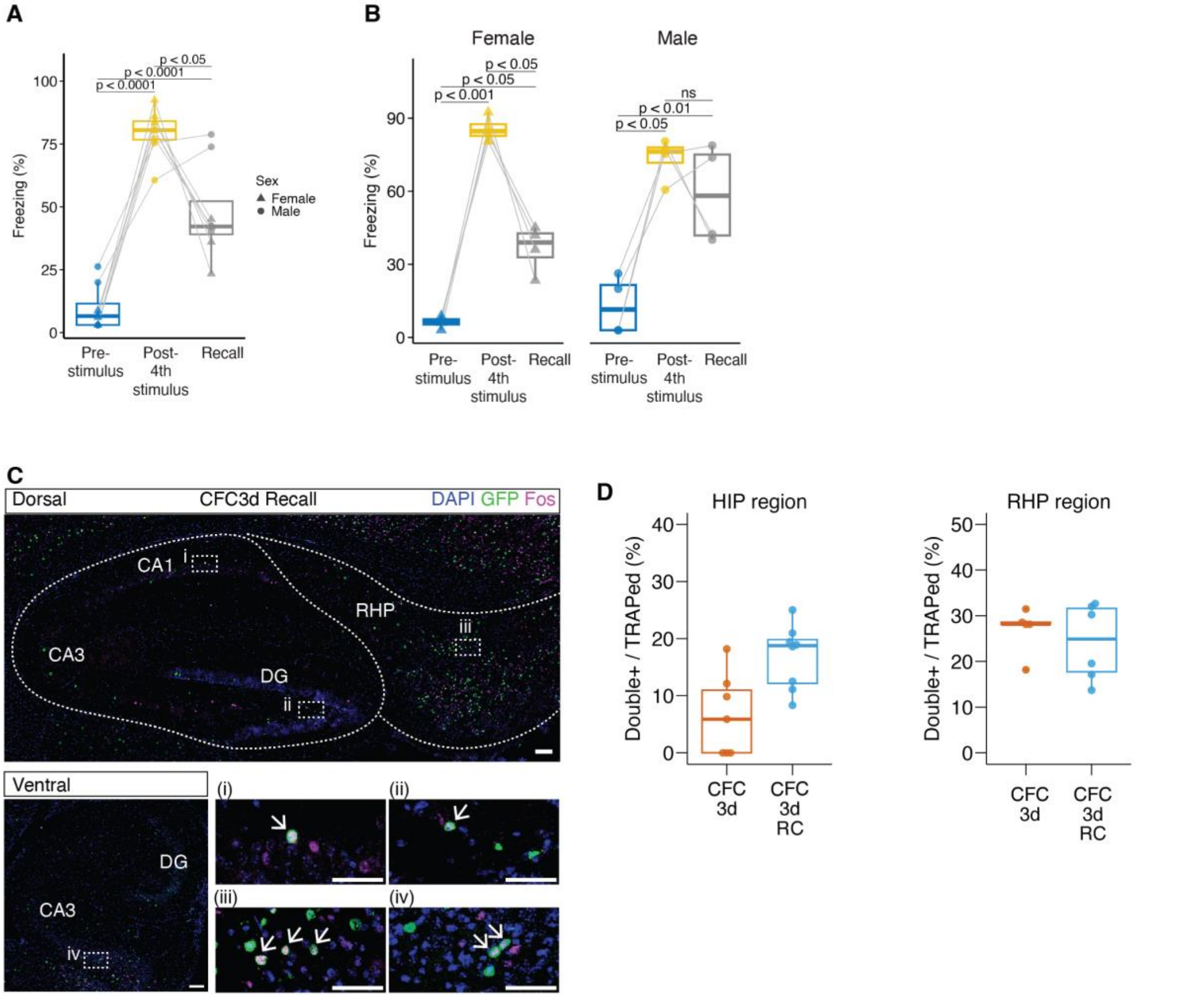


**Fig. S27. Behavioral analysis of TRAP2;INTACT transgenic mice upon contextual fear memory recall.**

**(A and B)** Quantification of freezing behavior (corresponding to Fig. 4) with recall testing at 3 days after CFC. Data are presented as **(A)** combined analysis and **(B)** segregated by sex. **(A)** One-way ANOVA with Holm–Sidak post hoc test, CFC +3 days + Recall (designated as CFC3d RC): F = 51.8, P =1.43 × $10^{-4}$. **(B)** One-way ANOVA with Holm–Sidak post hoc test, female CFC + 3 days + Recall: F = 116.4, P = 3.18× $10^{-5}$; male CFC + 3 days + Recall: F = 21.1, P = 0.02. Pvalues shown in the plot represent pairwise comparisons between measurements using two-tailed paired t-tests with Holm–Sidak post hoc correction. n = 8 per group; 4 males, 4 females. (**C**) Example confocal images of TRAPed cells that are TRAPed+Fos+ (Double+) in the dorsal hippocampal and retrohippocampal regions (top panel) and ventral hippocampal region in CFC + 3 days + Recall. (i-iv) show higher magnification views of the HIP and RHP regions. Arrows in (i-iv) indicate the TRAPed+Fos+ cells. (**D**) Quantification of percentage of TRAPed+ and Fos+ cells in hippocampal region (HIP, left panel) and retrohippocampal region (RHP, right panel), respectively. Scale bars: 100 µm; 50 µm (magnified views, i-iv). Analysis included n=2 mice per group, with each dot representing one slice.

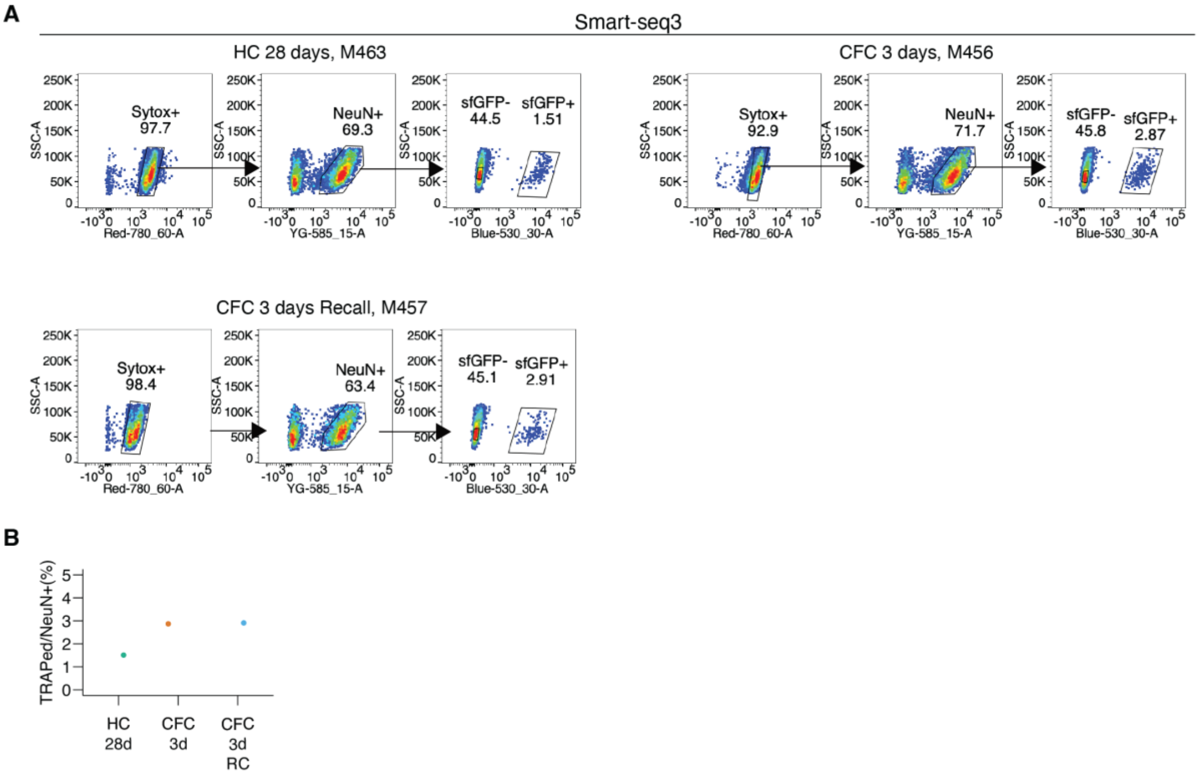


**Fig. S28. Representative flow cytometry plots and metrics for neuron sorting conducted for single nucleus Smart-seq3 experiments at 3 days post-training with and without context only recall.**

**(A)** Flow cytometry analysis of hippocampal formation neurons under different behavioral conditions. **(B)** Scatter plot showing the percentage of TRAPed cells among NeuN+ neurons for different experimental groups.

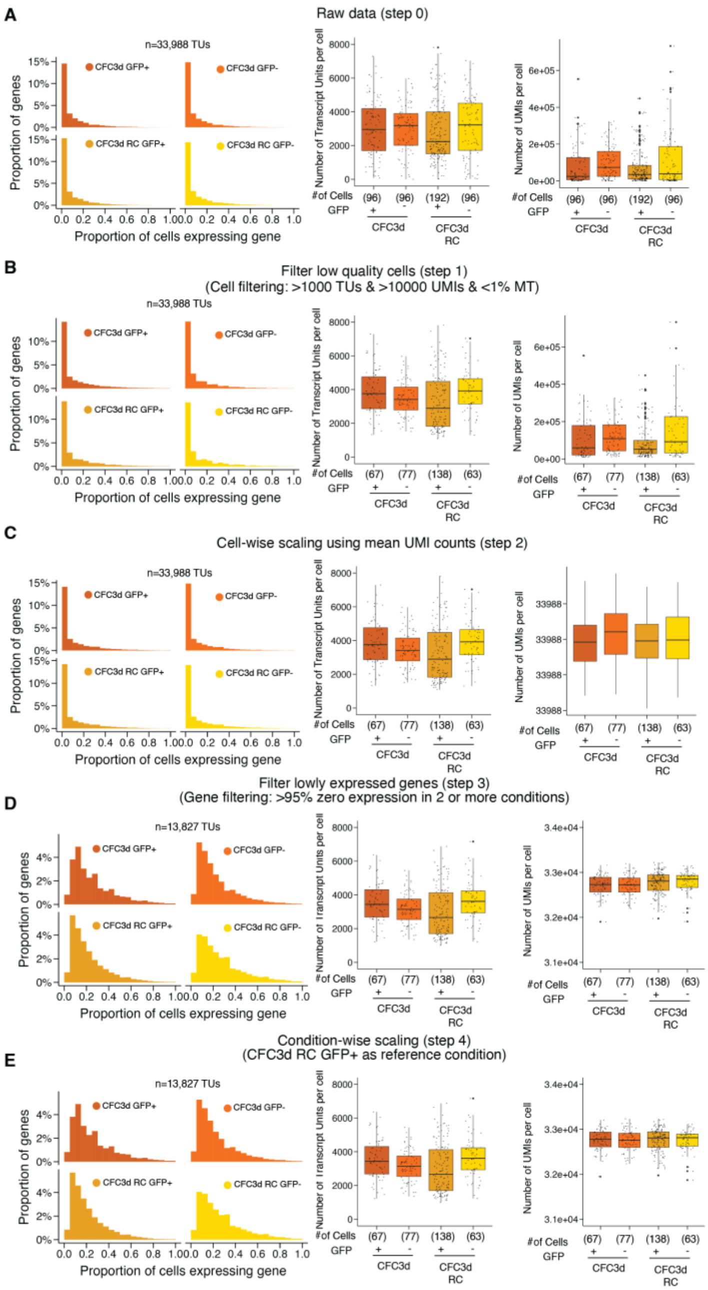


**Fig. S29: Quality control metrics to ensure rigorous and reproducible analysis of Smart-seq3 experiments.**

**(A)** Raw data (step 0). **(B)** data after filtering low-quality cells (step 1). **(C)** data after cell-wise scaling using mean UMI counts (step 2). **(D)** data after removing lowly expressed genes across cells (step 3). **(E)** data after condition-wise scaling using CFC3d RC GFP+ as reference condition (step 4). data after removing cells containing genes with more than 70% zero counts (step 3). Left panel: Distribution of gene expression frequencies across all detected genes (i.e., TUs). Each panel shows the proportion of genes (y-axis) expressed in a given proportion of cells (x-axis). Middle panel: Boxplots showing the number of TUs per cell for each condition. Right panel: Boxplots showing the number of UMIs (A to B) or normalized UMIs (C to E) per cell in each group.

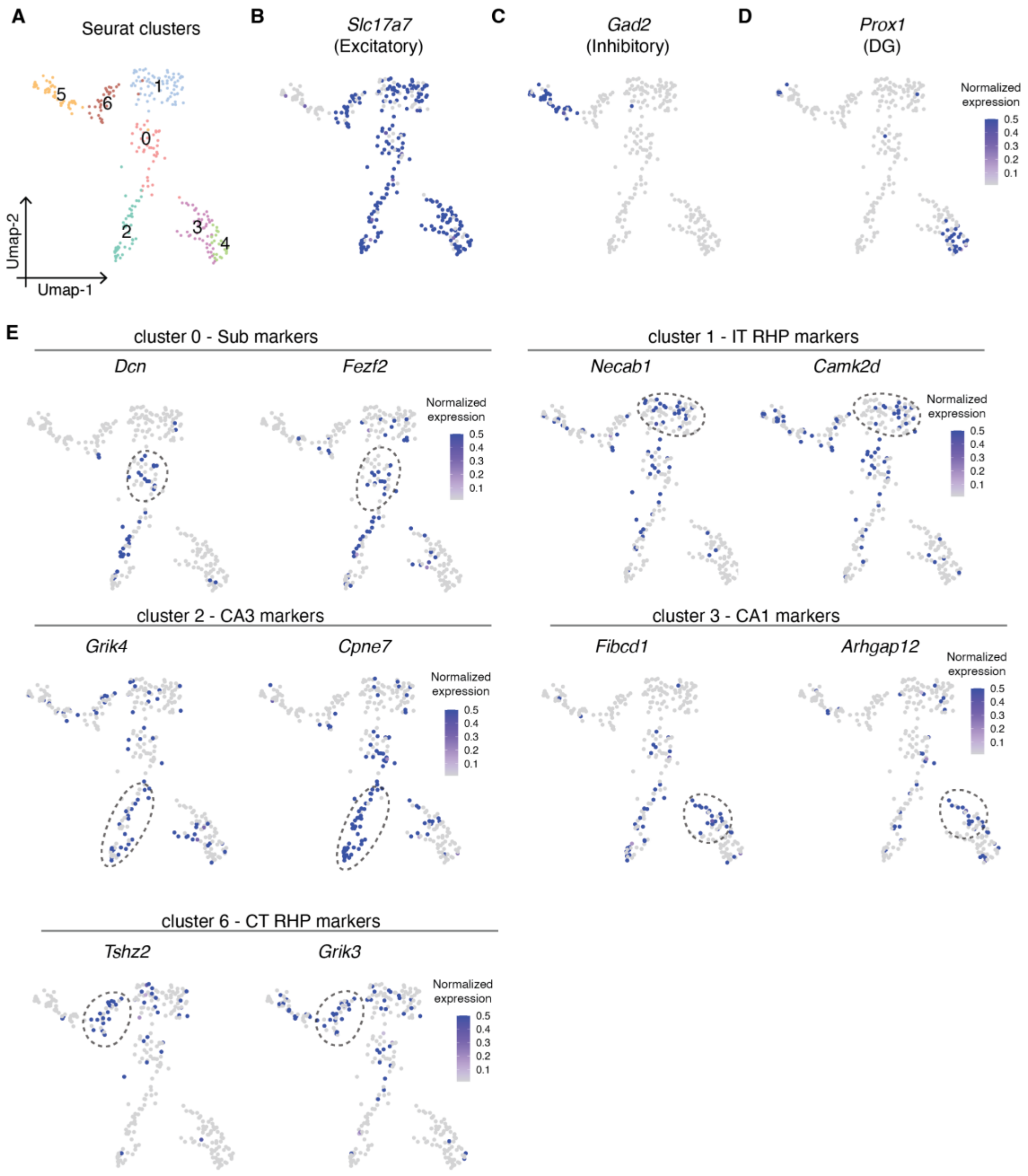


**Fig. S30. Annotation of neural subtypes in Smart-seq3 data via integration with Allen Brain Atlas Smart-seq2 gene expression data.**

**(A)** UMAP visualization of Seurat clusters. **(B-D)** Expression patterns of key marker genes mapped onto UMAP coordinates: *Slc17a7* (excitatory neurons) in (B), *Gad2* (inhibitory neurons) in (C) and *Prox1* (DG neurons) in (D). Color scale indicates normalized expression levels. **(E)** Expression patterns of representative marker genes for excitatory neuron subtypes based on Allen Brain Atlas data (*41*).

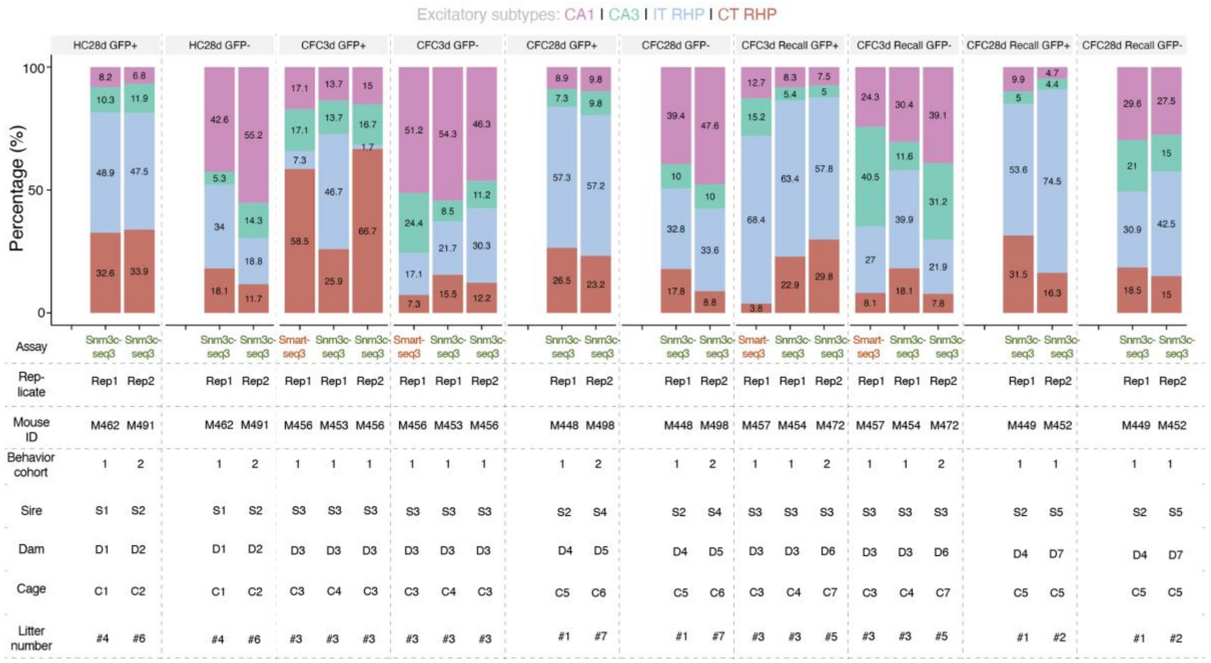


**Fig. S31. Quantification of excitatory neuronal subtype composition across behavioral conditions and assays demonstrating strong reproducibility of snm3C-seq and Smart-seq data between independent animal replicates.**

Bar plots showing relative percentages of excitatory neuronal subtypes (CA1, CA3, IT RHP, CT RHP) across indicated experimental groups (HC28d, CFC3d, CFC28d, CFC3d Recall, CFC28d, CFC28d Recall). Detailed metadata including experimental assay, mouse replicate, behavior cohort, animal sire and dam identifiers, cage numbers, and litter numbers are provided. The subiculum subtype was excluded due to insufficient cell numbers in the snm3C-seq assay. See **Table S2** for additional metadata details.

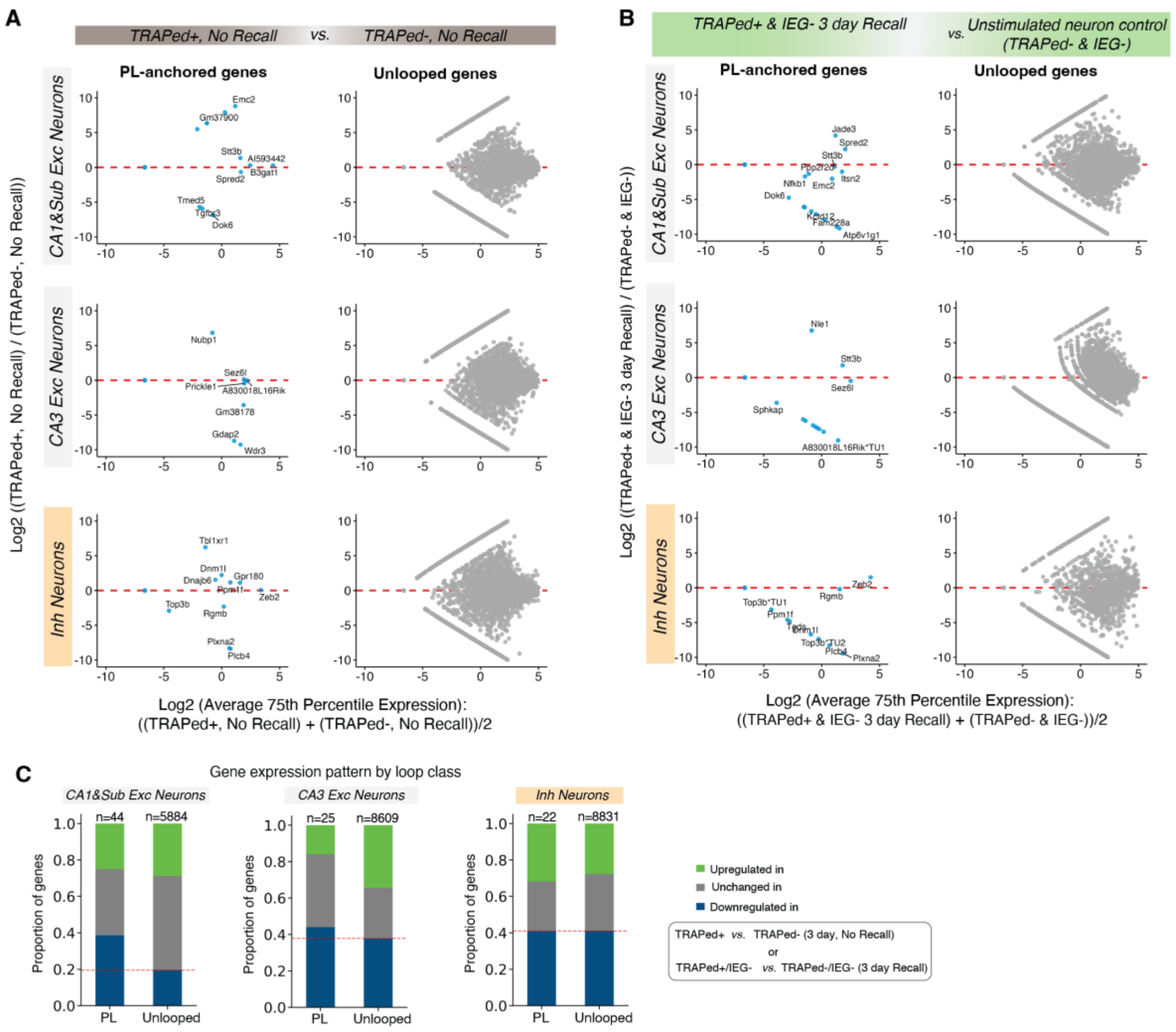


**Fig. S32. Gene expression pattern at PL loop-anchored genes in TRAPed+ neurons.**
**(A)** MA plots comparing TRAPed+, No Recall neurons to TRAPed-, No Recall neurons for PL loops and unlooped genes. **(B)** MA plots comparing TRAPed+ & Recall IEG- neurons to TRAPed- & Recall IEG- neurons for PL loops and unlooped genes. **(C)** Stacked bars show the fraction of genes that are downregulated (blue), upregulated (green), or unchanged (grey) within PL-anchored and unlooped gene sets for each neuronal subtype. Red lines indicate down-regulated ratio of the unlooped the controls. X-axis: log2 average 75th-percentile across the two conditions; Y-axis: log2 fold change between conditions. See also **Fig. 4** and **Table S14**.

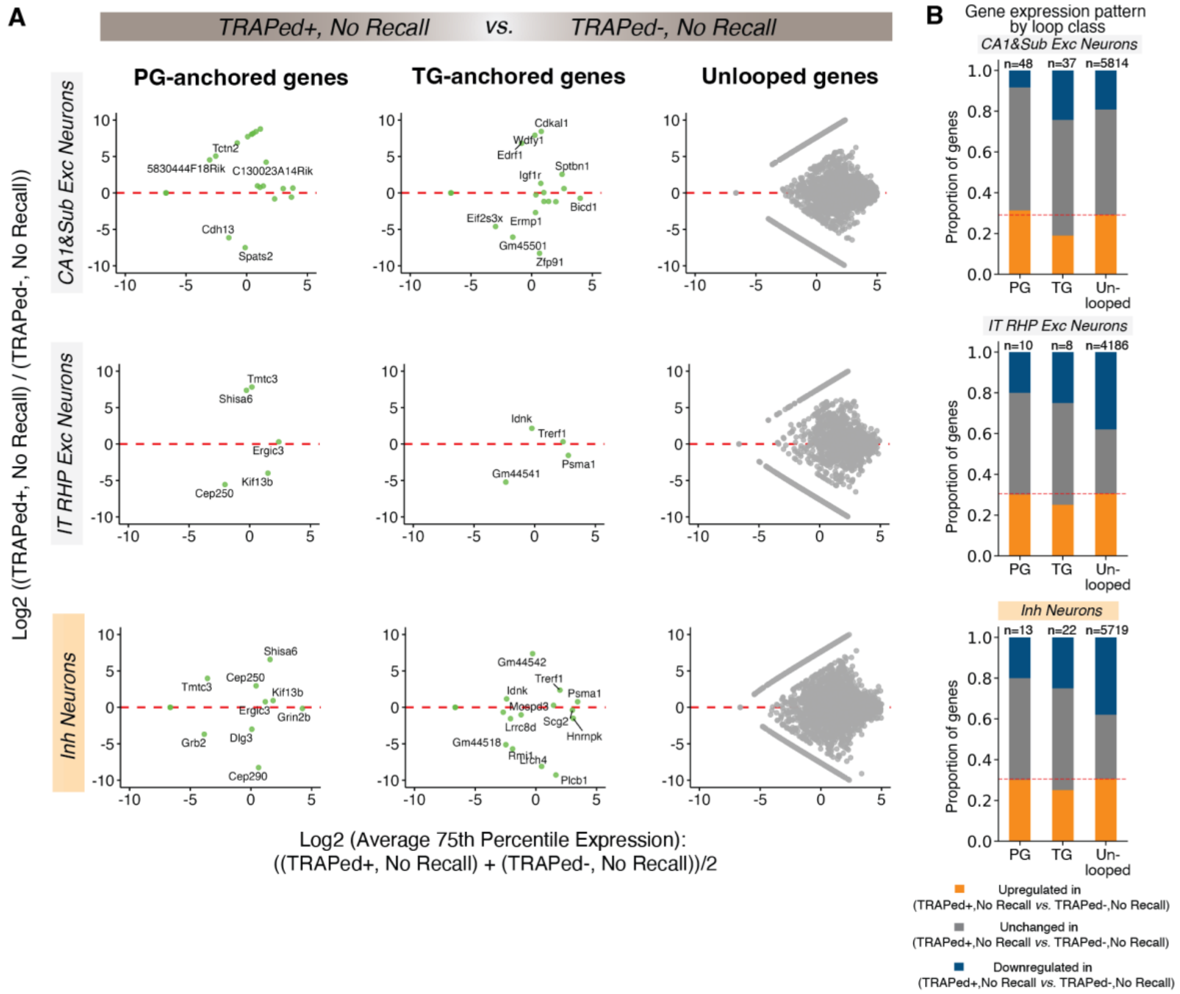


**Fig. S33. Gene expression pattern at TG or PG loop-anchored genes in TRAPed+ neurons.** **(A)** MA plots comparing TRAPed+, No Recall neurons to TRAPed-, No Recall neurons for PG-anchored, TG-anchored and unlooped genes. **(B)** Stacked bars show the fraction of genes that are downregulated (blue), upregulated (orange), or unchanged (grey) within PG-, TG-anchored and unlooped gene sets for each neuronal subtype. Red lines indicate down-regulated ratio of the unlooped the controls. X-axis: log2 average 75th-percentile across the two conditions; Y-axis: log2 fold change between conditions. See also **Fig. 5** and **Table S15**.

### Supplementary Tables

**Table S1.**
Detailed summary of all data generated in this study.

**Table S2.**
Detailed information of mice used in this study.

**Table S3.**
Barcodes and primers used in snm3C-seq3 and Smart-seq3.

**Table S4.**
Detailed mapping summary and neuronal annotation of single cells processed in snm3C-seq3 and Smart-seq3.

**Table S5.**
Number of different cell types by neuronal subtype and condition used for loop calling in Figure 1 in all neurons.

**Table S6.**
Number of matched neuronal subtypes representing a balanced excitatory neuron pseudobulk snm3C-seq analysis across condition used for loop calling in Figure 2.

**Table S7.**
Number of matched subtype numbers across conditions for loop calling on each individual CA1, CA3, IT and CT neuronal subtype for loop calling in Figures 3-5.

**Table S8.**
Resource of chromatin loop classes identified in pseudobulk snm3C-seq from all neurons (linked to Table S5, Figure 1).

**Table S9.**
Resource of chromatin loop classes identified in pseudobulk snm3C-seq from a merge of excitatory neurons and in each individual neuronal subtype (linked to Table S5-S7, Figure 2-5).

**Table S10.**
Number of activity-dependent chromatin loops called in each individual neuronal subtype.

**Table S11.**
Genes anchored at different loop classes in individual neuronal subtypes.

**Table S12.**
Gene ontology analysis of different loop classes in individual neuronal subtypes.

**Table S13.**
Persistent and transient hyper- and hypo-DMLs identified in individual neuronal subtypes.

**Table S14.**
List of downregulated genes anchoring persistently lost chromatin loops during fear memory recall.

**Table S15.**
List of upregulated genes anchoring persistently gained chromatin loops during fear memory recall.

**Table S16.**
List of packages used in the conda environments for processing snm3C-seq3 and Smart-seq3.

### Materials and Methods

Some of the methods described below have been detailed in our own previous manuscripts (*9, 12, 48-50, 52, 84-103*). To ensure reproducibility and clarity, we state the same or near similar methodological steps below. All data produced in this study is provided freely via GEO and linked detailed in **Table S1**.

#### *Mice breeding and housing*

All experimental procedures involving mice were performed in accordance with and approval from the Institutional Animal Care and Use Committee of University of Pennsylvania. We obtained TRAP2 mice ($Fos^{2A\text{-}iCreER}$, strain #030323) and $R26^{Sun1\text{-}sfGFP/+}$ (CAG-Sun1/sfGFP, strain #030952) from the Jackson Laboratory **(Fig. S1A)**. TRAP2 mice express both Fos and iCreERT2 under transient neural activity. $R26^{Sun1\text{-}sfGFP/+}$ mice express Sun1-sfGFP following the removal of a Stop cassette by iCreERT2 recombinase in the presence of 4-hydroxytamoxifen (4-OHT). We then crossed TRAP2 mice to CAG-Sun1/sfGFP to obtain double homozygous (TRAP2; CAG-Sun1/sfGFP) offspring **(Fig. S1A)**. We fed the mice a standard diet and housed them under a 12-hour light-dark cycle. We confirmed the genotype as described below and subsequently used them in behavior experiments as described below. Male and female mice aged 10 to 14 weeks were used at the time of brain collection. Animal experiment metadata are detailed in **Table S2**.

#### *Mice genotyping*

We collected tissue by ear punch from P14 pups and incubated at ~95-98°C in 50 µL of denaturing solution (25 mM NaOH, 0.2 mM EDTA) for 1 hour. We neutralized the solutions with 40 mM Tris-HCl pH 4.0 and purified the genomic DNA using a Zymo DNA Clean and Concentrate kit (Zymo, cat# D4014) following the manufacturer's protocol. Finally, we performed genotyping PCR using KAPA2G polymerase (Roche Diagnostics, cat# 07960891001) according to the manufacturer's instructions with the following primer sets:

For TRAP2 genotyping (WT Band = 357 bp; Mutant Band = 232 bp):

- WT forward: 5'-GTCCGGTTCCTTCTATGCAG-3'
- Mutant forward: 5'-CCTTGCAAAAGTATTACATCACG-3'
- Common Reverse: 5'-GAACCTTCGAGGGAAGACG-3'

For CAG-Sun1/sfGFP genotyping (WT Band = 557 bp; Mutant Band = 300 bp):

- Common forward: 5'-GCACTTGCTCTCCCAAAGTC-3'
- WT reverse: 5'-CATAGTCTAACTCGCGACACTG-3'
- Mutant reverse: 5'-GTTATGTAACGCGGAACTCC-3'

#### *Contextual fear conditioning (CFC)*

Starting three days prior to being placed into the fear chamber, we handled each mouse for 2 minutes before returning them to their cages each day. On the training day, we injected the mice subcutaneously with 4-OHT (40 mg $kg^{-1}$, Hello Bio, cat# HB6040) to initiate the temporal window of iCreERT2-mediated recombination, after which each mouse was placed in an individual conditioning chamber for a 3-minute acclimation period. Following acclimation, mice received four independent foot shocks (2 seconds duration, 0.75 mA intensity) delivered at 60-second intervals. After the administration of the final foot shock, mice remained undisturbed in the conditioning chamber for 2 minutes before being returned to their home cages **(Fig. S1B)**. Freezing behavior that was defined as no movement except for respiration, was determined before and after foot shock administration and scored by VideoFreeze software (Med Associates) **(Fig. S1C-D, S27A-B)**. To assess context-dependent learning, we placed the mice back into the same testing boxes 3 days or 28 days post-CFC for a total of 3 minutes without any shock, and again measured the total time spent freezing. We euthanized the mice at three different timepoints: 3 days or 28 days post-CFC for the no-recall groups (designated as CFC3d and CFC28d), or 3 hours following the recall test for the recall group (designated as CFC3d RC and CFC28d RC). Home cage control mice received the same 4-OHT treatment and were euthanized 28 days later but remained in their home cages throughout the experiment without undergoing behavioral testing (HC28d) **(Fig. S1B)**.

***Immunohistochemistry and imaging***

We anesthetized the mice with isoflurane and transcardially perfused with 10% formalin (Sigma, Cat# HT501128). Following perfusion, we removed the whole brain and placed it in 10% formalin overnight at 4°C. Following fixation, we preserved the individual brains for 48 hours in 30% sucrose at 4°C and then froze the brains at -20°C in optimal cutting temperature (OCT) compound in a cryopreservation mold. Once fully frozen, we removed the blocks from the molds and sectioned the brains using a Microm HM 550 cryostat, collecting 10 µm slices on Superfrost Plus Microscope slides. We then washed the brain sections in DPBS for 10 minutes to remove OCT compound and blocked for 1 hour at room temperature in 10% donkey serum (Jackson ImmunoResearch, cat# 017-000-121) containing 0.3% TritonX-100. Primary antibodies were diluted in 5% donkey serum containing 0.2% TritonX-100 and incubated overnight at 4°C. We used the following primary antibodies: rabbit anti-Fos (1:500, Synaptic Systems, cat# 226008) and goat anti-GFP (1:500, Novus Biologicals, cat #NB100-1770SS). After primary antibody incubation, we washed the sections three times in DPBS for 10 minutes each and diluted the secondary antibodies in DPBS and applied for 1 hour at room temperature in the dark, followed by three 10-minute washes in DPBS. We used the following secondary antibodies: Donkey anti-Rabbit IgG Alexa Fluor Plus 647 (1:1000, Invitrogen, cat# A32795), Donkey anti-Goat IgG (H+L) Alexa Fluor 488 (1:1000, Invitrogen, cat# A11055), and DAPI (1:1000, Invitrogen, cat# 62248). Finally, we mounted the sections with Fluoromount-GT (Electron Microscopy Sciences, cat# 17984-25) and imaged the slides with Zeiss LSM 880 Confocal microscopy with a 20x objective. Images were processed and quantified using ImageJ2 (Fiji, version 2.9.0/1.53t) **(Fig. S1E-F, S27C-D)**.

***Nuclei isolation***

We homogenized snap-frozen mouse brain tissue in a dounce homogenizer with 30 strokes of a loose pestle in 10 mL of Homogenization Buffer containing 320 mM sucrose (ThermoFisher, cat# BP2201), 5 mM $CaCl_2$ (ThermoFisher, cat# BP510-250), 10 mM Tris-HCl pH 8.0 (ThermoFisher, cat# 15568025), 3 mM $MgAc_2$ (ThermoFisher, cat# 63052-100ML), 0.1% (v/v) Triton X-100 (Sigma-Aldrich, cat# 93443-100ML), 0.1 mM EDTA (Sigma-Aldrich, cat# 15575020), 1 mM DTT (Sigma-Aldrich, cat# 10197777001). We then added another 12 mL of Homogenization Buffer and layered the homogenization solution over a 14 mL Sucrose Cushion (1.8 M sucrose, 10 mM Tris-HCl pH 8.0, 3 mM $MgAc_2$, 1mM DTT) in a Thinwall Polypropylene Tube (Beckman Coulter, cat# 326823). We ultra-centrifuged the samples at 25,700 rpm (Rotor, SW32Ti, cat# 369650) for 2 hours at 4°C. We aspirated the supernatant and washed the nuclei pellet with 1% Bovine serum albumin (BSA) (GeminiBio, cat# 700-100P-020) in Dulbecco's Phosphate-Buffered Saline (DPBS, cat# 14190-144) and centrifuged the samples at 800 × g for 5 minutes at 4°C and finally, stored at 4°C until further processing **(Fig. S2)**.

For Smart-seq3 experiments, we reduced the ultra-centrifuge time to 1 hour to minimize RNA degradation. We added RNase inhibitor (Takara, cat# 2313B) to the Homogenization Buffer and Sucrose Cushion at a working concentration of 0.04 U/µL. We also added Protease Inhibitor cocktail (Roche, cat# 11873580001, one tablet per 50 mL) into the Homogenization Buffer, Sucrose Cushion, and 1% BSA in DPBS.

***Nuclei crosslinking and target nuclei population sorting for snm3C-seq3***

For snm3C-seq3 experiments, we first crosslinked nuclei with DPBS-diluted methanol-free 2% formaldehyde (ThermoFisher, cat# 28906) for 10 minutes at room temperature with slow rotation (10 rpm) and then we quenched the fixation reaction by adding glycine (Sigma-Aldrich, cat# 50046) to a final concentration of 200 mM for 5 minutes with slow rotation at room temperature followed by a 5 minute incubation on ice. We pelleted the nuclei at 800 × g for 5 minutes and resuspended in 1% BSA in DPBS. We then repeated the washing step once more for a total of two washes. We resuspended nuclei in 1% BSA in DPBS and incubated with anti-NeuN-PE (1:1000, Sigma-Aldrich, cat# FCMAB317PE) for 1 hour at 4°C with slow rotation (10 rpm). Before fluorescence-activated nuclei sorting (FANS), we stained the nuclei with DAPI (1:1000) or 1:1000 SyTox (1:1000, Invitrogen, cat# S34859) and collected the sorted nuclei in 1% BSA in DPBS and centrifuged at 1000 × g for 5 minutes at 4°C for further library preparation (see below). FANS was performed at the Children's Hospital of Philadelphia Flow Cytometry Core Laboratory using a Cytek Aurora CS Cell Sorter, aBD FACSAria II cell sorter, or a BD FACSAria Fusion sorter **(Fig. S2, S28)**.

***snm3C-seq3 library preparation and sequencing***

We prepared snm3C-seq3 libraries using previous methods with modifications (*33, 34*). For nuclei conditioning, we gently resuspended the sorted nuclei population pellets in 50 µL of 0.5% sodium dodecyl sulfate solution (SDS) (Sigma-Aldrich, cat# 71736), incubated at 62°C for 10 minutes. We then added 145 µL water and 25 µL 10% Triton X-100 to quench the SDS and incubated at 37°C for 15 minutes. To ensure a consistent number of nuclei for the *in situ* 3C reactions, we divided the post-quenching mixture into multiple reactions. Each reaction contained between 200 and 800 nuclei, to allow for relatively tight control over cell number variation. This division was based on the total count of sorted nuclei. Each restriction enzymatic digestion reaction contained approximately 400-800 nuclei with 27 µL NlaIII (10,000 U/mL, NEB, cat# R0125L), 10 µL MboI (25,000 U/mL, NEB, cat# R0147M), and 14.2 µL 10X Cut Smart Buffer (NEB, cat# B6004S). We incubated the reactions at 37°C for 1 hour and then inactivated the restriction enzymes at 65°C for 20 minutes and cooled down at room temperature for 5 minutes. We performed the ligation by adding 900 uL Ligation Master Mix (669 µL water, 120 µL 10X NEB T4 DNA ligase buffer (NEB, cat# B0202), 100 µL 10% Triton X-100, 6 µL 20 mg/ml Recombinant Albumin (NEB, cat# B9200S) and 5 µL 400 U/ µl T4 DNA Ligase (NEB, cat# M0202)) to each tube, followed by slow rotation at room temperature for 4 hour and centrifugation at 1000 × g for 5 minutes. We then resuspended the nuclei in 1% BSA in DPBS with 1:1000 DAPI or SyTox. For single-nuclei sorting, we pre-filled a 384-well plate (Applied Biosystems, cat# 4483285) with 0.5 µL M-digestion buffer (Zymo, cat# D5021-9), 0.4 µL water and 0.1 µL Proteinase K (Zymo, cat# D3001-2-20) per well. After sorting, we centrifuged the 384-well plates at 2000 × g for 1 minutes at 4°C and incubated at 50°C for 20 minutes.

We performed the bisulfite conversion by adding 6 µL prepared CT Conversion Reagent (Zymo, cat# D5003-1) per well, incubating at 98°C for 8 minutes, 64°C for 3.5 hours, and holding at 4°C for up to 20 hours. For cleanup, we added 20 µL M-Binding buffer per well (Zymo, cat# D5049-3), mixed by pipetting and transferred into Zymo-Spin 384 Well Plate (Zymo, cat# C2012). We then centrifuged the plate at 4000 × g for 5 minutes at room temperature, added 100 µL M-Wash buffer (Zymo, cat# D5040-4) per well and centrifuged the plate again at 4000 × g for 5 minutes at room temperature. Then, we added 50 μL M-Desulphonation buffer (Zymo, cat# D5040-5) per well and incubated the plates for 15 minutes at room temperature. We repeated the washing step twice by adding 100 μL M-Wash buffer (Zymo, cat# D5040-4) per well and centrifuged the plates at 4000 × g for 5 minutes at room temperature. We finally eluted DNA from the plates with 4.4 μL M-Elution buffer (Zymo, cat# D5049-6) and combined the eluted DNA with 1.1 μL of 2.5 μM random barcoded primer (Integrated DNA Technologies, **Table S3**).

We denatured the plates at 98°C for 3 minutes in a thermocycler, followed by immediate cooling on ice to prevent rehybridization. We added 5 μL of random priming master-mix (1.025 μL 10X Blue Buffer (Enzymatics, cat# B0110), 0.025 μL Klenow Fragment (3'→5' exo–) (50 U/μL, Enzymatics, cat# P7010-HC-L), 0.5 μL Deoxynucleotide (dNTP) Solution Mix (10 mM, NEB, cat# N0447L) and 3.45 μL water) to each well. Then, we sealed the plates with Optical Adhesive Film (Thermo Scientific, cat# 4311971), quick-spun at 2000 × g for 1 minute at 4°C, and incubated in a thermocycler at 4°C for 5 minutes, 25°C for 5 minutes, 37°C for 60 minutes, and then held at 4°C. For inactivation of free primers and dNTPs, we added 1.5 μL master-mix to each well (0.2 μL 10X Blue Buffer, 0.1 μL Exonuclease I (20 U/μL, Enzymatics, cat# X8010L), 0.05 μL Shrimp Alkaline Phosphatase (rSAP) (1 U/μL, NEB, cat# M0371), and 1.15 μL water per reaction). We again sealed the plates, did a quick spin at 2000 × g for 1 minute at 4°C and incubated at 37°C for 30 minutes in a thermocycler and then held at 4°C.

To purify DNA products after random priming, we transferred contents from eight wells into one well in a new 96-well plate using a 12 multi-channel pipette per each 384-well plate. We added 73.6 μL (sample to beads ratio is 1:0.8) AMPure XP beads (Beckman Coulter, cat# A63881) to each well of the combined 96-well plate and thoroughly mixed by pipetting. Then, we incubated the plates for 8 minutes at room temperature and placed them on a magnet for 5 minutes. We carefully removed the supernatant and washed the beads twice by adding 180 μL of 80% ethanol to each well, incubating for 30 seconds per wash, and then allowed the beads to dry for 2 minutes. Each well received 10 μL Buffer EB (Qiagen, cat# 19086). We sealed the plates, vortexed, did a quick spin at 150 g for 5 seconds, incubated for 5 minutes and finally, placed on a magnet for 2 minutes. We further compressed the eluted samples into PCR tubes by transferring the 10 μL elutant from eight wells into one PCR tube. Each tube received 64 μL AMPure XP beads (sample to beads ratio is 1:0.8) per PCR tube and repeated beads purification step. We finally eluted the DNA by adding 10 μL of Buffer EB to each tube.

We performed adaptase reactions using xGen™ Adaptase™ Module 96rxn (Integrated DNA Technologies, cat# 10009826). First, we denatured the DNA samples in PCR tubes at 98°C for 3 minutes and placed them on ice to prevent rehybridization, followed by a quick spin. Each tube received 4.25 μL Low EDTA TE, 2 μL Buffer G1, 2 μL Buffer G2, 1.25 μL Buffer G3, 0.5 μL Enzyme G4, and 0.5 μL Enzyme G5. We then incubated the PCR tubes at 37°C for 30 minutes followed by 95°C for 2 minutes and finally held at 4°C. Each well received 30.3 μL of the indexed PCR master-mix, containing 25 μL 2X KAPA HiFi Hotstart® ReadyMix (Roche, cat# KK2602), 0.3 μL custom 100 μM P5 indexing primer (Integrated DNA Technologies, **Table S3**) and 5 μL custom 10 μM P7 indexing primer (Integrated DNA Technologies, **Table S3**). We conducted the PCR amplification with the following program: 95°C for 2 min, 15 cycles of 98°C for 30 s, 64°C for 15 s and 72°C for 30 s, followed by 72°C for 5 min, and held at 4°C. We then purified the amplified libraries with AMPure XP beads (sample to beads ratio is 1:0.8) and validated using Qubit™ dsDNA Quantification Assay Kits (Invitrogen, cat# Q32851) and Agilent 2100 Bioanalyzer. Finally, we pooled the libraries and sequenced on an Illumina NextSeq P2000 instrument using P4 XLEAP-SBS flow cells or a NovaSeq 6000 system using S2 flow cells with 150 bp paired-end mode.

***Smart-seq3 library preparation and sequencing***

For Smart-seq3 experiments, we first isolated the nuclei as described above. We omitted nuclei fixation and quenching steps and performed the antibody staining directly on unfixed nuclei. The antibody incubation time was reduced to 30 minutes, the centrifugation force was lowered to 300 × g, and RNase inhibitor (Takara, cat# 2313B) was added to the staining buffer (1% BSA in DPBS) at a working concentration of 0.4 U/μL.

We prepared the Smart-seq3 libraries using the previous methods (*104*) with the following modifications. Briefly, we sorted the single nuclei directly into a 96-well plate (ThermoFisher, cat# 4483348) with 3 μL lysis buffer per well, comprised of 0.5 U/μL recombinant RNase inhibitor (Takara, cat# 2313B), 0.1% (v/v) Triton X-100, 0.5 mM dNTPs (Thermo Scientific, #cat R0182), 0.5 μM Smart-seq3 oligo-dT primer (Integrated DNA Technologies, 5'-biotin-ACGAGCATCAGCAGCATACGAT30VN-3'), and 5% (w/v) PEG (Sigma, cat# 89510-250G-F). Immediately after sorting, we sealed the 96-well plates and centrifuged at 1000 × g for 1 minute, then promptly transferred to dry ice. We then incubated the plates at 72°C for 10 minutes and placed on ice. For reverse transcription (RT), we added 1 μL RT master-mix per well, containing 25 mM Tris-HCl pH 8.3 (Bioworld, cat# 40121113-2), 30 mM NaCl (ThermoFisher, cat# AM9760G), 1 mM GTP (ThermoFisher, cat# R1461), 2.5 mM MgCl2 (ThermoFisher, cat# AM9530G), 8 mM DTT (ThermoFisher, cat# 707265ML), 0.5 U/μL recombinant RNase inhibitor (Takara, cat# 2313B), 2 μM Smart-seq3 TSOs (Integrated DNA Technologies, 5'-biotin-AGAGACAGATTGCGCAATGNNNNNNNNrGrGrG-3'), and 2 U/μL Maxima H-minus reverse transcriptase (ThermoFisher, cat# EP0753). RT was conducted with the following program: 42°C for 90 min, 10 cycles of 50°C for 2 min and 42°C for 2 min, followed by 85°C for 5 minutes and held at 4°C. We performed PCR pre-amplification immediately after RT. We added 6 μL of PCR master-mix per well, containing 1× KAPA HiFi PCR buffer (Roche, containing 2 mM MgCl2, cat# KK2502), 0.02 U/μL DNA polymerase (Roche), 0.3 mM dNTPs, 0.5 μM Smartseq3 forward PCR primers (Integrated DNA Technologies, 5'-TCGTCGGCAGCGTCAGATGTGTATAAGAGACAGATTGCGCAATG-3'), and 0.1 μM Smartseq3 reverse PCR primer (Integrated DNA Technologies, 5'-ACGAGCATCAGCAGCATACGA-3'). Pre-amplification of cDNA was conducted with the following program: 98°C for 3 min, 25 cycles of 98°C for 20 seconds, 65°C for 30 seconds and 72°C for 4 minutes, followed by 72°C for 5 minutes and held at 4°C. After PCR pre-amplification, we purified each well with 0.6 X AMPure XP beads (sample to beads volume ratio of 1:0.6) and eluted the cDNA with 8 μL Buffer EB. We further diluted the cDNA with UltraPure water to an approximate range of 0.1 - 1 ng/uL and measured its concentration using a SpectraMax i3x plate reader with the Quant-iT PicoGreen dsDNA Assay Kit (ThermoFisher, cat# P7589). Only nuclei with original cDNA concentrations exceeding 4 ng/μL were considered successful amplifications and used for subsequent library construction. We next further diluted cDNA to 100 pg/μL per nuclei and performed tagmentation of cDNA in 2 μL master-mix, containing 10 mM Tris-HCl pH 7.5 (ThermoFisher, cat# 15567027), 5 mM $MgCl_2$, 5% (v/v) N,N-Dimethylformamide (Sigma-Aldrich, cat# D4551-250ML), 0.08 μL Amplicon Tagment Mix (Illumina, cat# FC-131-1024), 1 μL 100 pg/μL cDNA, and 0.42 μL water. We sealed the plates and incubated at 55°C for 10 minutes. 1 μL of 0.12% SDS (ThermoFisher, cat# AM9822) was then added to each well to release Tn5 from the DNA, followed by incubation at room temperature for 5 minutes. Each well received 2 μL 10-bp indexed Nextera index primers (1 uL each of 0.5 μM i5 and 0.5 μM i7 primers, **Table S3**), then 5 μL of tagmentation PCR master-mix, containing 1× Phusion HF Buffer (ThermoFisher, cat# F530S), 0.01 U/μL Phusion DNA polymerase, 0.2 mM dNTP, 0.05% Tween-20 (Sigma, cat#

P9416-50ML). Tagmentation PCR was conducted with the following program: 72°C for 3 min, 98°C for 3 min, 12 cycles of 98°C for 10 s, 55°C for 30 s and 72°C for 30 s, followed by 72°C for 5 min and holding at 4°C. After PCR, we combined all wells, purified with 0.6 X AMPure XP beads, and eluted with 40 µL Buffer EB. We validated the amplified libraries using Qubit™ dsDNA Quantification Assay Kits (Invitrogen, cat# Q32851) and Agilent 2100 Bioanalyzer. Finally, we pooled the libraries and sequenced on an Illumina NextSeq P1000/2000 instrument using P2 XLEAP-SBS flow cells with 150 bp paired-end mode.

***Installation of virtual environments for snm3C-seq3 data analysis***

We installed all required packages for processing snm3C-seq3 data in virtual environments using conda (v23.1.0) and mamba (v1.4.2). We created a conda virtual environment named "yap2" using the instruction given at https://hq-1.gitbook.io/mc/installation for demultiplexing FASTQ into single-cell FASTQ and running YAP pipeline to generate ALLC, Hi-C valid pairs and summary files for every single cell. We installed YAP (v1.6.9) and ALLCools (v1.1.0). Briefly, we created the environment using the command "mamba create" with the parameter -p denoting the location and python verion as 3.7. Then, we activated the environment using the command "mamba activate" and added the conda channels, such as defaults, bioconda, and conda-forge, using the command "conda config" with the parameter "--add channels". Since the random indices of our wells in V2 are slightly different, we changed the default index (<..>/yap2/lib/python3.7/site-packages/cemba_data/files/random_index_v2/random_index_v2.fa) using our random indices file. Then, we created a virtual environment called "allcools" using the instructions at https://lhqing.github.io/ALLCools/start/installation.html for generating MCDS files, clustering, processing ALLC files, generating pseudobulk ALLC files, pseudobulk bigwig files and DMR calls. We created our virtual environment and installed packages listed in **Table S16** as described.

***Running the YAP pipeline***

We first downloaded the snm3C-seq3 config.ini file from the GitHub page https://hq-1.gitbook.io/mc/prepare/prepare-mapping-config and modified it to process mouse genome mm10. We executed "yap demultiplex" to demultiplex the FASTQ data into single-cell FASTQ files, which generated Snakemake commands. We modified the Snakemake commands to optimize them for our HPC environment and subsequently executed them. After the completion of the YAP pipeline, we merged summary files across multiple runs and added additional columns, such as "celltype" (the sorted cell type, *e.g.*, cfc28d_gfppos means sorted GFP+ cells from mouse that went through contextual fear conditioning post 28 days), "plateinfo" (the human-readable 384-well plate identifier when preparing the library manually), "behavioral group" (behavioral condition, *e.g.*, CFC_28d means contextual fear conditioning post 28 days), "mouse ID" (mouse identifier) and "rep" (mouse replicate identifier). To keep the consistency in the naming convention and to improve readability of our methods, we named and added the columns, such as "TotalReads" as the sum of "R1InputReads" and "R2InputReads", "TotalMapped" as the sum of "R1UniqueMappedReads" and "R2UniqueMappedReads", "TotalDedupped" or "finalReads" as the sum of "R1DeduppedReads" and "R2DeduppedReads", "MappingRate" as the ratio of "TotalMapped" to "TotalReads", and "DeduppedRate" as the ratio of "TotalDedupped" to "TotalDedupped". For the 3C, we added a column named "cisTrans" as the sum of "CisShortContact", "CisLongContact", and "TransContact" and calculated the proportions of cis-short, cis-long and trans contacts respectively, such as "CisShortContactPro" as the ratio of "CisShortContact" to "cisTrans", "CisLongContactPro" as the ratio of "CisLongContact" to "cisTrans" and "TransContactPro" as the ratio of "TransContact" to "cisTrans". We also calculated cis-trans ratio "CisTransRatio" as the sum of "CisShortContact" and "CisLongContact" and then divided it by "TransContact".

***Mapping and quality control of snm3C-seq3 data***

We mapped snm3C-seq3 data using the YAP pipeline (v1.6.9) and ALLCools pipeline (v1.1.0) as previously described **(Table S4)** (*34*). Briefly, we demultiplexed FASTQ files into individual FASTQ files to cells using their corresponding random indices using the command "yap demultiplex" with the default parameters and the default configuration file (https://hq-1.gitbook.io/mc/prepare/prepare-mapping-config, mc-V2.mouse.mapping_config.ini). Then, we aligned the reads to the mm10 bismark reference genome using the reads-splitting strategy with bismark (v0.20). We first trimmed the reads with cutadapt (v2.10) and aligned them using bismark. The unmapped reads were trimmed again, split into 3 parts and aligned again with bismark. The total aligned reads from two mappings were merged, filtered, sorted and removed for duplicates using samtools (v1.6) and Picard (v3.2.0). For the Hi-C/3C contacts, split reads were merged to form valid pairs. For the methylome profiles, we generated ALLC files from the bam files using the function "bam-to-allc" with the allcools package.

Briefly, we removed low-quality cells (i.e., cells with mCCC fraction >0.03, mCH fraction >0.2, mCG fraction <0.5, mapping rate <0.5) as previously described **(Fig. S3)** (*34*). We also added a filter for chromatin contacts. We calculated cis/trans ratio as the sum of cis short contacts and cis long contacts and divided by trans contacts for every cell. Cells with a cis/trans ratio < 5 were filtered out. Additionally, cells were filtered based on the total number of deduplicated reads, which refers to reads remaining after the removal of PCR duplicates. Only unique read names from the deduplicated BAM file were counted. To pass quality control, cells required between 230,000 and 4,000,000 total deduplicated reads, referred to as "final reads". All these thresholds were applied such that all criteria must be met simultaneously for filtering. This approach excludes cells with exceptionally low or high values, which may represent technical artifacts or low-quality samples. With 6 filters, we removed 38.2% cells of poor-quality and proceeded with 61.8% cells of high-quality for the downstream analyses **(Fig. S3, Table S4)**.

***Clustering and annotation using DNA methylation***

We performed clustering of the cells based on the CpG and non-CpG DNA methylation at 100 kb resolution using established methods (*83*). Briefly, we generated an MCDS (Methylation Count Dataset) file, a specialized data used in ALLCools toolkit based on xarray.Dataset; https://lhqing.github.io/ALLCools/command_line/allcools_dataset.html) at 100 kb resolution using the reads that support CpG and non-CpG methylation using the command "allcools generate-dataset --regions chrom100k 100000 --quantifiers chrom100k count CGN,CHN". The parameters "regions" indicated the 100 kb non-overlapping bins genome-wide and "quantifiers" indicated the total reads and reads supporting for CpG and non-CpG methylation per 100 kb bin. We merged metadata containing the quality control metrics across cell to the MCDS file and calculated the average coverage of reads per 100 kb bin across cells.

*Filtering bins of low quality:* We loaded the MCDS file using the function "MCDS.open" and added the filtered metadata using the function "MCDS.add_cell_metadata". We first calculated the average of coverage across cells for every bin using the function "MCDS.add_feature_cov_mean" and plotted the histogram at 100 kb bin size **(Fig. S4)**. We then removed bins overlapping the chromosomes X, Y, and M using the function "MCDS.remove_chromosome" and ENCODE blacklist regions (https://github.com/Boyle-Lab/Blacklist/) using the function "MCDS.remove_black_list_region". We also removed low-quality and unusually high coverage bins because they might represent low-mappability regions and alignment artifacts using the function “MCDS.filter_feature_by_cov_mean”. Based on the distribution of average coverage, we determined minimum and maximum cutoff thresholds as follows: 500 and 1400 for 100 kb resolution, 250 and 800 for 50 kb resolution, and 120 and 400 for 25 kb resolution **(Fig. S4)**.

*Calculating methylated-cytosine fractions and cell-to-cell normalization:* With total counts and counts supporting methylation, we calculated methylation fractions independently for CpG and non-CpG methylation using the function “MCDS.add_mc_frac” and performed cell-to-cell normalization using beta-binomial distribution method as described before (*83*) by setting the parameter “normalize_per_cell” as TRUE. Internally, this function calculated the two shape parameters of the distribution $\alpha$ and $\beta$ using the sample mean ($m$) and variance ($v$) of the raw methylcytosine level (ratio of methylated-cytosine coverage to total cytosine coverage) as a prior estimation required for the beta-binomial distribution. Subsequently, we calculated normalized values using a methodology analogous to the counts per million reads (CPM) approach employed in single-cell RNA-seq analysis. We show the boxplots of methylated fractions before and after cell-to-cell normalizations at 100 kb resolution **(Fig. S5)**.

*Calculating highly variable features (HVFs):* Selection of highly variable features has been crucial in single-cell analysis. In snm3C-seq3, there is a non-linear correlation between methylated fraction, average coverage, and dispersion **(Fig. S6)**. Since removing the non-linear correlation is important before selecting HVFs, we removed the non-linear correlation independently for CpG and non-CpG methylation using a non-linear Support Vector Machine for Regression (SVR) using the function “MCDS.calculate_hvf_svr” based on the scanpy’s function “scanpy.pp.highly_variable_genes”. Using “n_top_feature” within the function “MCDS.calculate_hvf_svr”, we obtained HVFs independently for CpG and non-CpG methylation. We show the scatter plots before and after removing the correlations and HVFs at 100 kb resolution **(Fig S6)**.

*Principal component analysis (PCA):* We independently performed PCA for CpG and non-CpG methylation using the scanpy’s function “scanpy.tl.pca” and identified significant components using “significant_pc_test” function from ALLCools package. However, upon visual inspection of variance ratio plots for CpG and non-CpG methylation using the scanpy’s function “scanpy.pl.pca_variance_ratio” **(Fig. S7)**, we realized that the number of significant components was higher than what “significant_pc_test” gave. Hence, we changed the significant components to 25 for CpG and non-CpG methylation. Then, we concatenated the significant PCs of CpG and non-CpG methylation after normalizing for standard deviation.

*Choosing the Leiden resolution & embedding:* We built a k-nearest neighbors (KNN) graph, defined the clusters using the leiden algorithm and embedded the cells using tSNE and UMAP. Briefly, we obtained a KNN graph using the scanpy’s function “scanpy.pp.neighbors” after concatenating the significant dimensions from the PCA of CpG and non-CpG methylation. Then, we defined the clusters using the scanpy’s function “scanpy.tl.leiden”. Since the parameter “resolution” of the function “scanpy.tl.leiden” directly affected the leiden algorithm in defining the number of clusters, we tried 3 different resolution parameters, such as 0.25, 0.5, and 1 at 100 kb bin size **(Fig. S8)**. It was evident that increasing the resolution increased the number of clusters identified. Then, we embedded the cells with tSNE and UMAP using the functions “tsne” and "scanpy.tl.umap" from the packages "t-SNE" and "scanpy", respectively **(Fig. S9)**. Based on the non-CpG DNA methylation of the marker genes in Fig S10E, we preferred UMAP as our choice of embedding and concluded that 0.25 as the leiden's resolution that best fitted the clusters. To remove the randomness in defining the clustering, we performed consensus clustering with 100 rounds of leiden clustering using the function "ConsensusClustering" with the leiden resolution as 0.25 at 100 kb **(Fig. S9)**.

*Annotation based on the DNA methylation of the marker genes:* Based on the knowledge about the cell-types of hippocampal formation (*41*), we plotted the non-CpG DNA methylation of gene body +/-2 kb flanking regions of the known markers (*Prox1* and *Dock10* for DG neurons; *Gad1* and *Gad2* for Inh neurons; *Slc17a7* for Excitatory neurons) to inferred their expression. In general, higher non-CpG DNA methylation correlates with transcriptionally silenced genes, while lower non-CpG methylation is associated with transcriptionally active genes. Across various bin sizes **(Figs. S9-S12)**, our analysis consistently revealed 3 major neuron types, including DG neurons, Exc neurons and Inh neurons based on the non-CpG DNA methylation of markers genes. Finally, we directly compared the cluster annotation of clustering with different percentages of bins (i.e., 25%, 50% and 100% of bins) during HVF selection. Regardless of the percentages, the major neuron annotation among them remains largely same **(Fig. S10)**. To enhance cell type clustering precision, we also collected hippocampus neurons (NeuN+) 3 hours post-CFC (designated as CFC3h) from mice that received PBS injections. Therefore, we performed clustering with the following groups: CFC3h NeuN+, CFC3d GFP+, CFC3d GFP-, CFC28d GFP+, CFC28d GFP-, HC28d GFP+, and HC28d GFP- **(Fig. S3-S12)**.

***<u>Integration of snm3C-seq with Allen Smart-seq2 gene expression marker data to annotate neural subtypes</u>***

We integrated the snm3C-seq3 and public scRNA-seq datasets as previously described **(Fig. S11)** (*34*). We extracted cells belonging to the hippocampal formation region, based on the regional identity in Allen's dataset (*41*). We then filtered the cells based on the "subclass_label" annotation from Allen's dataset. Non-neuronal cells, including astrocytes, microglia, and oligodendrocytes (annotated as "Oligo", "Astro", "Micro-PVM"), were excluded. Additionally, we removed cells related to the isocortex, including those labeled as "L5 PT CTX", "L6b CTX", "L6 IT CTX", "L6 CT CTX", "L2/3 IT CTX", "L4/5 IT CTX", "Car3", "L4 RSP-ACA", "L5 IT CTX", and "L5/6 NP CTX". We normalized gene expression of the remaining cells by dividing each cell's total unique molecular identifier (UMI) count by the mean count across all cells, multiplying by a scale factor of 40,000, and then applying a natural log transformation.

We independently analyzed Leiden-based clustering results at 100kb bin sizes and integrated it with Allen's datasets. For mC cells, we used the posterior gene-body mCH level. We identified cluster-enriched genes (CEGs) within each cell subclass of Allen's datasets and within each cluster of mC data. We selected genes with mC variance exceeding 0.05 and expression variation surpassing 0.05 for further analysis. Given the inverse relationship between gene-body DNA methylation and gene expression, we inverted mC levels before integration. We trained a Truncated Singular Value Decomposition (TruncatedSVD) model on mC cells and applied the model to transform the merged RNA and mC dataset. To avoid dominance by the first few principal components, we normalized each component by its corresponding singular value, ensuring a balanced contribution in the embedding.

To identify anchors between mC and RNA cells, we z-score scaled the mC matrix and expression matrix of CEGs across cells and merged. We used Canonical Correlation Analysis (CCA) to find shared low-dimensional embeddings and identified five anchors using mutual nearest neighbors and Seurat-based scoring (*105*). We then employed Harmony algorithm to correct shared embeddings for batch effects (*106*). We constructed a KNN graph was using the Harmony embedding as the representation and performed clustering with the Leiden algorithm. To enhance the spatial representation of clusters, we applied PAGA (*107*) to infer the connectivity between the Leiden co-clusters, providing an initialization for the UMAP embedding. We generated a confusion matrix by calculating the overlap score between clusters from the original cluster designations of each dataset and their corresponding co-cluster assignments.

We conducted cell type annotation of snm3C-seq3 data using the overlap score of the confusion matrix and regional identity from Allen's reference dataset across various bin size clustering results. We used 100 kb size clustering results as final cell type annotation, with the following modifications. We merged minor inhibitory neuronal subtypes into a single Inh cluster. Additionally, we merged intratelencephalic neurons (IT) and corticothalamic (CT) neurons from the retrohippocampal region (RHP), including the entorhinal cortex (ENT), parasubiculum (PAR), postsubiculum (POST), presubiculum (PRE), subiculum (SUB), and prosubiculum (ProS) into one IT subtype (designated as IT RHP) and CT subtype (designated as CT RHP), respectively. At the major neuronal class level, we consolidated all excitatory neuronal subtypes into a single Exc neurons category. Due to dramatic cell type difference, we kept DG neurons separated from the rest of Exc neurons for chromatin loop calling and methylation analysis **(Fig. S12, S31)**.

***Interaction frequency heatmap generation and loop calling***

We generated Observed/Expected matrices from the pseudobulked data as detailed in **Tables S5-S7** using the 3C valid pairs obtained from snm3C-seq3 from individual neurons. We identified chromatin loops using our established computational pipelines with modifications **(Fig. S13-S15)** (*12, 50, 84, 88, 89, 96, 102*). Briefly, we assembled the contact maps chromosome-wise using the 3C contacts at single-cell resolution at 30 kb bin size after sorting the contacts into the upper-triangle using the "csort" function from the Cooler package with the parameters "-c1 2 -c2 6 -p1 3 -p2 7". Then, we developed a Hi-C contact distance-dependent method to scale the contact maps at the chromosome level to bring the single cells on the same scale and to remove the biases owing to the differences in read depth across cells. We read the contact maps stored as sparse matrices with the extension ".npz" using the function "sparse.load_npz" from scipy package. Then, we used a custom function to extract the union of all indices across cells. Then, we created a 2D array with rows corresponding to all indices and columns representing the single cells and added a pseudocount of 0.00001 to the 2D array. For a given contact distance, we extracted the rows and scaled using the geometric mean. The resulting contact probability maps across cells had a uniform distribution across cells **(Fig. S13)** and we stored the contact maps in the ".npz" file format.

We merged the single neurons to obtain pseudobulked data as described **(Tables S5-S7, Fig. S13-S15)** and proceeded to the downstream steps. For the loop calling, we first merged all neurons across five groups (CFC3d GFP+/-, CFC28d GFP+/-, and HC28d GFP+) into a pooled sample. We then removed bins corresponding to low mappability regions, balanced the maps using the Knight-Ruiz (KR) algorithm and implemented imputation from the scHiCluster package with a few modifications (*108, 109*) **(Fig. S13)**. The scHiCluster imputation involved transforming the data into sparse contact file format, imputation and a matrix balancing step using square root vanilla coverage normalization method. Since we already balanced our contact maps using KR balancing, we opted not to do a second balancing step. The imputed contacts were transformed by multiplying with a scalar (i.e., 1000) and used as the Observed (i.e., OBS) maps.

We converted the contact maps into sparse format contact matrices and implemented imputation using the function "impute_cell" from the package "scHiCluster" with the following parameters "logscale=False, rp=0.5, tol=0.01, window_size=500000000, step_size=10000000, output_dist=500000000" without the default balancing algorithm square root vanilla coverage (SQRTVC) post-imputation (*108*). Then, we multiplied the values with a scalar of value 1000 and used it as the Observed values and calculated the Hi-C Donut Expected (i.e., EXP) values. We used calculated Observed/Expected (i.e., OBS/EXP) for downstream analysis **(Fig. S13)**.

For loop calling in the pooled pseudobulk **(Table S5, Figure 1)**, we then computed the Expected (i.e., EXP) matrices using the Donut geometric filter, calculated Pvalues, and called loops using established in-house computational pipelines as previously described (*12, 50, 84, 88, 89, 96, 102*). We called loops using the pooled pseudobulk and applied 3 different combinations of Pvalues and minimum cluster sizes: v1 (permissive) with Pvalue 0.075 and minimum cluster size of 2, yielding 9,997 chromatin loops; v2 (intermediate) with Pvalue 0.075 and minimum cluster size of 3, resulting in 7,222 loops; and v3 (conservative) with Pvalue 0.05 and minimum cluster size of 3, identifying 6,171 loops. We used the v1 permissive loop clusters for loop classification (see next section). All the steps except imputation were done using established in-house computational pipelines as described previously (*12, 50, 84, 88, 89, 96, 102*).

***Thresholding paradigm for loop classification across conditions***

For each loop, we extracted the OBS and EXP values across pixels for a given sample and then calculated their averages as OBS_mean and EXP_mean. Then, we calculated the OBS/EXP as the ratio of OBS_mean to EXP_mean, resulting in an OBS/EXP (OE) value for that sample. We observed that loops with OE ratios $\leq$ 1.6 exhibited lower signal-to-noise ratios; therefore, we retained only loops with OE ratios greater than 1.7 for further analyses.

For the main cohort of our data (five conditions of: CFC3d GFP+/-, CFC28d GFP+/-, and HC28d GFP+), we defined activity-dependent chromatin loop categories as follows: *Transient Activity-Gained Loops*: OE $\geq$1.7 exclusively in CFC3d GFP+ and OE <1.6 in all other groups; *Persistent Activity-Gained Loops*: OE $\geq$1.7 at both CFC3d GFP+ and CFC28d GFP+, and OE <1.6 in CFC3d GFP-, CFC28d GFP- and HC28d GFP+; *Transient Activity-Lost Loops*: OE <1.6 exclusively in CFC3d GFP+ and OE $\geq$1.7 in all other groups; *Persistent Activity-Lost Loops*: OE <1.6 at both CFC3d GFP+ and CFC28d GFP+ and OE $\geq$1.7 in CFC3d GFP-, CFC28d GFP- and HC28d GFP+. We defined *Persistent Activity-Strengthened* loops by OE $\geq$ 1.7 in all groups and exhibiting at least a 1.25-fold higher OE ratio in CFC3d GFP+ *vs.* CFC3d GFP−, CFC28d GFP+ *vs.* CFC28d GFP−, and CFC28d GFP+ *vs.* HC28d GFP+. We defined *Persistent Activity-Weakened* loops by OE $\geq$ 1.7 in all groups and at least a 1.25-fold lower OE ratio in CFC3d GFP+ *vs.* CFC3d GFP−, CFC28d GFP+ *vs.* CFC28d GFP−, and CFC28d GFP+ *vs.* HC28d GFP+. We defined *Activity-Invariant Loops* as OE $\geq$1.7 across all groups and conditions, but not in *Persistent Activity-Strengthened* or *Persistent Activity-Weakened* **(Fig. S16-S21)**.

For cell type-specific analyses, we generated OBS/EXP pseudobulk contact maps and loop classifications separately for all cells merged together **(Table S5, Table S8, Figure 1),** each neuronal subtype (CA1, CA3, IT RHP, CT RHP, DG, and Inh), and a merged excitatory neuron pseudobulk balanced by equal cell numbers per subtype (Exc) **(Table S6-S7, Table S9-S10, Fig. S16-S21, Fig. 2-5)**. Loop classification and threshold criteria were applied similarly across different cell type populations.

***Classification of loops into promoter-promoter (PP) and promoter-enhancer (PE)***

For each loop, we checked upstream and downstream anchors independently for the presence of a promoter, non-coding enhancer, or UTRs-exons. Promoters were defined as upstream regions spanning the transcription start site (TSS) ±10 kb. Enhancers were defined from differentially enriched H3K27ac peaks in hippocampal tissue post-kainic acid (KA) treatment relative to unstimulated controls (GSE175952), identified using our established computational pipeline (*12,*

*50, 84, 88, 89, 96, 102, 110*). We merged H3k27ac peaks called within 400 bp and merged the peaks from both KA and unstimulated samples and identified KA-specific, unstimulated-specific and invariant peaks using our established computational pipeline. Post-KA-specific peaks were designated as KA-induced enhancers, while peaks lost after KA treatment were designated as KA-decommissioned enhancers. Enhancers were then restricted to regions not overlapping ±2 kb of a TSS, exons, and UTRs, and 10-kb windows were constructed to delineate enhancer regions.

To define loop anchored genes, we used transcription units (TUs) annotations to represent genes. Transcripts with TSSs located within 10 kb of each other were merged into a single TU using a custom script. For each TU, the most upstream TSS was designated as the TSS, and the most downstream transcription end site (TES) was designated as the TES. Anchors with both promoter and enhancer were called PE-anchors; anchors with just promoter, enhancer, or UTR/exon were classified as P-anchors, E-anchors, or UE-anchors, respectively; all others were designated N-anchors. Loops with upstream and downstream anchors such as PE-PE, PE-P, PE-E, P-PE, P-E, E-PE, and E-P were classified as promoter-enhancer (PE) loops. Loops with P-P anchors were classified as promoter-promoter (PP) loops, loops with E-E anchors as enhancer-enhancer (EE) loops, and all remaining combinations as ambiguous. For PE loops, we used KA-induced enhancers to define persistent activity-gained (PG), transient activity-gained (TG), persistent activity-strengthened (PS), and invariant-KA-induced loops; KA-decommissioned enhancers to define persistent activity-lost (PL), transient activity-lost (TL), persistent activity-weakened (PW), and invariant-KA-decommissioned loops; and KA-invariant enhancers to define invariant-KA-invariant loops.

For invariant-KA-induced, PG, PS, or TG loops, loop-anchored genes (*i.e.*, TUs) were removed if any of the following conditions were met: the P-anchor was associated with more than three promoters; the number of KA-decommissioned enhancers overlapping the enhancer anchor equaled or exceeded the number of KA-induced enhancers; or the numbers of KA-induced and KA-decommissioned enhancers were equal when the P-anchor was associated with more than three promoters.

For invariant-KA-decommissioned, PL, TL, or PW loops, loop-anchored genes (*i.e.*, TUs) were removed if any of the following conditions were met: the P-anchor was associated with three or more promoters; the number of KA-induced enhancers overlapping the enhancer anchor equaled or exceeded the number of KA-decommissioned enhancers; or the numbers of KA-decommissioned and KA-induced enhancers were equal when the P-anchor was associated with more than three promoters.

#### ***Gene ontology***

We extracted genes from PE and PP loops and performed gene ontology using WEB-based GEne SeT AnaLysis Toolkit (https://www.webgestalt.org) (**Table S11-S12**). Briefly, we transformed the upstream and downstream loop anchors into a bed file format and used "bedtools intersect" to obtain the gene names if there is an overlap with promoter region (-600 bp upstream and +200 bp downstream to TSS). Then, we performed gene ontology with the following parameters: Method of Interest as Over-Representation Analysis; Organism of Interest as Mus musculus; Functional Database as geneontology (Biological Process noRedundant); Select Reference Set as genome protein-coding; Significance Level as Top 10; rest as default **(Fig. S17)**.

#### ***Differentially methylation loci (DML) calling***

For DMR calling, we used the subtype-specific populations previously described in the *Contact maps generation and loop calling* section. We called DMLs using Dispersion Shrinkage for

Sequencing Data (DSS) (v2.54.0) (*83, 111*). Briefly, we merged the individual ALLC files from into pseudobulk ALLC files per neuron subtype. Then, we extracted the rows corresponding only to the cytosines (Cs) in the CpG context. To ensure high-quality genomic analyses, we excluded genomic blacklist regions, sex chromosomes, and mitochondrial chromosomes. Additionally, we filtered out CpGs with fewer than 3 coverage reads and removed cytosines not consistently present across all samples **(Fig. S22-S25)**. We performed statistical tests for differential methylation at each CpG site using the "DMLtest" method with smoothing enabled (smoothing.span = 50), comparing CFC3d GFP+ to CFC3d GFP- pseudobulk samples and CFC28d GFP+ to CFC28d GFP- pseudobulk samples. DMLs were identified by applying stringent thresholds of an absolute methylation fraction difference >= 0.2 between GFP+ and GFP- conditions and a significance cutoff of Pvalue < 0.001. We classified DMLs into four categories based on their temporal methylation dynamics. persistent activity-gained DMLs showed increased methylation fraction (>= 0.2, Pvalue < 0.001) consistently at both CFC3d and CFC28d, along with a sustained methylation fraction increase (>= 0.2) between CFC28d GFP+ and HC28d GFP+ conditions. Persistent activity-lost DMLs showed decreased methylation fraction (<= -0.2, Pvalue < 0.001) at both CFC3d and CFC28d and maintained reduced methylation fraction (<= -0.2) between CFC28d GFP+ and HC28d GFP+ conditions. Transient activity-gained DMLs showed increased methylation fraction (>= 0.2, Pvalue < 0.001) exclusively at CFC3d, with methylation fraction differences diminishing to <0.2 at CFC28d and changes within ±0.2 range relative to HC28d GFP+. Transient activity-lost DMLs showed decreased methylation fraction (<= -0.2, Pvalue < 0.001) exclusively at CFC3d, with methylation fraction differences reduced to > -0.2 at CFC28d time point and minimal differences changes within ±0.2 range compared to HC28d GFP+ **(Fig. S26, Table S13)**.

***Smart-seq3 seq data analysis***

We aligned the Smart-seq3 libraries as previously described (*104, 112*), using the zUMI pipeline (v2.9.7e) integrated with the STAR aligner **(Table S4)** (v2.7.10b). We installed the pipeline in a conda environment by following the instructions provided at https://github.com/sdparekh/zUMIs/wiki/Installation. The key packages along with their version numbers are given in the **Table S16**. To enable reproducibility, we also provide the specification file for the environment in our Bitbucket repository, which can be used to create a replica of our environment using the command "conda create" with the parameters -n as the environment and -file as the requirements file **(Table S16)**. Then, we installed (or re-installed some packages for version compatibility) the R packages in this environment **(Table S16)**. Next we modified yaml file based on the example file from (https://github.com/sdparekh/zUMIs/blob/main/ExampleData/runExample.yaml), which specifies the run parameters for the zUMI pipeline. We generated the STAR indexes and ran the zUMI pipeline for the mm10 genome using the following command "STAR" with the following parameters "--runThreadN" as 4, "--runMode" genomeGenerate, "--genomeFastaFiles" as the mm10 fasta file and "-y" as the path to yaml file. The zUMI output folder contains the aligned bam files along with the output counts matrix.

This mapping pipeline generated expression profiles capturing both UMI-containing 5' ends and internal reads from raw non-demultiplexed FASTQ files. For UMI detection in zUMIs, we specified "find_pattern": ATTGCGCAATG for Read 1 file. The base_definition in the YAML file was configured as follows: 150 bp paired-end mode, 573 cells total: including HC28d GFP+, HC28d GFP-, CFC3d GFP+, CFC3d GFP-, CFC3d RC GFP+, and CFC3d RC GFP- populations): cDNA 23-159 bp paired-end reads and UMI positions 12-19. HC28d GFP+ cells were excluded from analysis due to low cell numbers and sequencing depth. UMI deduplication employed a

Hamming distance threshold of 1. We aligned the reads to the mm10 reference genome using STAR with optimized parameters (--limitSjdbInsertNsj 2000000 --outFilterIntronMotifs RemoveNoncanonicalUnannotated --clip3pAdapterMMp 0.1 0.1 --clip3pAdapterSeq CTGTCTCTTATACACATCT CTGTCTCTTATACACATCT). Gene quantification was performed using GENCODE Release M23 (GRCm38.p6) annotation.

To generate transcript-level count matrices, we analyzed the Aligned.toTranscriptome.out.bam files generated from STAR aligner and zUMI pipeline output. We first demultiplexed these BAM files into individual single-cell BAM files using whitelist cell barcode information using Hamming distance matching with a maximum distance of 1 in cell barcodes. Each single-cell BAM file underwent transcript quantification using Salmon (v1.10.3) with the "quant" option, followed by merging all single-cell quantification matrices into a unified matrix. Since we have previously shown that the coordinates of chromatin loops correlate better with transcriptional units than genes (*51*), we didn't use the counts matrix generated by zUMI. Using a custom script, we merged different transcripts into transcription units (TUs). We merged transcripts of genes with transcription start sites (TSS) located within 10 kb of each other into transcriptional units (TUs). For each TU, the most upstream TSS was designated as its TSS, and the most downstream transcription end site (TES) was assigned as its TES. We filtered out cells containing fewer than 1000 detected TUs, as well as cells with fewer than 10,000 total counts **(Fig. S29)**. Additionally, we excluded cells with mitochondrial gene expression exceeding 1% of total counts, as elevated mitochondrial RNA can indicate compromised cell integrity or partial cytoplasmic contamination. We performed custom cell-wise scaling by dividing each cell's UMI values by that cell's mean UMI expression, effectively scaling each cell to have similar overall expression magnitudes. Furthermore, we filtered out the TUs that had >95% zeros in at least 2 cell conditions. We calculated the mean sum of counts per cell for each of four experimental conditions, then applied condition-specific scaling factors (normalized to CFC3d RC GFP+ as reference) to the expression data to correct for technical differences between conditions and stores the normalized data.

To clustering cells, we identified 3500 highly variable TUs using the FindVariableFeatures function and performed principal component analysis on the normalized data matrix using RunPCA of Seurat package (v5.1) (*61*). We employed 12 principal components with parameters min.dist = 0.1 and n.neighbors = 30 to calculate UMAP embeddings using RunUMAP and constructed a shared nearest neighbor graph using FindNeighbors, followed by cell clustering using the Louvain algorithm at a resolution of 2.8. After removing two clusters that showed mixed marker expression from different subtypes and was scattered across the UMAP embedding, we then categorized the resulting clusters into three major neuronal classes based on the expression patterns of key marker genes: excitatory neurons (*Slc17a7*), inhibitory neurons (*Gad2*), and DG neurons (*Prox1*) **(Fig. S29-30, Table S4)**. Excitatory neurons were further subdivided into five subtypes according to markers from the Allen Brain Atlas dataset (*41*). This classification aligned with the neuronal populations previously identified in snm3C-seq3 data.

To identify IEG+ cells, we implemented a binary classification approach based on the expression of three classical immediate early genes (*Arc*, *Egr1*, and *Nr4a1*) detected in our dataset. Cells were classified as IEG- if they expressed none of these genes or as IEG+ if they expressed at least one gene, using a threshold of any detectable expression (UMI count > 0).

We retained only transcription units whose promoters overlapped with loop anchors. For genes with multiple TUs in M-A plot, we applied a directional filtering approach to retain only biologically relevant transcripts based on the expected expression pattern of each gene list. Single-TU genes were retained without filtering. For multi-TU genes, the filtering logic operated as follows: For gene lists with positive expected directions, we retained all TUs showing upregulation

($log_2$ fold-change > 0) when at least one TU was upregulated. When all TUs showed downregulation, we retained only the TU with the highest expression (closest to zero). Conversely, for gene lists with negative expected directions, we retained all downregulated TUs ($log_2$ fold-change < 0) when downregulation was present, or the most downregulated TU when all showed upregulation. For unlooped genes without specific directional expectations, all TUs were retained regardless of expression direction. Log2 expression was calculated as fold change of the 75th percentile expression values between two conditions used.

***Odds Ratio and Fisher's exact test for enrichment of genes in double-stimulated over unstimulated control cells***

We use the genes upregulated or unchanged in TRAPed+, No Recall vs. TRAPed-, No Recall for each subtype TG P-E, PG P-E loop-anchored or unlooped genes. We used genes upregulated in TRAPed+, No Recall, then upregulated again in double-stimulated to test if they are enriched in PG or TG loops, we constructed a 2x2 contingency table with the following elements: a as the number of genes from PG or TG gene list with positive log2 expression (> 0), b as the number of genes from PG or TG gene with non-positive differential expression (<= 0), c as the number of unlooped genes with positive differential expression (> 0), and d as the number of unlooped genes with non-positive differential expression (<= 0). Log2 expression was calculated as fold change of the 75th percentile expression values between double-stimulated (TRAPed+, Recall IEG+) vs. unstimulated (TRAPed-, Recall IEG-) cells.

We used genes upregulated in TRAPed+, No Recall, then upregulated again in double-stimulated to test if they are enriched in PG or TG loops, we constructed a 2x2 contingency table with the following elements: a as the number of genes from PG or TG gene list with zero log2 expression ( = 0), b as the number of genes from PG or TG gene with non-zero differential expression (< 0 or > 0), c as the number of unlooped genes with zero differential expression (> 0), and d as the number of unlooped genes with non-zero differential expression (< 0 or > 0). Log2 expression was calculated as fold change of the 75th percentile expression values between double-stimulated (TRAPed+, Recall IEG+) vs. unstimulated (TRAPed-, Recall IEG-) cells.

We used genes unchanged in TRAPed+, No Recall, then upregulated in double-stimulated to test if they are enriched in PG or TG loops, we constructed a 2x2 contingency table with the following elements: a as the number of genes from PG or TG gene list with positive log2 expression (> 0), b as the number of genes from PG or TG gene with non-positive differential expression (<= 0), c as the number of unlooped genes with positive differential expression (> 0), and d as the number of unlooped genes with non-positive differential expression (<= 0). Log2 expression was calculated as fold change of the 75th percentile expression values between double-stimulated (TRAPed+, Recall IEG+) vs. unstimulated (TRAPed-, Recall IEG-) cells.

For each test, we calculated odds ratios and performed Fisher's exact test using the python function 'fisher_exact' from the package 'scipy.stats' and calculated the confidence intervals using the python functions 'Table2x2' and 'oddsratio_confint' from the package 'statsmodels.api'. Chi-square tests were performed using 'chi2_contingency' from 'scipy.stats' as an additional statistical validation.

***Odds Ratio and Fisher's exact test for neurological disorder risk loci***

We downloaded risk-loci associated with Autism Spectrum Disorder (ASD) from Simons Searchlight (https://gene.sfari.org/database/human-gene/), and risk-loci associated with Post-Traumatic Stress Disorder (PTSD) from Nievergelt et al. (*70*). We constructed a 2x2 contingency table with the following elements: a as the number of loop anchored genes overlapping with ASD/PTSD, b as the number of loop anchored genes not-overlapping with ASD/PTSD, c as the number of unlooped genes overlapping with ASD/PTSD, and d as the number of unlooped genes not-overlapping with ASD/PTSD. Statistical approach was same as described above for stimulated gene enrichment.
